# An Injectable Subcutaneous Colon-Specific Immune Niche For The Treatment Of Ulcerative Colitis

**DOI:** 10.1101/2023.10.03.560652

**Authors:** Kin Man Au, Justin E. Wilson, Jenny P.-Y. Ting, Andrew Z. Wang

## Abstract

As a chronic autoinflammatory condition, ulcerative colitis is often managed via systemic immunosuppressants. Here we show, in three mouse models of established ulcerative colitis, that a subcutaneously injected colon-specific immunosuppressive niche consisting of colon epithelial cells, decellularized colon extracellular matrix, and nanofibers functionalized with programmed death-ligand 1, CD86, a peptide mimic of transforming growth-factor-beta 1, and the immunosuppressive small molecule leflunomide, induced intestinal immunotolerance and reduced inflammation in the animals’ lower gastrointestinal tract. The bioengineered colon-specific niche triggered autoreactive-T-cell anergy and polarized pro-inflammatory macrophages via multiple immunosuppressive pathways, and prevented the infiltration of immune cells into the colon’s lamina propria, promoting the recovery of epithelial damage. The bioengineered niche also prevented colitis-associated colorectal cancer, and eliminated immune-related colitis triggered by kinase inhibitors and immune-checkpoint blockade.

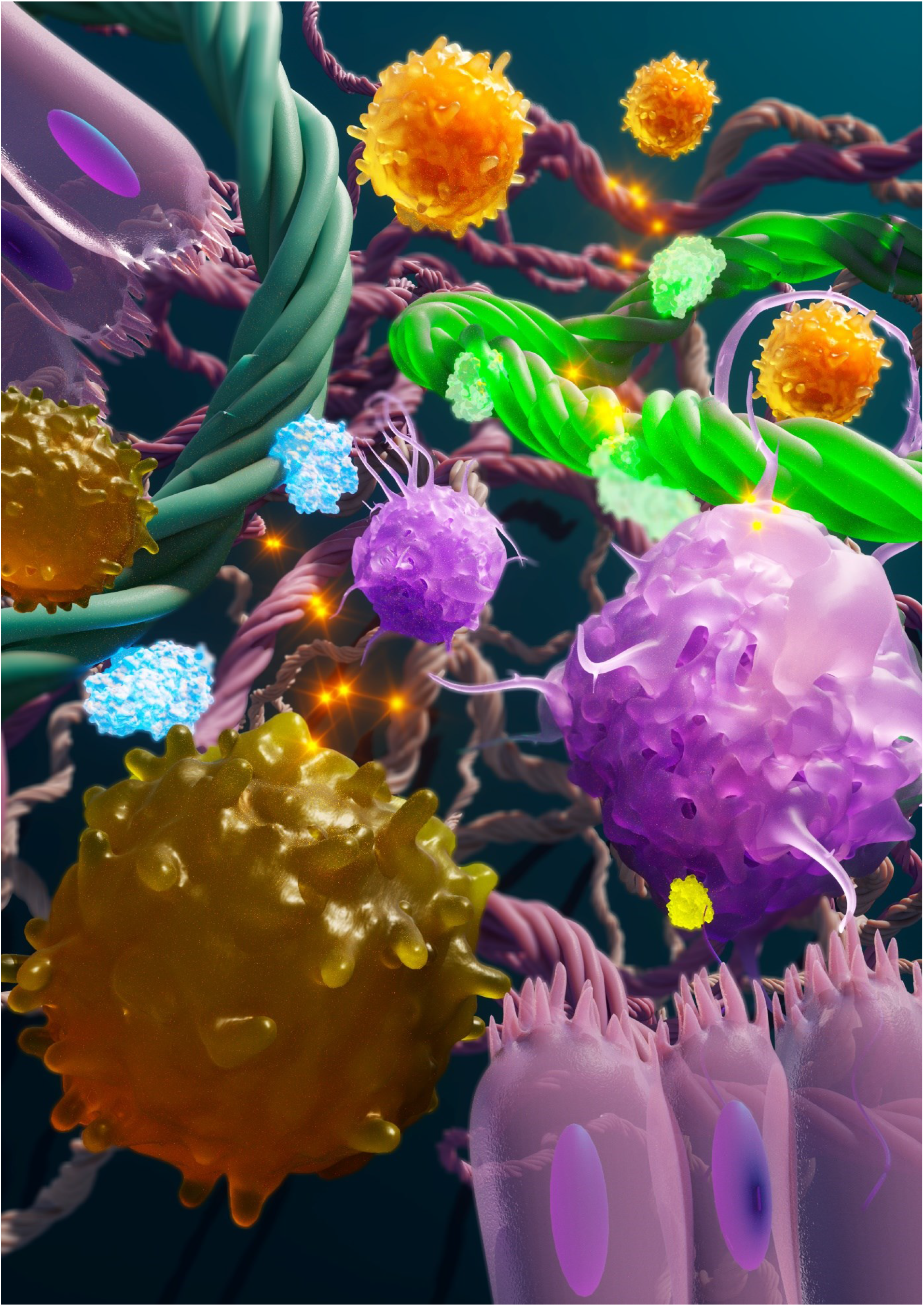

## One-sentence editorial summary

A subcutaneous colon-specific niche consisting of colon epithelial cells, decellularized colon extracellular matrix and immunosuppressive nanofibers reduced inflammation in the gastrointestinal tract of mice with established ulcerative colitis.

Ulcerative colitis is a chronic autoinflammatory bowel disease^1-3^ in which autoreactive T cells^4^ and macrophages^5, 6^ attack healthy colon epithelial cells in the colonic mucosa, leading to chronic inflammation and ulceration^7, 8^. Ulcerative colitis symptoms can be debilitating and can lead to life-threatening complications^7^, such as bowel perforation and toxic megacolon. While the causes of ulcerative colitis are not fully understood, it is widely believed to be triggered by genetic mutations^1-3, 9^, immune system malfunction^1-3, 10^, and environmental factors^1-3, 9^. Despite intense treatment development efforts, ulcerative colitis remains incurable. Current treatment strategies generally involve immunosuppressive therapies and surgical resection^11, 12^. However, long-term immune suppression has significant side effects, including infection^13^ and cancer development^14, 15^, and surgery has high morbidity and long-term health complications. Therefore, novel and curative treatment development for ulcerative colitis is urgently needed.

Antigen-specific immunotherapy^16, 17^, which establishes disease and antigen-specific immunotolerance, is a new immunosuppressive approach for autoimmune diseases. Several studies have shown that antigen-specific immunotolerance can be achieved by presenting disease-specific antigens to T cells in the presence of immunomodulators^17-19^, such as immunosuppressive immune checkpoint molecules and pro-regulatory cytokines. However, this approach is ineffective for autoinflammatory diseases such as ulcerative colitis that do not have common identifiable autoantigens. One potential solution is to present the entire cell targeted by autoimmunity to autoreactive T cells in the presence of immunomodulators in an organ-like tissue microenvironment. Here, we report the bioengineering of an injectable colon-specific immune niche containing an immune suppressive microenvironment to treat ulcerative colitis (Fig. 1a). We hypothesized that this niche would interact with autoreactive T cells in the immunogenic subcutaneous (s.c.) environment^20, 21^ involved in ulcerative colitis pathogenesis, exhausting and suppressing autoreactive T cells and polarizing proinflammatory classically-activated M1 macrophages. Moreover, the engineered niche would reduce autoimmunity and induce tolerance, resolving ulcerative colitis symptoms (Fig 1a). Consequently, it would restore a homeostatic tissue environment to allow the intestinal stem cells in the crypts to repair the damaged colon epithelium by replacing the dying colon epithelial cells^22, 23^. To examine this anti-colitis effect, we used three different established ulcerative colitis mouse models.

**Fig. 1.**
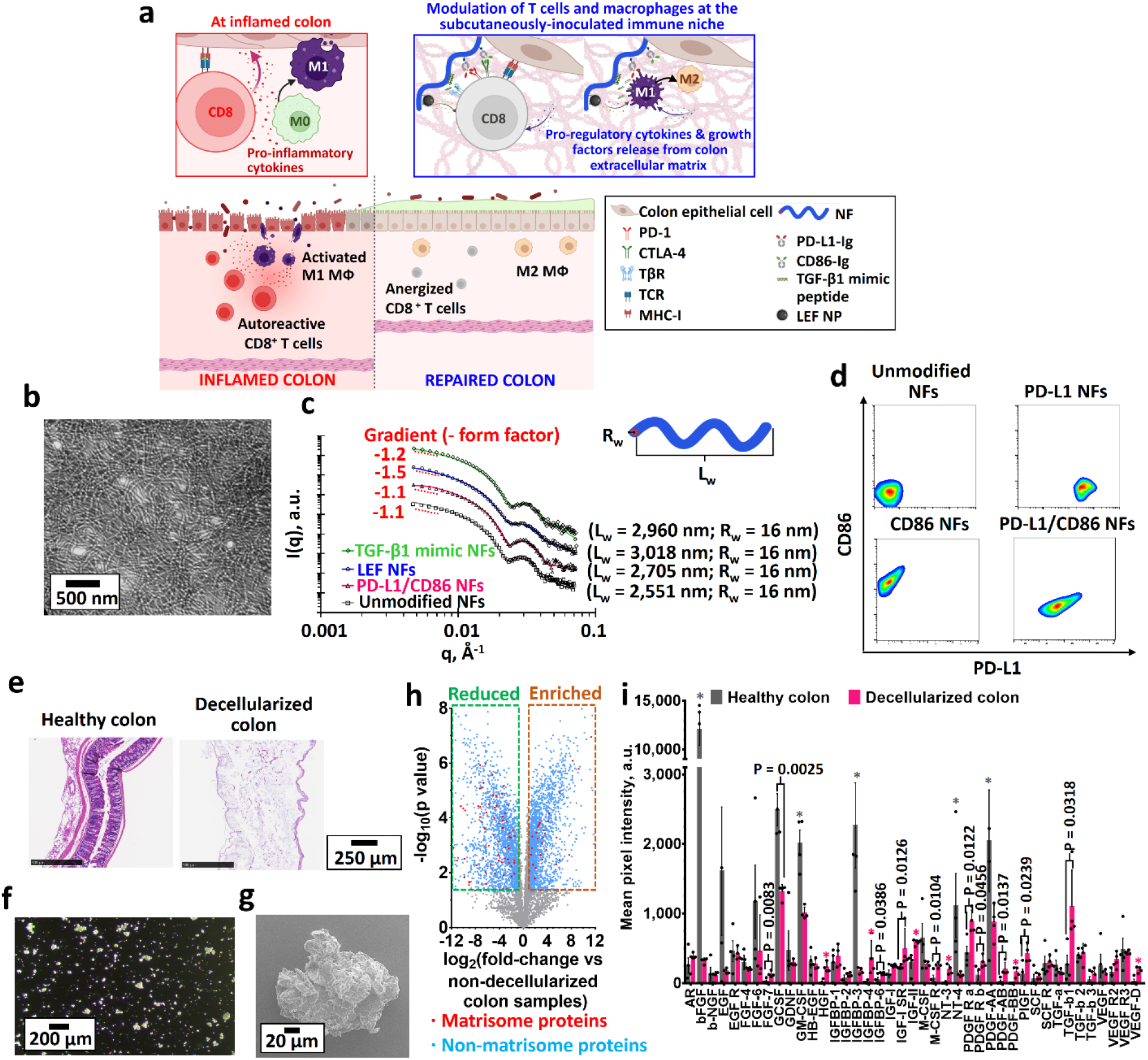
Bioengineering and characterization of the immunosuppressive combinational colon-specific immune niche. **a**, Schematic of the proposed combinational colon-specific immune niche mechanism of action. The niche anergize autoreactive CD8^+^ T cells and polarizes classically-activated M1 macrophages (M1 MФs) to alternatively-activated M2 macrophages (M2 MФs). The niche promotes the recovery of ulcerated colon epithelium by producing a non-inflammatory environment for colon stem cells at crypts to replace apoptotic colon epithelial cells. Graphic created with BioRender.com. MФ, macrophage; TβR, TGF beta receptor; TCR, T cell receptor; MHC, major histocompatibility complexes; PD-L1-Ig, PD-L1-immunoglobulin fusion protein; CD86-Ig, CD86 immunoglobulin fusion protein; LEF NP, leflunomide-encapsulated nanoparticles; NF, nanofiber. **b**, Representative transmission electron microscopy image of unmodified carboxylic acid-functionalized poly(2-(methacryloyloxy)ethyl phosphorylcholine)- poly(2-hydroxypropyl methacrylate) nanofibers. **c**, Background-subtracted solution small-angle X-ray scattering profiles of the unmodified and functionalized immunosuppressive nanofibers. The open dots represent the collected angle-dependent scattering intensities, and the solid lines represent the best fit to the cylinder scattering model. Lw, weight-averaged length; Rw, weight-averaged radius; PD-L1/CD86 NFs, PD-L1/CD86 dual-functionalized nanofibers; TGF-β1 mimic NFs, TGF-β1 mimic peptide-encapsulated nanofibers; LEF NFs, leflunomide-encapsulated nanofibers. **d**, *In vitro* binding of phycoerythrin (PE)-labeled anti-PD-L1 and A488-labeled anti-CD86 to PD-L1 and/or CD86 mono/dual-functionalized nanofibers NFs, as quantified *via* fluorescence-activated cell sorting. **e**, Representative H&E-stained images of the mouse colon before and after decellularization. **f**, Representative dark-field optical microscopy image of the ball-milled decellularized colon extracellular matrix (ECM). **g**, Representative scanning electron microscope image of the ball-milled decellularized colon extracellular matrix. **h**, Volcano plots showing proteomic data for a naïve mouse colon and a decellularized colon extracellular matrix. (n = 3 biologically independent samples) **i,** Analysis of growth factors and cytokines present in mouse colon before and after decellularization *via* sandwich-type antibody microarray assay. bFGF, basic fibroblast growth factor; EGF, epidermal growth factor; EGFR, epidermal growth factor receptor; FGF-4, fibroblast growth factor-4; FGF-6, fibroblast growth factor-6; FGF-7, fibroblast growth factor-7; G-CSF, granulocyte-colony stimulating factor; GDNF, glial-derived neurotrophic factor; GM-CSF, granulocyte macrophage-colony stimulating factor; HB-EGF, heparin-binding epidermal growth factor; IGFBP-1, insulin-like growth factor binding protein 1; IGFBP-3, insulin-like growth factor binding protein 3; IGFBP-4, insulin-like growth factor binding protein 4; IGFBP-6, insulin-like growth factor binding protein 6; IGF-I, insulin-like growth factor-I; IGF-1 SR, insulin-like growth factor-I receptor; IGF-II, insulin-like growth factor-II; M-CSF, macrophage-colony stimulating factor; M-CSF R, macrophage-colony stimulating factor receptor; NT-3, neurotrophin-3; NT-4, neurotrophin-4; PDGF-R α, platelet-derived growth factor receptor alpha; PDGF-R β, platelet-derived growth factor receptor beta; PDGF-AA, platelet-derived growth factor AA; PDGF-AB, platelet-derived growth factor AB; PDGF-BB, platelet-derived growth factor BB; PIGF, phosphatidylinositol glycan anchor biosynthesis class F; SCF, stromal cell-derived factor-1; SCF R, stromal cell-derived factor receptor; TGF-α, transforming growth factor alpha; TGF-β, transforming growth factor beta; TGF-β 2, transforming growth factor beta 2, TGF- β 3, transforming growth factor beta 3; VEGF, vascular endothelial growth factor; VEGF R2, vascular endothelial growth factor receptor 2; VEGF R3, vascular endothelial growth factor receptor 3; VEGF-D, vascular endothelial growth factor receptor D. (n = 4 biologically independent samples) Data are presented as the mean ± standard error of the mean (s.e.m.). All P values were analyzed using two-way ANOVA with Tukey’s HSD multiple comparisons post-hoc test. * denotes p < 0.001, hence statistically significant.

## Results

### Bioengineering of organ-specific immune niches

To create a colon-like tissue microenvironment for modulating autoreactive immune cells, we bioengineered colon-specific immune niches from colon epithelial cells and decellularized colon extracellular matrix. The niche was further engineered with nanofibers containing immunosuppressive molecules. The colon epithelial cells and decellularized colon extracellular matrix recapitulate a colon-like microenvironment to attract and interact with autoreactive immune cells involved in ulcerative colitis. The decellularized colon extracellular matrix also acts as a depot that gradually releases pro-regulatory growth factors and cytokines to modulate the activation of autoreactive T cells and macrophages. Immunosuppressive molecule-functionalized nanofibers were engineered to simultaneously anergize autoreactive T cells and polarize macrophages through multiple immunosuppressive signaling pathways by incorporating immune checkpoint molecules and other immunosuppressants. These included programmed death-ligand 1 (PD-L1), which triggers the suppressive PD-1/PD-L1 immune checkpoint pathway^24-26^ and induces the polarization of classically-activated M1 macrophages into alternatively-activated M2 macrophages^27^; cluster of differentiation 86 (CD86), which triggers the immunosuppressive cytotoxic T-lymphocyte-associated protein 4 (CTLA-4) pathway in activated CD8^+^ T cells^25, 26, 28^; immunomodulatory drug leflunomide which inhibits activated T cell and macrophage proliferation^29, 30^; and a pro-regulatory cytokine transforming growth factor-beta 1 (TGF-β1) mimetic peptide, which binds to the transforming growth factor-beta receptor II receptor to suppress the activation of CD8^+^ T cells^31, 32^ and polarize classically-activated M1 macrophages to alternatively-activated M2 macrophages^33, 34^.

Modular-based immunosuppressive molecule-functionalized nanofibers were engineered using poly(2-(methacryloyloxy)ethyl phosphorylcholine)-poly(2-hydroxypropyl methacrylate) diblock copolymer as a scaffold. The diblock copolymer was synthesized through a two-step reversible addition-fragmentation chain transfer polymerization^35-37^, and self-assembled into rod-shaped, fiber-like micelles *in situ* (Fig. 1b, Supplementary Fig. 1-6 and Supplementary Tables 1-2). The carboxylic acid terminus of the chain transfer reagent’s alpha-end group on the nanofiber surface was converted into a dibenzocyclooctyne group *via* an N-hydroxysulfosuccinimide/1- ethyl-3-(3-dimethylaminopropyl)carbodiimide coupling reaction with the sulfo-dibenzocyclooctyne amine (Supplementary Fig. 2). PD-L1 and CD86 mono- and dual-functionalized nanofibers and leflunomide encapsulated nanofibers were prepared from dibenzocyclooctyne functionalized nanofibers by strain-promoted azide-alkyne cycloaddition with the corresponding azide-functionalized immune checkpoint ligands (Supplementary Fig. 7-9) and azide-functionalized leflunomide-encapsulated poly(ethylene glycol)-poly(lactide-co-glycolide) nanoparticles^25, 26^ (Supplementary Fig. 10). TGF-β1 mimetic peptide-encapsulated nanofibers engineered through the ionic adsorption of the oligomerized cationic TGF-β1 mimetic peptide onto anionic carboxylic acid-functionalized nanofibers (Supplementary Fig. 11). These non-thermal responsive nanofibers (Supplementary Fig. 6) can be manufactured on industrial scale and stored at 4 °C before further use.

Transmission electron microscopy confirmed that the *in situ* self-assembled diblock copolymer nanofibers retained their fiber-like morphology after functionalization (Fig. 1b and Supplementary Fig. 9-11). Local nanofiber aggregation was observed in the leflunomide-encapsulated nanofibers and TGF-β1 mimic-encapsulated nanofibers since both functionalization components can act as bridges to cross-link the nanofibers. Furthermore, solution small-angle X-ray scattering analysis^38^ (Fig. 1c and Supplementary Fig. 12) showed that all functionalized nanofibers conformed to a core-shell cylinder model with gradients of -1.1 (carboxylic acid-functionalized nanofibers) to -1.5 (leflunomide-encapsulated nanofibers) in the Guinier region. Steeper gradients were recorded for the leflunomide-encapsulated nanofibers and transforming growth factor-beta 1 mimic-encapsulated nanofibers due to their partial aggregation. Nevertheless, the weight-averaged lengths of all nanofibers were approximately 3 µm, and their core weight-averaged radius was 16 nm. *In vitro* binding studies using the fluorescence-activated cell sorting method confirmed that both PD-L1 and CD86 mono-/dual-functionalized nanofibers were bound selectively to their corresponding fluorescently labeled antibodies (Fig. 1d). A drug release study confirmed the controlled release of the encapsulated leflunomide and adsorbed TGF-β1 mimetic peptide from the nanofibers under physiological conditions (Supplementary Fig. 10-11). All functionalized NFs showed minimal *in vitro* toxicity to cultured colon epithelial cells (Supplementary Fig. 13).

To engineer the colon-specific immune niche, we first generated decellularized colons from preserved mouse colons using a spin-decellularization protocol^39, 40^. Histological analysis confirmed the absence of cells and the presence of collagen in the decellularized colon scaffolds (Fig. 1e), and the DNA content of the colons was reduced by 92% on average (Supplement Fig. 14). Ball milling of the lyophilized decellularized colon scaffolds yielded sub-hundred micrometer-sized subcutaneously injectable fine powders (Fig. 1f,g). Proteomic analysis confirmed that decellularization preserved key nonmatrisome and matrisome proteins (Fig. 1h and Supplementary Table 3). Additionally, the multiplexed growth factor immunoassay showed that the decellularized colon extracellular matrix preserved most of the immunomodulatory growth factors (e.g., granulocyte colony-stimulating factor (GCSF)^41, 42^, granulocyte-macrophage colony-stimulating factor (GM-CSF)^42, 43^, and platelet-derived growth factor (PDGF)^42^) and pro-regulatory cytokines (e.g., TGF-β1^31-34^) (Fig. 1i). An *in vitro* proliferation studies were used to identify the optimal concentration of colon extracellular matrix (500 µg of colon extracellular matrix/10^6^ colon epithelial cells) colon epithelial cell growth in a reduced-serum colon epithelial cell culture medium since high growth factor and cytokine concentrations suppress colon epithelial cell growth (Supplementary Fig. 15). Although this study focuses on the use of mouse colon epithelial cells and colon extracellular matrix to engineer the immune niche, human colon epithelial cells and colon extracellular matrix can be isolated *via* similar methods. More specifically, human colon epithelial cells can be isolated from intestinal crypts collected through endoscopic epithelial biopsy^44^, and human colon extracellular matrix can be isolated from preserved donor colon (from colonic disease-free deceased donor) followed by detergent-facilitated decellularization and sterilization^39, 40^. In recent years, protocols have been established to sterilize tissue allografts to prevent disease transmission from the donor^45^. In fact, a recent study demonstrated the potential of using organ-specific extracellular matrix to repair inflamed tissues^46^. Furthermore, the decellularized colon extracellular matrix can be replaced by fully synthetic artificial extracellular matrix in the near future, like the artificial spinal cord extracellular matrix for the maturation of human stem cell-derived neurons^47^.

### Immunosuppressive molecule-functionalized nanofibers effectively inhibit autoreactive CD8^+^ T cell activation and induce classically-activated M1 macrophage polarization *in vitro*

We next evaluated the immunosuppressive effects of nanofibers formulated with different immunosuppressive molecules and colon extracellular matrix *in vitro*. We first evaluated their abilities to modulate autoreactive CD8^+^ T cells *in vitro*. The NFs were co-cultured with CD8^+^ T cells with anti-CD3/anti-CD28-functionalized microparticles (Dynabeads^TM^) and interleukin-2 (IL-2)^48^. Both unconjugated (“free”) and nanofiber-conjugated PD-L1 and CD86 effectively upregulated T cell exhaustion marker expression (e.g., LAG-3 and TIM-3)^49^ in stimulated CD8^+^ T cells. However, the PD-L1-functionalized nanofibers showed a higher inhibitory efficiency than free PD-L1 and CD86 at a 1:1 ratio (Extended Data Fig. 1 and Supplementary Fig. 17). The PD- L1/CD86 dual-functionalized nanofibers were more effectively upregulated the inhibitory markers than the free immune checkpoint ligands. As in previous studies, both forms of the TGF-β1 mimetic peptide upregulated inhibitory markers on the stimulated CD8^+^ T cells^31, 32^, with further upregulation observed after coculturing with small-molecule leflunomide or leflunomide-encapsulated nanofibers (Extended Data Fig. 1 and Supplementary Fig. 16). Furthermore, culturing the activated CD8^+^ T cells with the combination of PD-L1, CD86, leflunomide, and TGF- β1 mimetic peptide, especially the combination of all three different immunosuppressive molecule-functionalized nanofibers (combinational nanofibers), significantly upregulated LAG-3 and TIM-3 expression by 7- and 26- fold (Fig. 2a, Extended Data Fig. 1 and Supplementary Fig. 16-17), respectively. The optimized colon extracellular matrix concentration was as effective as the combinational nanofibers in anergizing stimulated CD8^+^ T cells and upregulating T cell exhaustion markers (Fig. 2a and Supplementary Fig. 17) due to the preserved immunomodulatory growth factors and cytokines. Additionally, the mixture of the combinational nanofibers and colon extracellular matrix cooperatively upregulated TIM-3 and LAG-3 expressions (Fig. 2a and Supplementary Fig. 17). Further *in vitro* cell proliferation studies indicated that all immunosuppressive nanofibers, especially the combinational nanofibers, and colon extracellular matrix effectively inhibited proliferation of carboxyfluorescein succinimidyl ester-labeled CD8^+^ T cells^50^ under the same activation conditions (Fig. 2b, Extended Data Figure 1 and Supplementary Fig. 18-19). The combination of combinational nanofibers and colon extracellular matrix effectively inhibited the stimulated CD8^+^ T cell proliferation, with less than 5% of the T cells proliferating (Fig. 2b and Supplementary Fig. 19). The combined inhibition effect was particularly significant when a low concentration of combinational nanofibers was combined with a low concentration of colon extracellular matrix in the T cell proliferation assay (Fig. 2b and Supplementary Fig. 19).

**Fig. 2.**
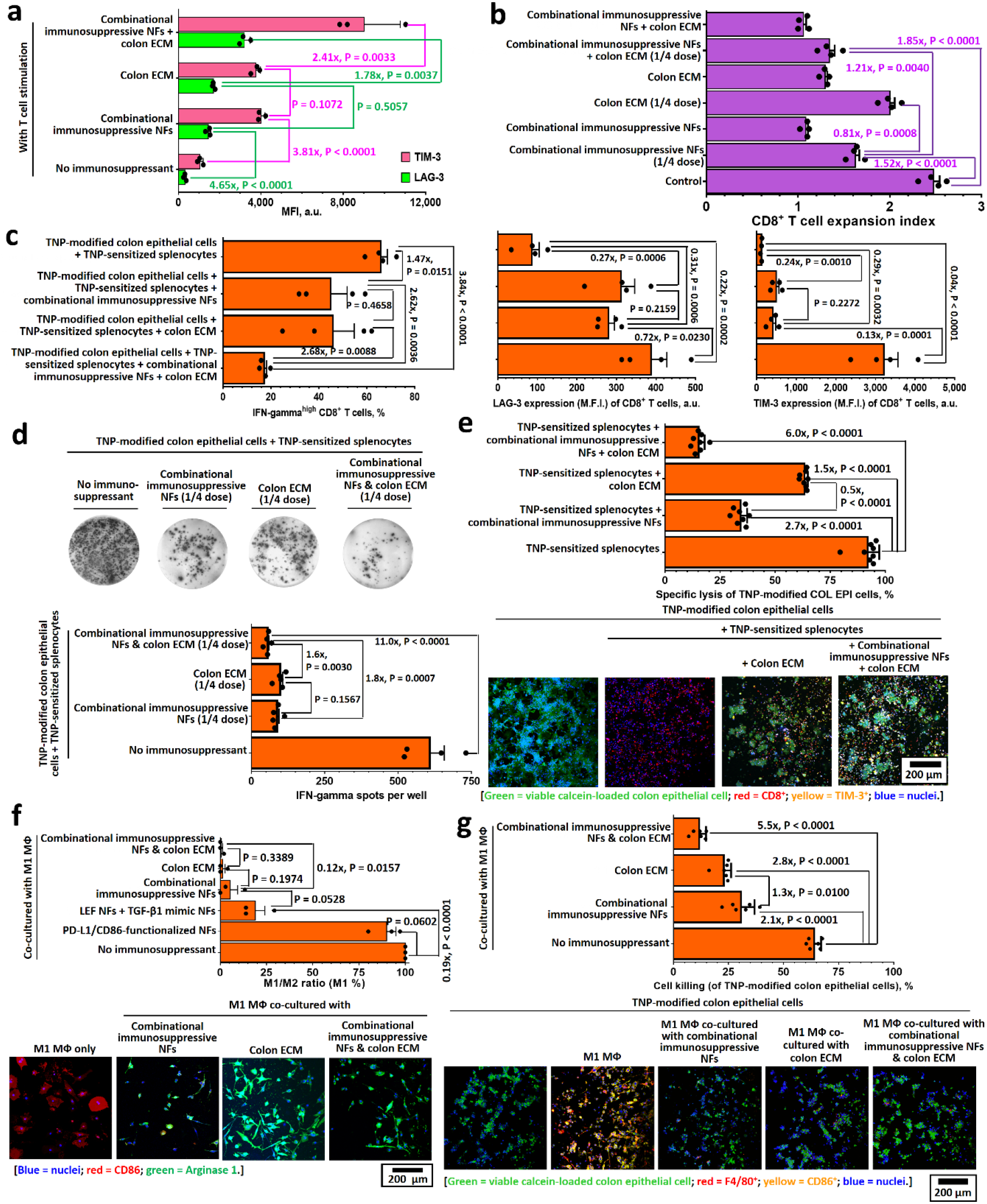
Combinational immunosuppressive nanofibers and colon extracellular matrix effectively inhibit antigen-specific CD8^+^ T cell activation and promote classically-activated M1 macrophages differentiation into alternatively-activated M2 macrophages *in vitro*. **a-b,** T cell inhibition (exhaustion) markers (**a**) and proliferation indices (**b**) of CD8^+^ T cells cultured with combinational nanofibers, colon extracellular matrix and their mixture in the presence of anti-CD3/anti-CD28-functionalized T cell activation beads (Dynabeads) at a 1:1 ratio for 48 h, as quantified through fluorescence-activated cell sorting. The proliferation study was performed on carboxyfluorescein succinimidyl ester-labeled CD8^+^ T cells. (n = 4). **c,** T cell activation and exhaustion markers expressed on trinitrophenol-sensitized CD8^+^ T cells (in splenocytes) after co-culturing with trinitrophenol-modified colon epithelial cells in the presence or absence of combinational nanofibers, colon extracellular matrix, and their mixture for 48 h. (n = 4). **d,** ELISpot assay of IFN-γ in splenocytes after culture with trinitrophenol-modified colon epithelial cells in the presence or absence of combinational nanofibers, colon extracellular matrix, and their mixture (n = 4). **e,** Cytotoxicity of trinitrophenol-sensitized splenocytes against trinitrophenol-modified colon epithelial cells in the presence or absence of combinational nanofibers, colon extracellular matrix, and their mixture (n = 7). **f,** Ratio of classically-activated M1 macrophages to alternatively-activated M2 macrophages after IFN-γ- and lipopolysaccharides-induced classically-activated M1 macrophages were co-cultured with combinational nanofibers, colon extracellular matrix, and their mixture, as determined by confocal laser scanning microscopy. Classically-activated M1 macrophages are characterized by roundish cell bodies with a few cytoplasmic extensions on the cell surface, and alternatively-activated M2 macrophages are characterized by its elongated cell body with cytoplasmic extensions on their apical ends. (n = 80 – 100 cells per group) **g,** Quantification of trinitrophenol-modified colon epithelial cells killed by induced classically-activated M1 macrophages in the presence of combinational nanofibers, colon extracellular matrix, and their combination. (n = 7) Data are presented as the mean ± s.e.m. All P values were analyzed using two-way ANOVA with Tukey’s HSD multiple comparisons post-hoc test.

To demonstrate the abilities for the combinational nanofibers and colon extracellular matrix to inhibit antigen-specific CD8^+^ T cell activation, we analyzed CD8^+^ T cell phenotypes and their cell-killing efficiency after co-culturing with trinitrophenol-modified colon epithelial cells^51, 52^. Immunogenic trinitrophenol-modified colon epithelial cells effectively activated CD8^+^ T cells in both trinitrophenol-sensitized and non-sensitized splenocytes (Fig. 2c, Extended Data Figure 1 and Supplementary Fig. 20). The sensitized CD8^+^ T cells showed significantly higher (over 2-fold) intercellular interferon-gamma (IFN-γ) and granzyme B expressions^52^ than the non-sensitized CD8^+^ T cells (Extended Data Fig. 1 and Supplementary Fig. 20). In contrast, no significant CD8^+^ T cell activation was observed when the trinitrophenol-sensitized splenocytes were cultured with the unmodified colon epithelial cells (Extended Data Figure 1 and Supplementary Fig. 20). The co-culturing of the sensitized splenocytes with the trinitrophenol-modified colon epithelial cells in the presence of the combinational nanofibers or colon extracellular matrix markedly reduced the expression of both intercellular T cell activation markers (Fig. 2c and Supplementary Fig. 21). A more profound inhibitory effect was observed when the sensitized splenocytes were cultured with the combinational nanofibers and colon extracellular matrix, where the intercellular IFN-γ expression in the sensitized CD8^+^ T cells was synergistically reduced by 74% (Fig. 2c and Supplementary Fig. 21), whereas combinational nanofibers or colon extracellular matrix alone only reduced the frequencies of IFN-γ^high^ CD8^+^ T cells by approximately 30%. Phenotypic analysis showed that co-culturing the trinitrophenol-modified colon epithelial cells with combinational nanofibers and colon extracellular matrix effectively upregulated inhibitory marker expression (LAG-3 and TIM-3) in the sensitized CD8^+^ T cells (Fig. 2c, Extended Data Figure 1 and Supplementary Fig. 22-23). Furthermore, the inhibition reduced pro-inflammatory cytokine (e.g., IFN-γ) secretion when the sensitized splenocytes were cocultured with the trinitrophenol-modified colon epithelial cells (Fig. 2d, Extended Data Figure 1 and Supplementary Fig. 24-25). Consequently, the specific lytic activity against the trinitrophenol-modified colon epithelial cells was reduced by 91% when the sensitized splenocytes were cocultured with the combinational nanofibers and colon extracellular matrix (both at one-quarter of the optimized doses) (Fig. 2e and Supplementary Fig. 25). Therefore, combining combinational nanofibers and colon extracellular matrix can effectively inhibit antigen-specific CD8^+^ T cell lysis.

The upregulation of colonic classically-activated M1 macrophages contributes to ulcerative colitis development^5, 6^. Proinflammatory cytokines (e.g., IFN-γ and tumor necrosis factor-alpha) produced by colonic CD8^+^ T cells induce the polarization of colonic unpolarized M0 macrophages^5, 6^. To demonstrate that combinational nanofibers and colon extracellular matrix can promote proinflammatory classically-activated M1 macrophages to differentiate into anti-inflammatory alternatively-activated M2 macrophages, we analyzed the phenotypes of IFN-γ- and lipopolysaccharide (LPS)-induced splenic classically-activated M1 macrophages after co-culturing with different immunosuppressive components. Similar to free TGF-β1^33, 34^, TGF-β1 mimic-encapsulated nanofibers effectively polarized approximately 74% of classically-activated M1 macrophages into alternatively-activated M2 macrophages (Fig. 2f and Supplementary Fig. 26). Although other mono-functionalized NFs were not adequate to induce classically-activated M1 macrophage polarization, approximately 94% of classically-activated M1 macrophage were differentiated into alternatively-activated M2 macrophages after co-culturing with combinational nanofibers (Fig. 2f and Supplementary Fig. 26). Due to the retention of pro-regulatory growth factors and chemokines after colon decellularization (Fig. 1i), the colon extracellular matrix effectively induced more than 98% of classically-activated M1 macrophages to differentiate into alternatively-activated M2 macrophages (Fig. 2f and Supplementary Fig. 26). Consequently, combining combinational nanofibers and colon ECM effectively converted all classically-activated M1 macrophages into alternatively-activated M2 macrophages (Fig. 2f and Supplementary Fig. 26).

To demonstrate that the polarization of classically-activated M1 macrophages reduces their cell-killing efficiency, we co-culture trinitrophenol-modified colon epithelial cells with IFN-γ- and LPS-polarized classically-activated M1 macrophages with different immunosuppressants. Due to the immunogenic nature of the trinitrophenol-modified colon epithelial cells, classically-activated M1 macrophages lysed approximately 65% of the colon epithelial cells (Fig. 2g). Co-culturing the colon epithelial cells with the combinational nanofibers inhibited cell killing by approximately half and reduced the CD86^+^ classically-activated M1 macrophages (Fig. 2g), consistent with the ability of the combinational nanofibers to polarize classically-activated M1 macrophages to alternatively-activated M2 macrophages. A similar inhibitory effect was observed when the colon epithelial cells were co-cultured with colon extracellular matrix. Co-culturing the colon epithelial cells with the combinational nanofibers and colon extracellular matrix significantly inhibited cell killing of classically-activated M1 macrophages by 5.5-fold with < 12% of colon epithelial cells being lysed by the classically-activated M1 macrophages (Fig. 2g). These findings show that combining the combinational nanofibers and colon extracellular matrix can additively inhibit non-specific cell killing by polarizing classically-activated M1 macrophages into alternatively-activated M2 macrophages.

### Colon-specific immune niches effectively ameliorate colitis in established ulcerative colitis mouse models

Based on the promising *in vitro* findings, we examined whether the subcutaneous inoculation of colon-specific immune niche could induce organ-specific immunotolerance, ameliorate acute ulcerative colitis symptoms and speed up the recovery of ulcerated colonic mucosa using a well-established dextran sodium sulfate-induced ulcerative colitis mouse model^53, 54^. We first evaluated the therapeutic efficacy of immune checkpoint molecule-functionalized colon-specific immune niches (Fig. 3a). We found that all colon-specific immune niche bioengineered with immune checkpoint molecules effectively suppressed colitis symptoms (e.g., diarrhea and rectal bleeding; Fig. 3b), minimized bodyweight loss (Extended Data Fig. 2a), and prevent inflammation-associated colon shortening in mice (Fig. 3c and Extended Data Fig. 2b). Specifically, at the study endpoint, mice treated with PD-L1/CD86 dual-functionalized niches exhibited complete bodyweight recovery (Extended Data Fig. 2a), reduced inflammation-associated myeloperoxidase activity (Extended Data Fig. 3a), attenuated local pro-inflammatory cytokine expression (e.g., interleukin-6 and TNF-α) (Extended Data Fig. 3b), and increased pro-regulatory cytokine expression (e.g., interleukin-10 and TGF-β1; Extended Data Fig. 3c). Histological analysis confirmed that all treatments with immune checkpoint molecule-functionalized niches significantly reduced immune cell infiltration into the colonic mucosa and submucosa (Fig. 3d,e and Extended Data Fig. 2c).

**Fig. 3.**
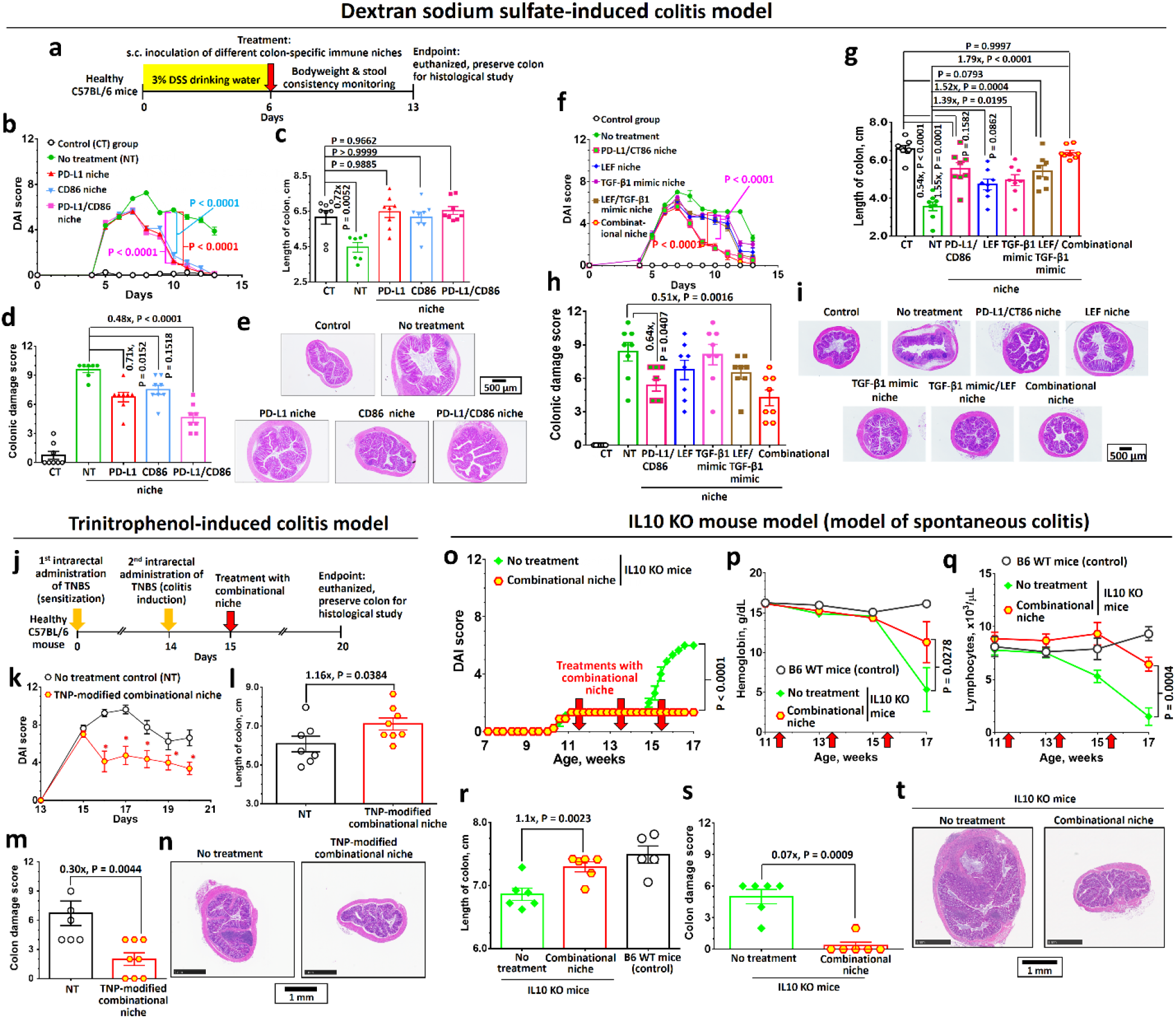
Combinational colon-specific immune niche effectively ameliorated colitis and promoted recovery in established mouse ulcerative colitis models. **a-i,** Optimization of Colon-specific immune niche formulations in the dextran sodium sulfate (DSS)-induced colitis model. Colitis treatment schedule (**a**): each colitis mouse received a single subcutaneous inoculation of colon-specific immune niche at day 6 (onset of colitis symptoms). Bodyweight changes and disease activity index (DAI) scores were monitored daily for 13 days. The colons were preserved at the study endpoint (day 13). Disease activity index score (DAI) (**b**) and length of colon preserved at the study endpoint (**c**) after colitis treatment with different immune checkpoint molecule-functionalized niches. (n = 8). Colon damage score (**d**) and representative H&E-stained colon sections (**e**) preserved from colitis mice after different treatments with immune checkpoint molecule-functionalized colon-specific immune niches. Disease activity index score (**f**) and length of colon preserved at the study endpoint (**g**) after colitis treatment with different non-immune checkpoint molecule-functionalized niches and combinational niche. (n = 8) Colon damage score (**h**) and representative H&E-stained colon sections (**i**) preserved from colitis mice after different treatments with immune checkpoint molecule-functionalized niches. **j-n,** Evaluation of the optimized combinational niche formulation for colitis treatment in a trinitrophenol-induced colitis model. Colitis induction and treatment schedule (**j**): each colitis mouse received a single subcutaneous inoculation of trinitrophenol-modified combinational niche (engineered with trinitrophenol-modified colon epithelial cells) at day 15 (onset of colitis symptoms). The colon was preserved at the endpoint (day 20) for mechanistic study. TNBS, trinitrobenzene sulfonic acid. Disease activity index score of trinitrophenol-induced colitis mice after colitis treatment with trinitrophenol-modified combinational colon-specific immune (**k**). Length of colon preserved at the experiment endpoint (**l**). Colon damage score (**m**) and representative H&E-stained colon sections (**n**) preserved from TNP-induced colitis mice after treatment with trinitrophenol-modified combinational niche. (n = 8, 1 mouse in the non-treatment group died shortly before euthanization, and the colon was not preserved for histological study.) **o-t,** Evaluation of the optimized combinational niche formulation for colitis treatment in IL10 KO mice that spontaneously developed colitis at an early age. IL10 KO mice developed colitis symptoms at 10-11 weeks of age and received combinational niche treatment at weeks 11.5, 13.5 and 15.5. DAI score of colitis IL10 KO mice after receiving combinational niche treatments (**o**). Blood hemoglobin counts (**p**) and lymphocyte counts (**q**) recorded for IL10 KO mice after colitis treatments with combinational niche. Length of the colon of IL10 KO mice preserved at the study endpoint (17 weeks old) (r). Colon damage score (**s**) and representative H&E-stained colon sections (**t**) preserved from 17 week old IL10 KO mice after treatment with combinational niche. (n = 6) Data are presented as the mean ± s.e.m. All P values were analyzed using one-way (**c**,**g**,**l**,**r**) or two-way. (**b**,**d**,**f**,**h**,**k**,**m**,**o**,**p**,**q**,**s**) ANOVA with Tukey’s HSD multiple comparisons post-hoc test. * denotes p < 0.05, hence statistically significant. In the Dextran sodium sulfate-induced colitis model, disease activity index scores were compared based on data recorded up to day 10 (**b**,**f**), i.e., before the non-treated mice started to recover.

Next, we evaluated the therapeutic efficiencies of colon-specific immune niches bioengineered with non-immune checkpoint immunomodulators. Consistent with the weaker T cell inhibition efficiency observed for the leflunomide-encapsulated nanofibers or TGF-β1 mimetic peptide-encapsulated nanofibers *in vitro* (Extended Data Fig. 1 and Supplementary Fig. 16-18,20, 22-23), the leflunomide-encapsulated niche and TGF-β1 mimetic peptide encapsulated niche were less effective in relieving colitis symptoms (Fig. 3f,g) and promoting repair of colon epithelium damage (Fig. 3h,i) than immune checkpoint immunomodulator-functionalized niche. However, the colitis mice treated with the combination of leflunomide and TGF-β1 mimetic peptide-encapsulated niche showed full bodyweight recovery (Extended Data Fig. 4a), colon length maintenance (Fig. 3g and Extended Data Fig. 4b), reduced local myeloperoxidase activity (Extended Data Fig. 3a), and proinflammatory cytokine expression (Extended Data Fig. 3b,d). The improved therapeutic efficiency of the niche encapsulated with leflunomide and TGF-β1 mimetic peptide can be explained by the abilities of both immunosuppressive components to simultaneously inhibit CD8^+^ T cell activation and polarize classically-activated M1 macrophages through multiple regulatory pathways.

Based on these findings, we further evaluated the colon-specific immune niches prepared from the combinational nanofibers and colon extracellular matrix (combinational colon-specific immune niche). Unlike the earlier treatments, subcutaneous treatment with the combinational immune niche accelerated the amelioration of colitis symptoms compared to single immunomodulator-functionalized niches and promoted complete recovery within 7 days after treatment (Fig. 3f and Extended Data Fig. 4a,b). The treatment also reduced local myeloperoxidase activity (Extended Data Fig. 3a) and proinflammatory cytokine expression to baseline levels (Extended Data Fig. 3b) and increased the local production of pro-regulatory cytokines by nearly 3-fold compared to healthy mice (Extended Data Fig. 3c). Further histological analysis showed that the combinational niche was the only treatment that promoted complete recovery of epithelium damages within 7 days after treatment and prevented crypt loss (Extended Data Fig. 4c). Fecal microbial analysis at the study endpoint indicated that combinational niche treatment partly resolved microbiome dysbiosis (Supplementary Fig. 27 and Supplementary Table 4). To avoid fighting between male mice after the randomization (just before the colitis treatment) affecting the therapeutic study, most of the therapeutic studies were performed on female mice. Nevertheless, we were able to observe a similar effective therapeutic effect in male colitis mice that were randomized 1 week before the induction of colitis (Extended Data Fig. 5). This finding confirms that the subcutaneous combinational niche treatment effect is sex-independent.

Given the promising CD8^+^ T cell anergization and classically-activated M1 macrophage polarization abilities observed for the colon extracellular matrix *in vitro* (Fig. 2a-g), we performed control studies to investigate the therapeutic effect of colon extracellular matrix. Administrating a therapeutic dose of colon extracellular matrix (0.5 mg/treatment, the same quantity as in the combinational niche) failed to relieve colitis symptoms (Extended Data Fig. 6-7) and prevent colitis-associated colon epithelium damage and immune cell infiltration (Extended Data Fig. 6-7). Colitis treatment with a high-dose of colon extracellular matrix (4 mg/treatment) was as effective as combinational niche in ameliorating colitis symptoms (Extended Data Fig. 6-7). However, the treatment did not prevent inflammation-associated colon shortening, colon epithelium damage or immune cell infiltration into the lamina propria. These findings indicate that colon extracellular matrix alone was ineffective in treating ulcerative colitis. Obtaining a large quantity of colon tissues from patients with ulcerative colitis to engineer the immune niche would be difficult.

Additional control studies administered immune niches engineered from combinational nanofibers (Extended Data Fig. 6), pancreatic beta cells (MIN6 cells), and decellularized pancreas extracellular matrix (combinational pancreas-specific immune niche). The treatment failed to alleviate the colitis symptoms (Extended Data Fig. 8) or prevent inflammation-associated colon shortening (Extended Data Fig. 8). This finding confirms that the colon extracellular matrix releases pro-regulatory growth factors and cytokines, generating a colon-specific tissue microenvironment to modulate autoreactive immune cells that were observed *in vitro* (Fig. 2a-g). Additional control studies administered isotype control colon-specific immune niche engineered from colon epithelial cells, colon extracellular matrix and combinational isotype control nanofibers (composed of isotype control mouse immunoglobulin G-functionalized nanofibers, drug-free nanofibers-functionalized nanofibers and non-functionalized nanofibers). These showed moderate therapeutic effects and relieved the colitis symptoms (Extended Data Fig. 6-7). However, they were less effective in promoting the repair of epithelium damage and preventing immune cell infiltration into the lamina propria (Extended Data Fig. 6-7) than the combinational niche treatment. Nevertheless, these findings confirm that combinational nanofibers play a key role in ameliorating colitis symptoms.

Recognizing the translational challenges for bioengineering combinational colon-specific immune niche, we further investigated whether intraperitoneally-administered fully synthetic combinational nanofibers could treat ulcerative colitis since they can directly engage autoreactive immune cells in the abdominal cavity. Similar to the subcutaneously-administered combinational immune niche, the intraperitoneally-administered combinational nanofibers effectively suppressed colitis symptoms (Extended Data Fig. 9a,b), prevented inflammation-associated colon shortening (Extended Data Fig. 9c), promoted repair of epithelium damage (Extended Data Fig. 9d), and reduced immune cell infiltration into the colon (Extended Data Fig. 9d). Fecal microbial analysis showed that the intraperitoneal treatment is more effective than the subcutaneously-administered combinational niche treatment resolved microbiota dysbiosis (Supplementary Fig. 27 and Supplementary Table 4) and upregulated several commensal microbiomes (e.g., RF39). An *ex vivo* biodistribution study performed 5 days after the intraperitoneal treatment showed that most administered rhodamine-labeled nanofibers were distributed throughout the abdominal cavity (Supplementary Fig. 28), whereas the subcutaneously-administered combinational niche was retained at the injection site. These findings highlight the direct therapeutic potential combinational niche when autogenous colon epithelial cells and colon extracellular matrix are unavailable.

To demonstrate that combinational niche can ameliorate different types of colitis, we further evaluated combinational niche in the trinitrophenol-induced colitis model^52^ and IL10 KO mouse model of spontaneous colitis^55^. The trinitrophenol-induced colitis model is an established CD8^+^ T cell-driven relapsing colitis model in which trinitrophenol-sensitized CD8^+^ T cells attack trinitrophenol-modified colon epithelial cells^52^. In the TNP-induced colitis mice (Fig. 3j), treatment with a trinitrophenol-modified combinational niche (a combinational colon-specific immune niche engineered with trinitrophenol -modified colon epithelial cells) significantly relieved the severe colitis symptoms (e.g., rectal bleeding and rectum prolapse) within 24 h after treatment (Fig. 3k), minimized colitis-associated colon shortening (Fig. 3l and Supplementary Fig. 29a), and promoted repair of the ulcerated epithelium (Fig. 3m,n and Supplementary Fig. 29b). In the IL10 KO mouse model, mice spontaneously develop enterocolitis due to the dysfunction of regulatory T cells that cannot produce pro-regulatory IL10^55^. Repeated treatments with combinational niche upon colitis onset prevented disease progression (Fig. 3o), such as hematochezia, colitis-induced anemia (Fig. 3p), lymphocytic colitis (Fig. 3q), inflammation-induced colon shortening (Fig. 3r and Supplementary Fig. 30), and immune cell infiltration into the lamina propria (Fig. 3s-t and Supplementary Fig. 30). This finding confirms that the combinational niche treatment strategy is applicable to a broad range of colitis.

### Combinational colon-specific immune niche ameliorates colitis by inhibiting autoreactive CD8^+^ T cell activation and polarization of classically-activated M1 macrophages into alternatively-activated M2 macrophages

We performed immune profiling (Fig. 4a and Supplementary Fig. 31) to better understand different immune treatment^56^ action mechanisms in the dextran sodium sulfate-induced colitis model. Colitis significantly increased the frequencies of proinflammatory CD86^high^ classically-activated M1 macrophages and IFN-γ^high^ and/or TNF-α^high^ CD8^+^ T cells, and decreased the frequencies of CCR7^+^ CD4^+^ memory T cells and pro-regulatory FoxP3^+^ CD4^+^ regulatory T cells infiltrating the lamina propria compared to those in the healthy colon (Fig. 4a,b and Supplementary Fig. 32). 5 days after colitis treatment, combinational niche significantly reduced the frequencies of CD86^high^ classically-activated M1 macrophages (thus the classically-activated M1 macrophage-to-alternatively-activated M2 macrophage ratio; Fig. 4b and Supplementary Fig. 32) and IFN-γ^high^ and TNF-α^high^ CD8^+^ T cells compared to the non-treatment group (Fig. 4b and Supplementary Fig. 32), normalized the frequency of CD206^high^ alternatively-activated M2 macrophages to the background level (Fig. 4b and Supplementary Fig. 32), and increased the frequency of exhausted CD8^+^ T cells (e.g., CD69) (Supplementary Fig. 32). The treatment also significantly increased the frequency of CCR7^+^ CD4^+^ memory T cells compared to the non-treatment group (Supplementary Fig. 32), but did not normalize the frequency of regulatory T cells (Supplementary Fig. 32). Similar reductions in proinflammatory classically-activated M1 macrophages and CD8^+^ T cells were observed in the lamina propria of colitis mice receiving intraperitoneal treatment with combinational nanofibers (Fig. 4a,b and Supplementary Fig. 32). This result agrees with the reduced infiltration of CD8^+^ T cells, especially INF-γ^+^ CD8^+^ T cells, observed in the histological analysis, (Fig. 4c and Extended Data Fig. 3d) and the reduced therapeutic efficacy in CD8^+^ T cell- and macrophage-depleted colitis mice (Fig. 4d,e and Supplementary Fig. 33). The immune cell depletion study supports the anti-colitis effect of combinational niche observed in the IL10 KO mice. If the combinational niche works by modulating CD4^+^ T cells in both models, we should not observe any anti-colitis effect in the IL10 KO mice because its CD4^+^ T cells cannot produce IL10 to regulate the CD8^+^ T cells and macrophages. Further immune profiling of colons preserved 12 days post-treatment showed a similar reduction of in lamina propria-infiltrated CD8^+^ T cells and classically-activated M1 macrophages (Fig. 4a,b and Supplementary Fig. 32). However, the reductions were less significant than those in the non-treatment group (Fig. 4a,b and Supplementary Fig. 32) due to the recovery from inflammation after feeding with normal drinking water. Nevertheless, this mechanistic study confirms our hypothesis that the proposed combinational niche could directly inhibit CD8^+^ T cell activation and promote classically-activated M1 macrophages differentiation into alternatively-activated M2 macrophages.

We performed immunohistological analysis on preserved colon specimens 4 days after treatment to show that reduced CD8^+^ T cell and classically-activated M1 macrophage infiltration creates less inflammatory tissue microenvironments that allow healthy colon epithelial stem cells in crypts to repair epithelial damage^22, 23^. Without any colitis treatment, significant apoptosis of colon epithelial cells (TUNEL^+^ cells) in the villi and immune cell invasion were observed (Fig. 4f), and most colon epithelial cells at the crypt were arrested in the G0 phase or underwent apoptosis (i.e., Ki67^−^ cells; Fig. 4g). Therapeutic treatments with colon-specific immune niches significantly reduced immune invasion and apoptotic TUNEL^+^ epithelial cells in the villi, and increased the frequency of proliferating Ki67^+^ colon epithelial cells in the crypt (Fig. 4g). Specifically, treatment with combinational niche reduced apoptotic TUNEL^+^ colon epithelial cells in the villi by approximately 65% (Fig. 4f) and nearly completely repaired the damaged colon epithelium. The recovery of colitis-induced epithelial damage was observed after intraperitoneal treatment with the combinational nanofibers.

We performed a therapeutic study in a DSS-induced chronic colitis mouse model^53^ to demonstrate that the partial restoration of colonic CCR7^+^ CD4^+^ memory T cells after colitis treatment with combinational niche induces memory immune responses (Extended Data Fig. 10). Similar to the monophasic DSS dextran sodium sulfate-induced colitis study, mice that repeatedly received treatment with combinational niche showed complete bodyweight recovery (Extended Data Fig. 10), maintained colon length (Extended Data Fig. 10), and reduced inflammation-associated colon epithelium damage (Extended Data Fig. 10) after three colitis induction-recovery cycles. However, another experimental group that received only one treatment with combinational niche in the first colitis onset cycle also showed comparable recovery patterns to those who repeatedly received treatment (Extended Data Fig. 10). At the study endpoint, both groups of mice showed similar bodyweight recovery (Extended Data Fig. 10), colon lengths (Supplementary Fig. 26b-c), and colon damage scores (Extended Data Fig. 10). Hence, memory T cells likely develop in the first-phase of treatment, preventing further colitis development.

An additional immune profiling study was performed to understand the immune modulation mechanism of the s.c. administered combinational niche (Supplementary Fig. 34). Significant CD8^+^ T cell and macrophage infiltration was observed 3 days after administration, with approximately 63% and 10% of the graft infiltrated lymphocytes CD8^+^ and F4/80^+^ (Fig. 4h), respectively. Specifically, approximately 2% of the infiltrated CD8^+^ T cells expressed T cell exhaustion markers (e.g., CD69) and 13% of the F4/80^+^ macrophages highly expressed the CD86 classically-activated M1 macrophage marker (Fig. 4i). The magnitude of immune cell infiltration was significantly reduced 5 days after inoculation (Fig. 4j), but the relative frequencies of TNF-α^+^ activated CD8^+^ T cells and CD86^+^ classically-activated M1 macrophage had not significantly change (Fig. 4i). The relatively high number of CD8^+^ T cells infiltrated into the s.c. graft is consistent with our hypothesis that the colon-specific immune niche acts as a site for autoreactive CD8^+^ T cell modulation. On the other hand, the relatively low frequency of macrophages detected in the s.c. graft suggests that the anti-inflammatory growth factors and cytokines may be released from the colon extracellular matrix and systemically dampen the classically-activated M1 macrophage, as observed in other similar studies^57, 58^ and clinically-use Sargramostim^59^.

### Combinational immune niche ameliorates colitis-associated colorectal cancer

Chronic colonic inflammation increases the risk of developing colorectal cancer^60, 61^. Therefore, we assessed whether combinational colon-specific immune niche could prevent the development of colitis-associated colorectal cancer in an established azoxymethane/dextran sulfate sodium model of colorectal cancer^62, 63^ (Fig. 5a). Without treatment, azoxymethane/dextran sodium sulfate-induced colitis mice experienced more severe colitis and bodyweight loss than dextran sodium sulfate colitis mice (Fig. 5b and Supplementary Fig. 35). At the study endpoint, severe colon shortening (Fig. 5c) and higher tumor burdens were evident in the distal colons of azoxymethane/dextran sodium sulfate-induced colitis mice (Fig. 5d-e and Supplementary Fig. 35). In contrast, mice consistently responded to combinational niche treatment (Fig. 5b). At the study endpoint, treated mice completely recovered their bodyweight (Supplementary Fig. 35) and had 60% lower tumor burdens than the non-treatment group (Fig. 5d-e and Supplementary Fig. 35). This finding highlights the potential of using combinational niche to prevent colitis-associated colorectal cancer.

**Fig. 4.**
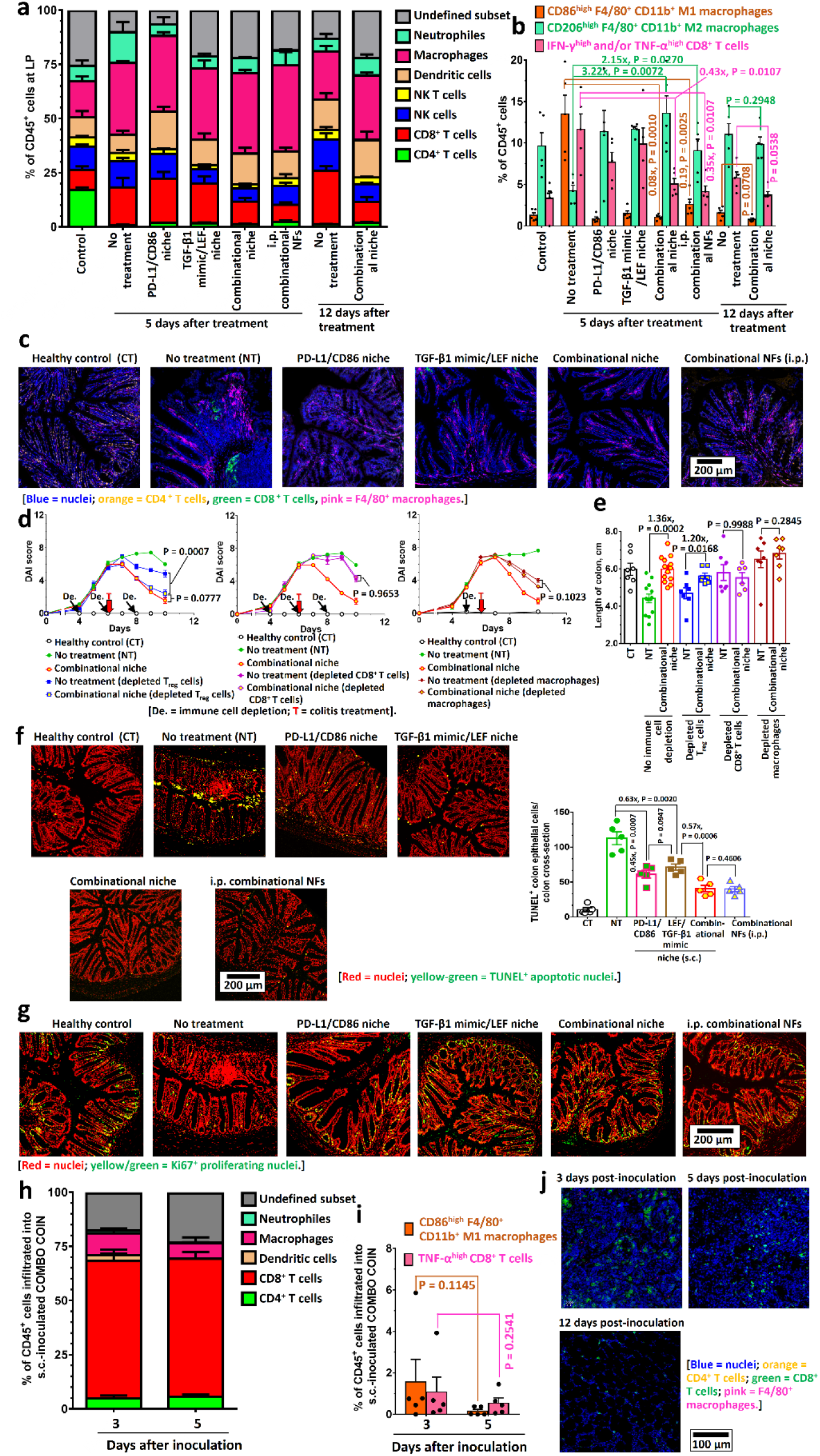
Mechanistic insight: combinational colon-specific immune niche ameliorated dextran sodium sulfate-induced colitis by inhibiting colon-specific CD8^+^ T cell activation and polarizing classically-activated M1 macrophages into alternatively-activated M2. **a-b,** Immune profiling of colons preserved from dextran sodium sulfate-induced colitis mice after different combinational niche treatments. Stacked bar graph representing the frequencies of each type of lamina propria-infiltrated lymphocyte at different time points after different combinational niche treatments (**a**). Frequencies of proinflammatory CD86^high^ F4/80^+^ CD11b^+^ classically-activated M1 macrophages, anti-inflammatory CD206^high^ F4/80^+^ CD11b^+^ alternatively-activated M2 macrophages, and proinflammatory IFN-γ^high^ and/or TNF-α^high^ CD8^+^ T cells in the lamina propria after different colitis treatments (**b**). (n = 5) **c**, Representative immunofluorescence images of colons preserved from DSS colitis mice, captured four days after different therapeutic treatments, reveal PL-infiltrated CD4^+^ T cells, CD8^+^ T cells and F4/80^+^ macrophages. **d-e**, Evaluation of the therapeutic efficiencies of combinational niche in regulatory T (Treg) cell-, CD8^+^ T cell- and macrophage-depleted dextran sodium sulfate -induced colitis mice. Regulatory T cells and CD8^+^ T cells were depleted (De.) by intraperitoneal administration of anti-CD25 or anti-CD8a at days 4, 6 and 8. Macrophages were depleted by a single intravenous administration of Clodrosome (clodronate-encapsulated liposomes) on day 5. All mice received colitis treatment (T) with combinational niche at day 6. Disease activity index scores of different immune cell-depleted mice after colitis treatment with combinational niche (**d**). Length of the colon of different immune cell-depleted mice measured at the study endpoint (**e**). (n = 6 or 7) **f**, Representative TUNEL- stained colon sections preserved from healthy and colitis mice 4 days after treatment with different niches. The TUNEL staining identified TUNEL^+^ apoptotic colon epithelial cells. (n = 5) **g**, Representative Ki67-stained colon sections preserved from health and colitis mice 4 days after treatment with different niches. The Ki67 stain highlights the proliferating colon epithelial stem cells in the colonic crypts. **h-j,** Immune profiling of combinational niche at different time points after subcutaneous inoculation in dextran sodium sulfate-induced colitis mice. Stacked bar graph representing the frequencies of each type of infiltrated lymphocytes at different time points after inoculation (**h**). (n = 5) Frequencies of proinflammatory CD86^high^ F4/80^+^ CD11b^+^ classically-activated M1 macrophages and TNF-α^high^ CD8^+^ T cells infiltrated into the niche at different time points after inoculation (**i**). Representative immunofluorescence images of subcutaneously inoculated combinational niche preserved 3, 5 or 12 days after inoculation. Data are presented as the mean ± s.e.m. All P values were analyzed using one-way (**e**) or two-way (**b**,**d**,**e**,**f**,**i**) test. ANOVA with Tukey’s HSD multiple comparisons post-hoc test. In the dextran sodium sulfate-induced colitis model, the disease activity index scores were compared based on data recorded up to day 10 (**d**) before the non-treated mice started to recover.

### Combinational colon-specific immune niche ameliorates kinase inhibition-associated colitis in cell signaling pathway-targeted cancer therapy

Phosphoinositide 3-kinase inhibitors, such as copanlisib^64, 65^, have been developed to suppress tumor growth by inhibiting the overexpressed phosphoinositide 3-kinase signal pathway in cancer cells. Colitis is one of the most common immune-related adverse events associated with phosphoinositide 3-kinase inhibitors due to their ability to deplete regulatory T cells^65, 66^. We used the B16OVA melanoma tumor model with copanlisib cancer treatment to show that combinational colon-specific immune niche can attenuate copanlisib-associated colitis without affecting its anticancer activity (Fig. 5f). Similar to a previous study^67^, copanlisib effectively inhibited B16OVA tumor growth (Fig. 5g) but induced mild colitis symptoms (Fig. 5h). The colitis symptoms worsened after feeding with 3% dextran sodium sulfate-containing drinking water (Fig. 5f,h). Colitis treatment with combinational niche effectively relieved colitis symptoms within 24 h after treatment (Fig. 5h). At the study endpoint, combinational niche treatment completely prevented inflammation-associated colon shortening (Fig. 5i and Supplementary Fig. 36) and facilitated partial repair of colon epithelium damage (Fig. 5j,k and Supplementary Fig. 36) without affecting the anticancer activity of copanlisib (Fig. 5g). These findings confirmed the therapeutic potential of combinational niche for treating colitis-triggered by regulatory T cell-depletion cancer therapy.

**Fig. 5.**
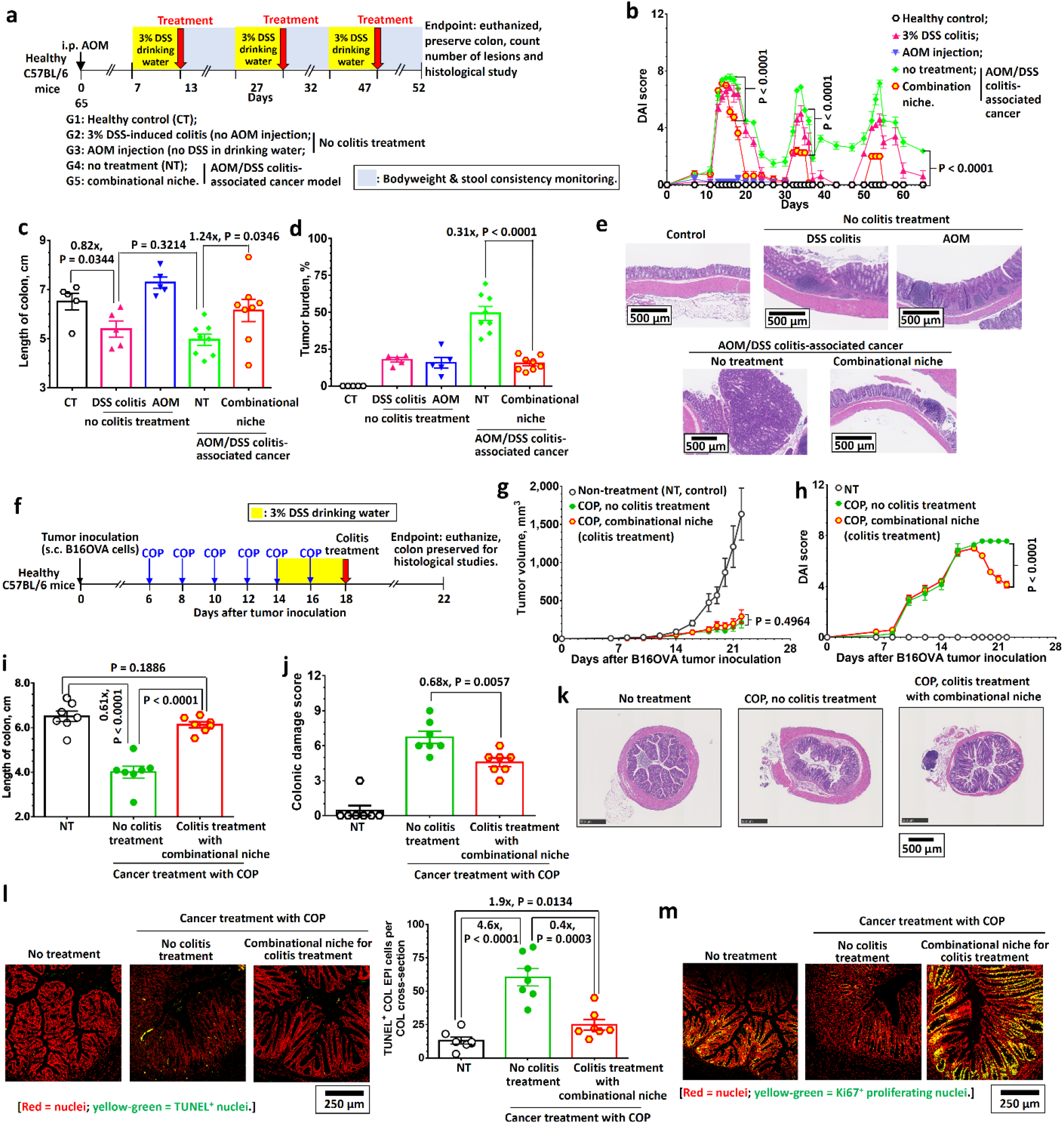
Combinational colon-specific immune niche effectively ameliorated colitis-associated colorectal cancer and phosphatidylinositol 3-kinase inhibition associated colitis in cell signaling pathway-targeted cancer treatment. **a**, Schematic of the treatment schedule for the evaluation of combinational colon-specific immune niche in preventing colitis-associated colorectal cancer through the azoxymethane/dextran sodium sulphate colitis-associated cancer model. **b,** Time-dependent disease activity index scores. **c,** Length of colons preserved at the study endpoint after different treatments, and representative digital photographs of colons from different control and treatment group mice harvested at the study endpoint. The preserved colons were stained with Alcian blue to identify the tumor boundaries. **d**, Colorectal cancer tumor burdens of azoxymethane/dextran sodium sulphate colitis mice after different treatments. **e**, H&E-stained histological images of preserved colons at the study endpoint after different treatments against azoxymethane/dextran sodium sulphate colitis. (n = 5, except azoxymethane/dextran sodium sulphate colitis-associated cancer groups with 8 mice per group.) **f,** Schematic of cancer and colitis treatment schedules for the evaluation of combinational niche in relieving copanlisib (COP)- induced colitis in B16OVA tumor model. **g-h,** Tumor growth curves (**g**) and disease activity index scores (**h**) recorded for B16OVA tumor-bearing mice after cancer treatment with copanlisib and colitis treatment with copanlisib. (n = 7) **i,** Length of colon preserved at the experimental endpoint (day 22 post-inoculation). **j-k,** Colon damage score (**j**) and representative H&E-stained colon sections (**k**) preserved from copanlisib-treated B16OVA tumor-bearing mice after cancer and colitis treatments. **l,** Representative TUNEL-stained colon sections preserved from healthy and colitis mice 4 days after treatment (day 22 post-inoculation of B16OVA tumor) with combinational niche. TUNEL staining identified TUNEL^+^ apoptotic colon epithelial cells. **m,** Representative Ki67-stained colon sections from copanlisib-treated B16OVA tumor-bearing mice preserved 4 days after colitis treatment with combinational niche. Ki67 staining highlights proliferating colon epithelial stem cells in the colonic crypts. Data are presented as mean ± s.e.m. All P values were analyzed using the one-way (**c**,**d**,**i**) or two-way (**b**,**g**,**h**,**j**,**l**) ANOVA with Tukey’s HSD multiple comparisons post-hoc test.

### Combinational niche effectively ameliorates immune checkpoint blockade-associated colitis without impairing anticancer efficacy

Dual blockade of inhibitory immune checkpoint pathways, such as the PD-(L)1 and CTLA- 4 pathways, can synergistically improve cancer immunotherapy efficacy but also increases the chance of immune-related adverse events^68-72^. Acute ulcerative colitis is a common immune-related adverse events that leads to the discontinuation of cancer treatment^68-72^. In the DSS-induced colitis mouse model, dual immune checkpoint blockade with anti-PD-1 and anti-CTLA-4 antibodies induced profound colitis symptoms and eventually led to severe inflammation-associated colon contraction (33% shorter than healthy mice) and immune cell infiltration into the colon mucosa (Supplementary Fig. 37).

A key concern regarding immunosuppressive treatments for immune-related adverse events is their potential interference with immune checkpoint blockade treatment. We hypothesized that an optimized immune checkpoint blockade treatment schedule would provide a therapeutic window for colitis treatment without impairing cancer treatment efficacy due to the gradual detachment of anti-PD-1 and anti-CTLA-4 antibodies from T cells (Fig. 6a and Supplementary Fig. 38). Therefore, we performed an *in vivo* efficacy study in the B16F10 tumor model to investigate whether our organ-specific colitis treatment affected the anticancer efficacy of dual immune checkpoint blockade immunotherapy. In the B16F10 melanoma tumor model (Fig. 6b), colitis treatment with combinational niche four days after the final dual immune checkpoint blockade treatment did not significantly affect cancer treatment efficacy (Fig. 6b). However, it significantly reduced colitis symptoms and colon inflammation at the study endpoint (Fig. 6c,d and Supplementary Fig. 39). Histological analysis confirmed that our colitis treatment significantly reduced epithelial damage and immune cell infiltration into the colon epithelium (Fig. 6f-I and Supplementary Fig. 39). Similar therapeutic effects were observed for the intraperitoneal treatment with combinational nanofibers (Fig. 6c,d,f-i), and colitis treatment did not affect immune checkpoint blockade treatment efficacy (Fig. 6b,e). Since combinational niche ameliorates colitis partially through the immunosuppressive PD-L1/PD-1 and CD86/CTLA-4 immune checkpoint pathways, we further investigated whether both colitis treatments would affect tumor antigen-specific CD8^+^ T cells. In the B16OVA tumor model (Fig. 6j), the frequency of tumor-infiltrated ovalbumin (OVA)-specific CD8^+^ T cell and expression of TIM-3 in the tumor preserved three days after colitis treatment was comparable with those who had not received colitis treatment (Fig. 6k and Supplementary Fig. 40). These findings show that colon-specific colitis treatment does not affect the anticancer activities of CD8^+^ T cells.

**Fig. 6.**
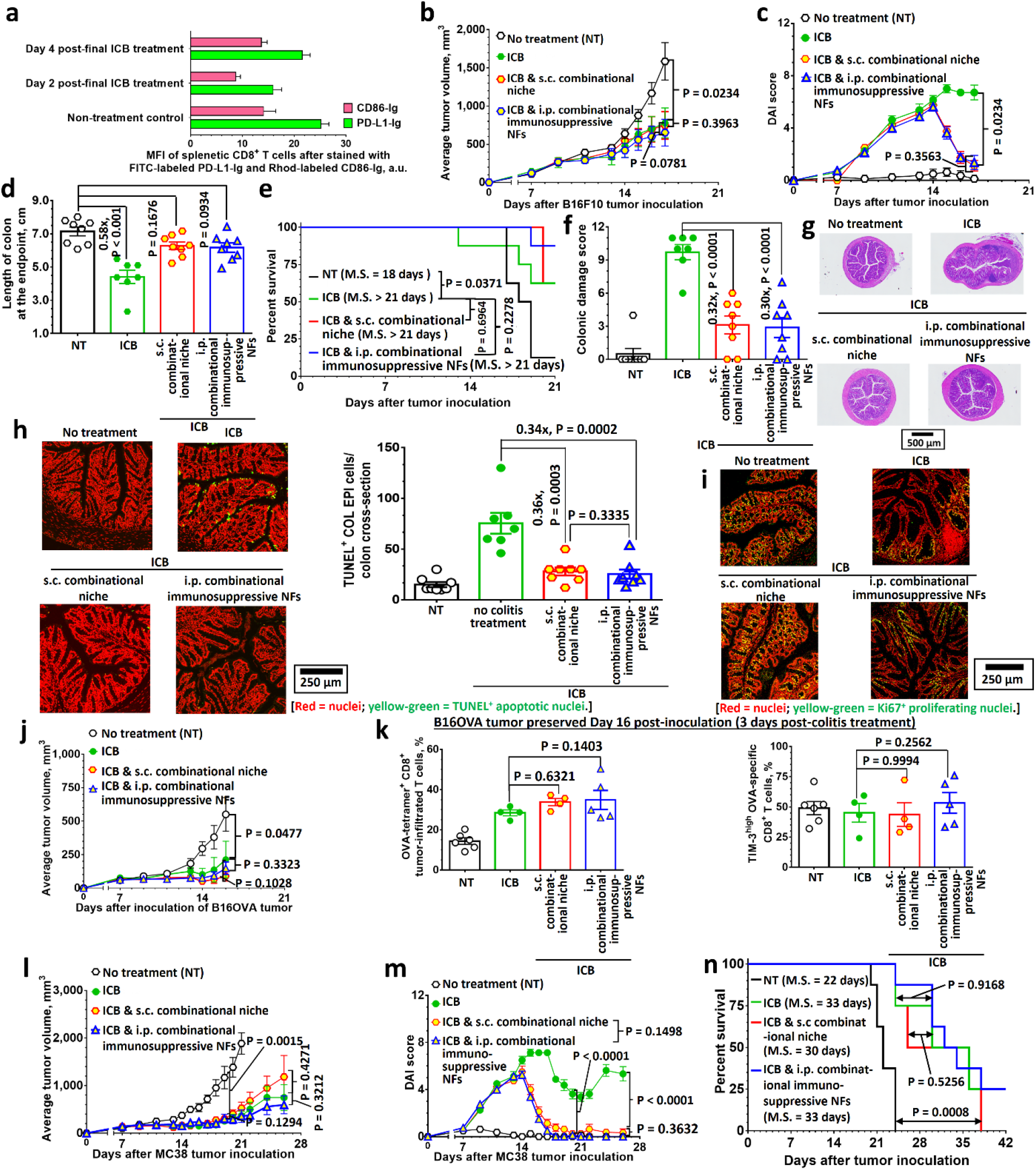
Subcutaneously-administered combinational colon-specific immune niche and intraperitoneal-administered combinational nanofibers are equally effective in ameliorating immune-related colitis without affecting the anticancer activities of immune checkpoint blockade (ICB). **a**, An *ex vivo* binding study quantified the binding affinities of splenetic CD8^+^ T cells to fluorescein isothiocyanate (FITC)-labeled PD-L1-Ig and rhodamine (Rhod)-labeled CD86- Ig after immune checkpoint blockade, as determined using fluorescence-activated cell sorting. **b- d**, Evaluation of subcutaneously-inoculated combinational niche and intraperitoneal-administered combinational nanofibers ameliorating immune-related colitis after immune checkpoint blockade with anti-PD-1 (100 µg/injection) and anti-CTLA-4 (100 µg/injection) in a B16F10 tumor model. Tumor growth curves (**b**), DAI scores (**c**), and colon length at the endpoint (**d**) after cancer immune checkpoint blockade and colitis treatment with subcutaneously-administered combinational niche and intraperitoneal-administered combinational nanofibers (n = 8). **e**, Survival of B16F10 tumor-bearing mice after receiving immune checkpoint blockade treatment with or without colitis treatment. **f-g,** Colon damage scores of colons preserved at the endpoint (**f**), and representative H&E-stained colon sections (**g**). **h,** Representative TUNEL-stained colon sections preserved from B16F10 tumor bearing mice after receiving cancer and colitis treatments. TUNEL staining identified TUNEL^+^ apoptotic colon epithelial cells. **i,** Representative Ki67-stained colon sections preserved from B16F10 tumor-bearing mice after receiving cancer and colitis treatments. Ki67 staining highlights proliferating colon epithelial stem cells in the colonic crypts. **j-k,** Evaluation of the impact of colitis treatment with subcutaneously-inoculated combinational niche or intraperitoneal-administered combinational nanofibers on immune checkpoint blockade treatment with anti-PD-l and anti-CTLA-4 in the B16OVA tumor model. Tumor growth curve (**j**). Frequencies of tumor-infiltrated OVA-specific CD8^+^ T cells and TIM-3^high^ OVA-specific CD8^+^ (exhausted) T cells 3 days after colitis treatment with combinational niche or combinational nanofibers (**k**). **l-n,** Evaluation of combinational niche and combinational nanofibers ameliorating immune-mediated colitis after immune checkpoint blockade with anti-PD-1 and anti-CTLA-4 in the MC38 synergetic colorectal cancer model and the impact of colitis treatment on subsequent maintenance immune checkpoint blockade treatment. Tumor growth curves (**l**), disease activity index scores (**m**), and survival curves (**n**) of MC38 tumor-bearing mice after receiving immune checkpoint blockade-based cancer treatment and colitis treatment with combinational niche or combinational nanofibers (n = 8). Data are presented as the mean ± s.e.m. All P values, except for the survival data, were analyzed using one-way (**a,k**) or two-way (**b-c,d,f,h,j-l,m,n**) ANOVA with Tukey’s HSD multiple comparisons post-hoc test. P values for the survival data (**e,n**) were analyzed using the log-rank (Mantel-Cox) test.

Guided by these promising treatment outcomes, we performed a separate anticancer efficacy study in the MC38 colorectal tumor model (Fig. 6l). Similar to the B16F10 tumor model, dual immune checkpoint blockade effectively inhibited tumor growth (Fig. 6l). However, colitis necessitated the discontinuation of immune checkpoint blockade treatment (Fig. 6m). Colitis treatment was performed four days after the second immune checkpoint blockade treatment and effectively reversed weight loss and ameliorated colitis symptoms (Fig. 6m and Supplementary Fig. 41). Although MC38 tumors are derived from colon epithelial cells, colitis treatment with combinational niche did not affect the anticancer efficacy of subsequent immune checkpoint blockade treatments (Fig. 6l,m). The colitis treatment protected mice from developing further colitis symptoms during subsequent maintenance immune checkpoint blockade immunotherapy (Fig. 6m and Supplementary 41). This is consistent with the development of colon epithelial cell-specific memory T cells observed in the chronic colitis model. Overall, this organ-specific colitis treatment did not affect the anticancer efficacy or survival benefit of immune checkpoint blockade treatment (Fig. 6n), that an optimized colitis treatment schedule would be tolerated during immune checkpoint blockade treatment.

## Discussion

Ulcerative colitis is an autoinflammatory disease that affects millions of individuals worldwide^1-3^. In addition to genetic and environmental factors^1-3, 10^, modern cancer therapies such as cell signaling pathway-targeted therapy^66^ and immune checkpoint blockade cancer immunotherapy^68-72^ increase the risk of developing ulcerative colitis. Indeed, ulcerative colitis is the most common immune-related adverse events leading to the discontinuation of phosphoinositide 3-kinase-targeted therapy and anti-CTLA-4-based cancer immunotherapy^65, 68- 72^. It often develops over time and can lead to life-threatening complications, such as bowel perforation, without appropriate treatment^1-3^. Ulcerative colitis remains incurable, but immunosuppressive therapies^11, 12^ have been developed to manage symptoms and achieve long-term remission. However, traditional immunosuppressive drugs suppress the entire immune system, and increase the probability of infection and cancer development^14^. Studies have found that patients with ulcerative colitis are nearly 2.5 times more likely to develop colorectal cancer than the general population^60, 61^. Therefore, developing new ulcerative colitis treatments is urgently needed.

Ulcerative colitis progression is widely believed to involve cross-talk between autoreactive T cells and innate immune cells (e.g., macrophages)^1-3^. One possible mechanism is that the proinflammatory cytokines (e.g., IFN-γ and TNF-α) produced by activated autoreactive CD8^+^ T cells induce residual colonic undifferentiated M0 macrophages to differentiate into proinflammatory classically-activated M1 macrophages and kill colon epithelial cells. In recent years, antigen-specific immunotherapies have been developed to treat autoimmune diseases by inducing antigen-specific immunotolerance by simultaneously presenting disease antigens and immunosuppressants (e.g., inhibitory immune checkpoint molecules and inhibitory cytokines) to autoreactive T cells. However, this strategy is limited by its ability to target adaptive immune cells, and prior knowledge of autoantigens is required.

To address all these challenges, we developed subcutaneous injectable immune niches that can simultaneously anergize autoreactive CD8^+^ T cells and polarize proinflammatory classically-activated M1 macrophages to anti-inflammatory alternatively-activated M2 macrophages through multiple immunosuppressive pathways. The immune niche was engineered from colon epithelial cells, decellularized colon extracellular matrix, and combinational nanofibers. The nanofibers were functionalized with two immunosuppressive immune checkpoint ligands (PD-L1 and CD86) to activate two key inhibitory immune checkpoint pathways in CD8^+^ T cells, leflunomide-encapsulated nanofibers to inhibit the proliferation of overreactive immune cells, and the pro-regulatory cytokine TGF-β1 mimetic peptide to inhibit CD8^+^ T cell proliferation and polarize classically-activated M1 macrophages. Comprehensive *in vitro* studies demonstrated that combinational nanofibers effectively anergized CD8^+^ T cells and induced classically-activated M1 macrophages polarization into alternatively-activated M2 macrophages. The immunosuppression effects were further improved in the presence of colon extracellular matrix due to its preserved pro-regulatory growth factors and cytokines.

Comprehensive correlative studies in the dextran sodium sulfate-induced colitis model confirmed that subcutaneous treatment with combinational niche could ameliorate colitis symptoms and speed up colon epithelial stem cells in the crypts in repairing the epithelial damage. The therapeutic effect is most significant for combinational niche formulation engineered with four different types of immunosuppressants. The control study showed that colon epithelial cells and colon extracellular matrix were essential for recapitulating colon-like tissue microenvironments to modulate autoreactive immune cells. Furthermore, *in vivo* studies showed that the combinational niche could ameliorate colitis symptoms, facilitate the recovery of ulcerated colon epithelium in a trinitrophenol-induced colitis model and minimize immune cell infiltration in IL10 KO mice.

Comprehensive immune profiling confirmed that the therapeutic treatment decreased the pro-inflammatory autoreactive CD8^+^ T cells and classically-activated M1 macrophages in the lamina propria, and increased the frequencies of anti-inflammatory alternatively-activated M2 macrophages However, the treatment did not completely return pro-regulatory colonic regulatory T cells to the background level. The histological study confirmed that the colon-specific immune niche treatments, especially combinational niche treatment, significantly reduced the number of apoptotic colon epithelial cells in the villi and preserved the viability of the colon epithelial stem cells in the crypts. In addition to ulcerative colitis, we demonstrated that treatment with combinational niche can prevent the development of colitis-associated colorectal cancer and ameliorate phosphoinositide 3-kinase inhibitor- and immune checkpoint blockade-induced colitis without affecting cancer treatment in established mouse models.

Recognizing the potential challenges of isolating colon epithelial cells and colon extracellular matrix from inflamed colon tissue in patients with colitis, we further evaluated an intraperitoneal administration of combinational nanofibers to treat ulcerative colitis. We found that the intraperitoneal administration of combinational nanofibers was as effective as the subcutaneously-administered combinational niche in relieving colitis symptoms in the dextran sodium sulfate-induced colitis model. Both immunosuppressive colitis treatments were also equally effective in promoting colon epithelial stem cells to repair the ulcerated colon epithelium. Furthermore, the *in vivo* study confirmed that intraperitoneally-administered immune checkpoint blockade were equally effective in relieving the colitis symptoms-triggered by immune checkpoint blockade in two established mouse tumor models, without affecting the cancer treatment efficiency. However, the *in vivo* biodistribution study indicated that the combinational nanofibers were rapidly distributed throughout the abdominal cavity after intraperitoneal injection. As in a similar study^73^, this challenge can be resolved by directly administrating the combinational nanofibers to the inflamed tissues identified through colonoscopy to prevent broad immunosuppression.

One limitation for our approach is the need for an autologous source of colonic epithelial to engineer colon-specific immune niche. This requires endoscopic biopsy of the patient, with risk of complications as well as financial cost. However, this clinical approach has already been explored in attempts to repair immune checkpoint blockade damage^74^. Future clinical translational will likely reply on modifying existing methods and clinical protocols to incorporate combinational niche as a treatment for ulcerative colitis. Another potential limitation is that colon-specific immune niche treatment does not fully capture the microbiome related biology in ulcerative colitis. It is known that microbiome plays an important role in the pathogenesis of ulcerative colitis though the exact biology is unclear^74^. To remedy the fact that the colon-specific immune niche construct does not contain any elements related to microbiome, colon-specific immune niche can be implanted orthotopically into the colon, as described in experiment using the trinitrophenol-induced model. The ultimate goal of this work is to clinically translate the colon-specific immune niche treatment. We recognize that the described colon-specific immune niche formulation is complex, which makes it difficult to translate. It has several different components: cells, nanofibers, extracellular matrix, and therapeutic agents. This high complexity makes chemistry, manufacturing and controls very difficult to achieve. Thus, future work will focus on simplify the formulation to enable easy manufacturing. We will aim to better understand the mechanisms of action to determine the most critical components of niche. We will also develop alternative formulation strategies to engineer colon-specific immune niche with fewer components and reactions.

In summary, we have shown that immunosuppressive nanofibers functionalized with immune checkpoint molecules, immunomodulatory drugs and pro-regulatory cytokines, can inhibit the activation of autoreactive CD8^+^ T cells and polarize classically-activated M1 macrophages into alternatively-activated M2 macrophages *in vitro*. We also demonstrated that combinational niches bioengineered from colon epithelial cells, decellularized colon extracellular matrix, and combinational nanofibers effectively ameliorated colitis by inhibiting autoreactive CD8^+^ T cell activation and allowing healthy colon epithelial cells to repair the damaged colon epithelium. Furthermore, the immune niche effectively prevented the development of colitis-associated colorectal cancer and ameliorated colitis-related immune-related adverse events in two established syngeneic tumor mouse models. The modular design of the immunosuppressive immune niche enables the on-demand introduction of alternative immunomodulators, targeted cells, and decellularized tissue to treat other autoimmune diseases.

## Online content

Any methods, additional references, Nature Research reporting summaries, source data, statements of data availability, and associated accession codes are available at …..

## Acknowledgments

We thank the University of Texas Southwestern Medical Center (UTSW) Animal Resource Center, UTSW ARC Diagnostic Laboratory, UTSW ARC Veterinary Services, UT Southwestern Electron Microscopy Facility, UTSW Quantitative Light Microscopy Core, UTSW Flow Cytometry Core, UTSW Histo Pathology Core, UTSW Proteomics Core, UTSW Microarray and Immune Phenotyping Core, UTSW Microbiome Research Lab, UTSW Preclinical Radiation Core Facility and UTSW Whole Brain Microscopy Facility at the University of Texas Southwestern Medical Center, UT Austin Texas Materials Institute, University of North Carolina at Chapel Hill Pathology Services Core, Akina, Inc. (West Lafayette, IN), RayBiotech Life, Inc. (Corners, GA), IDEXX BioAnalytics (Columbia, MI), and iHisto, Inc. (Salem, MA) for their assistance with the procedures in this article. UT Southwestern Electron Microscopy Facility is supported by the National Institutes of Health grant 1S10OD021685-01A1. UTSW Quantitative Light Microscopy Core is supported by the National Institutes of Health grant 1S10OD021684-01. UT Austin Texas Materials Institute is supported by the National Science Foundation Major Research Instrument Grant CBET-1624659. Pathology Services Core at the University of North Carolina-Chapel Hill is supported in part by an NCI Centre Core Support Grant 5P30CA016080-42. A.Z.W. was supported by the National Institutes of Health Grants R01GM130590 and R01EB025651.

## Author contributions

K.M.A., A.Z.W., J.P.Y.T. and J.E.W. conceived and designed the experiments. K.M.A. conceived the experiments. K.M.A. and A.Z.W. analyzed the data. K.M.A. and A.Z.W. co-wrote the paper. All authors discussed the results and edited the manuscript at all stages.

## Additional information

Supplementary information is available for this paper at ……

Reprints and permissions information is available at ……

Corresponds and requests for materials should be addressed to K.M.A. or A.W.

Publisher’s note: Springer Nature remains neutral with regard to jurisdictional claims in published maps and institutional afflictions.

## Methods Materials

Unless specified, all reagents were purchased from Fisher Scientific or Millipore Sigma.

### Synthesis of carboxylic acid (COOH)-functionalized diblock copolymer nanofibers

Carboxylic acid-functionalized poly(2-(methacryloyloxy)ethyl phosphorylcholine)- poly(hydroxypropyl methacrylate) (PMPC-PHPMA) amphiphilic diblock-copolymer nanofibers were prepared *via* an optimized two-step reversible addition-fragmentation chain-transfer (RAFT) polymerization. An idealized morphology-based phase diagram for PMPC-PHPMA was constructed from four PMPC macro-charge transfer agents (CTAs) with different PHPMA block lengths^35-37^. Briefly, 4 different PMPC macro-CTAs (building blocks) were synthesized *via* RAFT polymerization with actual degrees of polymerization (DPs, determined by ^1^H NMR spectroscopy) of 13, 21, 27, and 50. The PMPC macro-CTAs were chain extended with HPMA in H2O through RAFT aqueous dispersion polymerization with target DPs (for the HPMA block) varying from 140 to 490 at 25% solid content. The morphologies of all diblock copolymers were inspected *via* TEM.

The diblock copolymer NFs used in this study have a DP of 27 for the hydrophilic PMPC block and an actual DP of 280 for the hydrophobic PHPMA block. Briefly, a well-defined PMPC27 macro-CTA was first prepared *via* RAFT polymerization of MPC (catalog number: 67881-98-5; TCI Chemicals) using 4-cyano-4-(phenylcarbonothioylthio)pentanoic acid (CPDB; catalog number: 722995; Sigma) as a RAFT chain-transfer agent and 2,2’-azobis(2- methylpropionamidine) dihydrochloride (VA044; Fujifilm) as a thermal initiator at a target [MPC]:[CPDB]:[VA044] molar ratio of 30:1:0.25. The polymerization was performed at 40% solid content in a 1:9 water:methanol mixture at 50 °C for 150 min (approximately 90% conversion). The PMPC macro-CTA was isolated and purified by repeated precipitation into a 20- fold excess of a 1:20 methanol:acetone mixture (3 times). The collected polymer powders were first dried under vacuum before being redissolved in deionized water and lyophilized to remove residual solvents. The mean DP of the PMPC homopolymer calculated from the aromatic RAFT agent end group was 27, as determined by ^1^H NMR spectroscopy. The PMPC27 macro-CTA was chain extended *via* RAFT aqueous polymerization of HPMA (mixture of isomers, purity > 97%; catalog number: A17951.AK; Alfa Aesar) at a target [HPMA]:[PMPC27 CTA]:[VA044] molar ratio of 270:1:0.33. The polymerization proceeded at 50 °C for 180 min to yield the desired carboxylic acid-functionalized PMPC27-PHPMA280 diblock copolymer. To induce the diblock copolymer to self-assemble into nanofibers (“wormlike” micelles), polymerization was carried out at a solid content of 25 wt/wt%. ^1^H NMR spectroscopy (in CD3OD) confirmed the complete polymerization of the HPMA monomer under the optimized conditions. The pH of the as-prepared COOH-PMPC27-PHPMA280 diblock copolymer was determined to be approximately 4.0 after being diluted to 5 wt/wt% with deionized water. Rhodamine-labeled PMPC27-(PHPMA280-stat-RhodMA0.2) diblock copolymer was prepared using the same method except that 0.2 molar equivalents of methacryloxyethyl thiocarbamoyl rhodamine B (Polysciences, Inc.) was added to the HPMA during the second chain extension step.

### Functionalization of carboxylic acid-functionalized nanofibers with sulfo dibenzocyclooctyne (DBCO)-amine

Carboxylic acid-functionalized nanofibers were functionalized with sulfo dibenzocyclooctyne-amine *via* an amine-NHS ester coupling reaction. Briefly, a measured amount of carboxylic acid-functionalized NFs was diluted to 5 wt/wt% with deionized water before being activated with 15 molar equivalents (versus the COOH end-group) of 1-ethyl-3-(3-dimethylaminopropy)carbodiimide (EDC, 40 mg/mL in water) and N-hydroxysulfosuccinimide (sulfo-NHS, 40 mg/mL in water) at 20 °C for 1 h. The activated nanofibers were purified *via* 3 centrifugation (20,000 g, 20 min)-redispersion cycles to remove the unconsumed reagents. At the final centrifugation-redispersion cycle, the sulfo-NHS-activated nanofibers were suspended in 1X PBS. The sulfo-NHS-activated s nanofibers were allowed to react with 50 molar equivalents (vs. the COOH end-group) of sulfo dibenzocyclooctyne-amine (catalog number: 1227; Click Chemistry Tools) (20 mg/mL in water) at pH 8.5 (in 0.1 M PBS) and 20 °C in the dark for 12 h. The dibenzocyclooctyne-functionalized NFs were purified *via* equilibrium dialysis against water for 6 cycles.

### Preparation of azide-functionalized PD-L1-Ig, CD86-Ig and isotype IgG

PD-L1-Ig fusion protein (UniProt accession number: Q9EP73, mouse IgG1; catalog number: Pr00112-1.9; Absolute Antibody NA), CD86-Ig fusion protein (UniProt accession number: P42082, mouse IgG1; catalog number: Pr00226-1.9; Absolute Antibody NA) and mouse IgG1 isotype control (clone: MOPC-21, catalog number: BE083, BioXCell) were functionalized with azide-PEG8- NHS ester (catalog number: QBD10503; Sigma) *via* amine-NHS ester coupling reaction as previously reported^57^. The target degree of functionalization was 40, and the actual degree of functionalization was approximately 8 after purification *via* Pierce Polyacrylamide Spin Desalting Columns (7K MWCO; ThermoFisher), as determined by UV-visible spectroscopy after reaction with 50 molar equivalents of sulfo-dibenzocyclooctyne MB488 (catalog number: 11905; Click Chemistry Tools) and purification *via* spin desalting columns. Rhodamine- or fluorescein-labeled fusion proteins were also prepared for quantitative studies in the presence of 4 molar equivalents (vs. the fusion protein) of rhodamine B isothiocyanate (catalog number: 283923; Sigma) or fluorescein isothiocyanate (catalog number: 46950; Sigma) during functionalization with azide-PEG8-NHS.

### Fabrication of PD-L1 and/or CD86 nanofibers

PD-L1-, CD86-, and PD-L1/CD86- functionalized nanofibers (PD-L1 NFs, CD86 NFs, and PD-L1/CD86 NFs) were prepared *via* stain-promoted 1,3-dipolar cycloaddition as previously reported^26^. The target degree of functionalization for mono-functionalized NFs was 10 μg of azide-functionalized PD-L1-Ig or CD86-Ig (mono-functionalized nanofibers) per 1 mg of DBCO-functionalized nanofibers. The target degree of functionalization for dual-functionalized NFs was 5 μg of azide-functionalized PD-L1-Ig plus 5 μg of azide-functionalized CD86-Ig per milligram of DBCO-functionalized nanofibers. The bioconjugation was carried out at 5 wt/wt% of nanofiber solid content at 37 °C for 1 h, followed by 20 °C for another 1 h. Unconjugated fusion proteins were removed *via* 3 repeated centrifugations (20,000 g, 20 min)-redispersion cycles. Rhodamine-labeled azide-functionalized PD-L1-Ig and CD86-Ig were used for the quantification of ligand-conjugation efficiencies. The functionalized nanofibers were sterilized by 100 Gy of X-ray irradiation (*via* an XRD320 X-ray irradiator operated at 320 kV and 12.5 mA at a dose rate of 364 cGy/min) before *in vitro* and *in vivo* studies were conducted. Isotype mouse IgG-functionalized nanofibers were prepared via the sample method, except azide-functionalized isotype control IgG was used.

### Preparation of leflunomide-encapsulated nanoparticles

Azide-functionalized lefluomide-encapsulated poly(ethylene glycol)-block-poly(lactide-co-glycolide) nanoparticles were prepared *via* the nanoprecipitation method as previously reported^26^. The NPs were constructed from a polymer blend composed of 33.3 wt/wt% poly(lactide-co-glycolide) (Mn = 35,000-45,000 Da; catalog number: AP045; Akina, Inc.), 33.3 wt/wt% poly(ethylene glycol) methyl ether-block-poly(lactide-co-glycolide) (Mn = 3,000 + 36,000 Da; catalog number: AK029; Akina, Inc.), and 33.3 wt/wt% azide-functionalized poly(ethylene glycol)-block-poly(lactide-co-glycolide) (Mn = 15,000 + 5,000 Da; catalog number: AI085; Akina, Inc.). The target leflunomide (EP Reference Standard; catalog number: Y0000654; Sigma) loading was 10 wt/wt%, and the actual leflunomide loading was 8.7±1.5 wt/wt%, as determined by fluorescence spectroscopy^75^. Cy5-labeled leflunomide-encapsulated nanoparticle were prepared using a polymer blend composed of Cy5-labeled PLGA (Mn = 30,000-55,000 Da; catalog number: AV034; Akina, Inc.) instead of non-fluorescently labeled poly(lactide-co-glycolide) for the quantification of leflunomide-encapsulated nanoparticle conjugated to the dibenzocyclooctyne-functionalized nanofibers. Control drug-free azide-functionalized poly(ethylene glycol)-block-poly(lactide-co-glycolide) nanoparticles were prepared *via* the same method, except no drug was added to the polymer blend for the preparation of nanoparticles.

### Fabrication of leflunomide-encapsulated nanofibers

Leflunomide-encapsulated nanofibers (LEF NFs) were fabricated *via* stain-promoted 1,3-dipolar cycloaddition as previously reported^26^. The target degree of functionalization was 200 µg of azide-functionalized leflunomide-encapsulated nanoparticles per milligram of dibenzocyclooctyne-functionalized NFs. The bioconjugation was carried out at 5 wt/wt% solid content at 37 °C for 1 h, followed by 20 °C for another 1 h. Unconjugated leflunomide-encapsulated nanoparticles were removed *via* 3 repeated centrifugations (20,000 g, 20 min)-redispersion cycles. NFs functionalized with Cy5-labeled leflunomide-encapsulated nanoparticles were prepared *via* the same method for quantification of leflunomide-encapsulated nanoparticles conjugation efficiency *via* fluorescence spectroscopy. The actual quantity of leflunomide-encapsulated nanoparticles conjugated to the nanofibers was 118±8 µg of nanofibers per milligram of nanofibers, as quantified *via* fluorescence spectroscopy. Thus, each milligram of leflunomide-encapsulated nanofibers contained approximately 10 µg of encapsulated leflunomide. The functionalized nanofibers were sterilized by 100 Gy of X-ray irradiation (*via* an XRD320 X-ray irradiator operated at 320 kV and 12.5 mA at a dose rate of 364 cGy/min) before *in vitro* and *in vivo* studies were conducted. Control drug-free nanoparticle-functionalized nanofibers were prepared *via* the same method, except drug-free azide-functionalized poly(ethylene glycol)-block-poly(lactide-co-glycolide) nanoparticles were used instead of leflunomide-encapsulated nanofibers.

### Fabrication of TGF-β1 mimetic peptide-encapsulated nanofibers (TGF-β1 mimic NFs)

Cationic TGF-β1 mimetic peptide^31, 32^ (ACESPLKRQCGGG; purity > 95%; LifeTein, LLC) was physically adsorbed onto the anionic (unmodified) COOH-functionalized nanofibers at a target loading of 50 µg peptide per milligram of nanofibers. Prior to adsorption, the peptide was first dissolved in 1X PBS at 20 mg/mL and incubated at 20 °C for 4 h to allow oligomerization of the thiol-rich peptide. The peptide solution was then added to the nanofibers solution (final concentration = 50 mg/mL) and incubated at 20 °C for 2 h. Rhodamine-labeled TGF-β1 mimetic peptide (ACESPLKRQCGGGS-Lys(Rhodamine B); purity > 95%; LifeTein, LLC) was adsorbed onto the non-labeled NFs through the same method for *in vitro* release and T cell uptake studies. The functionalized NFs were sterilized by 100 Gy of X-ray irradiation (*via* an XRD320 X-ray irradiator operated at 320 kV and 12.5 mA at a dose rate of 364 cGy/min) before *in vitro* and *in vivo* studies were conducted.

### Mouse colon tissue decellularization

Decellularized colon extracellular matrix was isolated from healthy C57BL/6 mice (female, 6 – 7 weeks old) through a detergent-facilitated spin-decellularization protocol as previously reported^39, 40^. Briefly, freshly preserved colons were washed three times with 1X PBS before being incubated in 1% sodium dodecyl sulfate (SDS; ThermoFisher) in deionized water for 4 h with gentle shaking, during which the sodium dodecyl sulfate solution was changed once. The colons were then incubated with 1% Triton-X100 (Thermo Fisher) for another 1 h with gentle shaking. The acellular colons were then washed six times (20 min per wash) with 1X PBS (Gibco) supplemented with 1X Anti-Anti (Gibco). The acellular colons were than incubated in 1X PBS supplemented with 1X Anti-Anti at 37 °C for 24 h to destroy the residual microbes. The decellularized colons were then washed another 6 times with 1X PBS (5 min per wash) before being lyophilized. Lyophilized colons were then ball-milled (50 MHz, 5 min, 4 °C) into fine powders. The lyophilized colons were stored at -80 °C for up to 6 months before being used in the *in vitro* and *in vivo* studies. The DNA contents of freshly preserved mouse colon and decellularized colon extracellular matrix were determined by Promega QuantiFluor dsDNA binding dye (catalog number: E2671,Promega) according to the manufacturer’s protocol. Prior to all *in vitro* and *in vivo* studies, the decellularized colon extracellular matrix was suspended in serum-free high-glucose DMEM (for *in vitro* study) or 1X PBS (for *in vivo* study) and sterilized *via* 100 Gy of X-ray irradiation (*via* an XRD320 X-ray irradiator operated at 320 kV and 12.5 mA at a dose rate of 364 cGy/min) before further use in *in vitro* and *in vivo* studies. Three batches of mouse colons, with each batch pooled from 40 mice, were preserved for the whole study. Decellularized mouse pancreas extracellular matrix was isolated through a similar spin-decellularization method as previously reported^25^.

## Characterizations

### NMR spectroscopy

^1^H NMR spectra of crude and purified samples were recorded in a Bruker AN400 NMR spectrometer in the Biochemistry Nuclear Magnetic Resonance Core at the University of Texas Southwestern Medical Center (UTSW).

### Transmission electron microscopy (TEM) and scanning electron microscopy (SEM)

The morphologies of all NF samples were inspected through a JEOL 1400+ TEM microscope in the Electron Microscopy Core Facility at UTSW. An imaging study was performed on glow-discharged carbon-coated copper grids in which all samples were negatively stained with 1% phosphotungstic acid. The morphologies of decellularized colon extracellular matrix were inspected by SEM *via* an FEI Tecnai Spirit transmission electron microscope in the Electron Microscopy Core Facility at UTSW.

### Dynamic light scattering (DLS) and aqueous electrophoresis

Sphere-equivalent intensity-average diameters (Dh) and zeta potentials of all nanofibers samples were measured in a Malvern Zetasizer Pro instrument.

### Small-angle X-ray scattering measurements

Small-angle X-ray scattering measurements were performed in a Xenocs Ganesha 300-XL Small Angle Scattering Instrument in the Texas Materials Institute at the University of Texas at Austin. The small-angle X-ray scattering instrument was equipped with a microfocus Cu-Kα radiation source operated at 50 kV and 0.6 mA. X-ray diffraction patterns were recorded by a Dectris 300K detector to record extremely small-angle scattering data. The scattering intensity I(q) was recorded in the interval 0.005 Å < q < 0.015 Å^-1^, where q is defined as q = 4π/λ·sin(θ/2), in which θ is the scattering angle and λ is the wavelength of the X-ray source. Solution samples were analyzed in sealed thin-walled capillary tubes under vacuum conditions at ambient temperature (20 °C). The acquisition time for each sample was approximately 90 min. Data were corrected to give absolute intensities by measuring I0 directly on a Dectris detector. Two-dimensional scattering patterns were azimuthally integrated to a one-dimensional profile of intensity versus scattering vector using the two-dimensional data reduction utility SAXSGUI. Scattering patterns of the capillary tubes and PBS buffer or water (background solution) were collected and subtracted from the corresponding solution data. No attempt was made to convert the data to an absolute scale. Data analysis was performed by fitting the scattering curve to a polydisperse core-shell cylinder model using the least squares method in the IGOR Pro software pack (NIST, version 9.0; https://usaxs.xray.aps.anl.gov/software/irena) equipped with the Irena macro data analysis package^38^.

### Proteomic study

Matrisome and non-matrisome proteins in the native mouse colon samples, and decellularized colon extracellular matrix were analyzed using a Thermo Orbitrap Fusion^TM^ Lumos^TM^ Tribrid^TM^ mass spectrometer in the UTSW Proteomic Core. Three biologically independent mouse colon specimens preserved at different times and three batches of decellularized colon extracellular matrix were analyzed. For quantitative comparison, the protein contents were determined by Pierce bicinchoninic acid assay Proteins Assay Kit (Thermo Fisher) prior to the proteomic study.

### Growth factor antibody array

The relative levels of growth factors and cytokines in the non-decellularized mouse colon, decellularized mouse colon and decellularized mouse pancreas were quantified *via* the RayBio Human Growth Factor Antibody Array G-Series 1 by RayBiotech (Norcross, GA).

### Oscillatory rheology

An oscillatory rheology study of unmodified PMPC27PHPMA 280 diblock copolymer nanofibers (12.5 wt/wt% and 5 wt/wt%, in H2O) was performed in a TA Instrument AR2000 rheometer from Akina, Inc. The sample was equilibrated at a desired temperature from 45 to 5 °C (step size: 4 °C) for 5 min before each temperature-dependent measurement. Each measurement was performed at an angular frequency of 1 rad/s^-1^ and a stain amplitude of 1.0%.

### In vitro binding study

To confirm that nanofiber-conjugated PD-L1 and CD86 remained active, we performed an *in vitro* binding study with phycoerythrin (PE)-labeled anti-PD-L1 (clone: MIH6; catalog number: 153611; BioLegend) and A488-labeled anti-CD86 (clone: GL-1; catalog number: 153611; BioLegend). Briefly, 0.5 mg of unmodified and functionalized nanofibers were incubated with 5 µg of PE-labeled anti-mouse PD-L1 antibody (clone: 10F.9G2; catalog number: 124308; BioLegend) and 5 µg of A488-labeled anti-mouse CD86 antibody (clone: GL-1; catalog number: 105018; BioLegend) in fluorescence-activated cell sorting buffer at 20 °C for 30 min. After removal of unbound antibodies *via* 3 centrifugation-redispersion cycles, the degree of labeling was quantified by a Calibur flow cytometer at the University Flow and Mass Cytometry Facility at UTSW.

### *In vitro* release studies

The release of encapsulated leflunomide and physiosorbed rhodamine-labeled TGF-β1 mimetic peptide from the nanofibers was quantified *via* fluorescence spectroscopy (leflunomide: λex = 280±15 nm, λem = 410±15 nm^75^; rhodamine: λex = 550±15 nm, λem = 590±15 nm). Briefly, 30 mg/mL of the NFs were incubated against 4 times their volume in 1X PBS (in the presence or absence of 5 mM of dithiothreitol (DTT)) through Slide-A-Lyzer MINI Dialysis Units (20,000 MWCO; ThermoFisher) at 37 °C in the dark. At the desired time points, a known volume of the sample was collected from the dialysis device and diluted to 5 mg/mL. The amount of leflunomide and rhodamine-labeled TGF-β1 mimetic peptide retained in the NFs was quantified and compared with the fluorescent intensities of leflunomide and rhodamine in the crude NF samples *via* fluorescence spectroscopy.

The *in vitro* CD8^+^ T cell uptake of the released rhodamine-labeled TGF-β1 mimetic peptide from the TGF-β1 mimetic peptide-encapsulated nanofibers was quantified by fluorescence-activated cell sorting and confocal laser scanning microscopy. CD8^+^ T cells were harvested from the splenocytes of healthy C57BL/6 mice through the EasySep^TM^ Mouse CD8^+^ T Cell Isolation Kit (STEMCELL Technologies, Cambridge, MA) according to the manufacturer’s instructions. Briefly, isolated CD8^+^ T cells (50,000 cells per well in a 96-well plate) were cultured with 10 µg of nanofibers containing 0.5 µg of TGF-β1 mimetic peptide in T cell culture medium (100 µL of medium per well) under physiological conditions for up to 48 h. At desired time points, the samples were removed, washed 3 times with fluorescence-activated cell sorting buffer, and fixed before fluorescence-activated cell sorting or confocal laser scanning microscopy studies were conducted. FACS samples were analyzed in a Calibur flow cytometer at the University Flow and Mass Cytometry Facility at UTSW. The confocal laser scanning microscopy images were recorded with a Zeiss LSM880 confocal microscope at the Quantitative Light Microscopy Core at UTSW.

## In vitro studies

### Cell culture

Mouse primary colonic epithelial cells (catalog number: C57-6047, passage 1) isolated from healthy C57BL/6 mice were purchased from Cell Biologics, Inc. Primary mouse colonic epithelial cells and fibroblasts were cultured in Complete Epithelial Cell Medium with Supplement Kit (supplemented with 2% FBS, 5 ng/mL epidermal growth factor, and 1X insulin-transferrin-selenium; catalog number: M6621-Kit; Cell Biologics, Inc.) in gelatin-coated tissue culture flasks. Colon epithelial cells at passage 4 were used for *in vitro* and *in vivo* studies. Mouse macrophages (catalog number: C57-2313F) isolated from C57BL/6 mice were purchased from Cell Biologics, Inc. Macrophages were cultured in RPMI-1640 tissue culture medium (Gibco) supplemented with 10% FBS, and 1X Anti-Anti (Gibco). The primary cells were cultured in plasma-treated tissue culture flasks or well plates. Macrophages were detached and bypassed using scraping method.

B16F10 cell lines were obtained from the Tissue Culture Facility at the University of North Carolina (UNC) Lineberger Comprehensive Cancer Center at the UNC School of Medicine under the Material Transfer Agreement. B16F10 cells were cultured in high-glucose DMEM (4.5 g/L glucose; Gibco), 10% FBS (Corning), and 1X Anti-Anti (Gibco). Passage 13 or 14 B16F10 cells was used in the *in vitro* and *in vivo* studies.

MC38 cells were purchased from Kerafast. These cells were cultured in high-glucose DMEM supplemented with 10% FBS, 1X Anti-Anti (Gibco), 0.1 mM of non-essential amino acids (Gibco), 1 mM of sodium pyruvate, 2 mM GlutaMAX (Gibco), 10 mM HEPES buffer (Gibco), and 0.5 mg/mL gentamicin (Gibco) for positive selection. MC38 cells at passage 8 or 9 were used in the *in vitro* and *in vivo* studies.

B16OVA cells were kindly provided by Dr. Jonathan Serody in the UNC Lineberger Comprehensive Cancer Center at the UNC School of Medicine. B16OVA cells were cultured under positive selection conditions in RPMI 1640 culture medium (Gibco) supplemented with 10% FBS (Corning), 1X Anti-Anti (Gibco), 2 mM of GlutaMAX (Gibco), 0.1 mM non-essential amino acid (Gibco), 1 mM sodium pyruvate (Gibco), and 0.5 mg/mL Geneticin (Gibco) for positive selection. B16OVA cells at passage 15 or 16 were used in the *in vitro* and *in vivo* studies.

MIN6 cells were purchased from American Type Culture Collection. These cells were cultured in high-glucose DMEM supplemented with 20% FBS (Corning) and 1X Anti-Anti (Gibco). MIN6 cells at passage 29 were used in the *in vivo* study.

Primary colon epithelial cells were bypassed using TrypLE cell dissociation buffer (Gibco) and detached according to the manufacturer’s protocol. All cell lines were bypassed using 0.05% trypsin (Gibco) according to the manufacturer’s protocol. Macrophages were detached and bypassed using the scraping method.

Unless specified, red blood cells were depleted from splenocytes *via* ACK Lysing Buffer (Gibco) prior to conducting all *in vitro* studies. Red blood cell-depleted splenocytes and isolated CD8^+^ T cells were cultured in RPMI 1640 medium (Gibco) supplemented with 10% FBS (Sigma), 1X Anti-Anti (Gibco), and 2 mM GlutaMAX (Gibco).

### In vitro cell proliferation studies

The *in vitro* toxicities of different functionalized nanofibers to colon epithelial cells were evaluated using a 3-(4,5-dimethylthiazol-2-yl)-5-(3- carboxymethoxyphenyl)-2-(4-sulfophenyl)-2H-tetrazolium (MTS) assay. Briefly, colon epithelial cells (10,000 cells per well, 1/100 time used in the *in vivo* study) were cultured with unconjugated PD-L1 (0.1 µg/well, or 0.05 µg/well for the combo formulation), unconjugated CD86 (0.1 µg/well, or 0.05 µg/well for the combo formulation), small-molecule leflunomide (0.1 µg/well), free TGF- β1 mimetic peptide (0.5 µg/well) or 10 µg/well of each functionalized nanofibers (or a total of 30 µg/well combining all three different functionalized NFs) in complete colon epithelial cell culture medium (100 µL of medium per well) in a gelatin-coated 96-well plate for 3 days. The viabilities of colon epithelial cells were evaluated using the Promega CellTiter 96^TM^ AQueous One Solution Cell Proliferation Assay (MTS assay) according to the manufacturer’s protocol. The *in vitro* toxicity of copanlisib against B16OVA cells was determined by MTS assay after co-cultured with 0.1 – 10 µM copanlisib (catalog number: HY15346A100MG, Med Cehem Express, LLC.) in complete medium for 48 h.

The proliferation of colon epithelial cells cultured in different quantities of decellularized colon extracellular matrix was evaluated by MTS assay. Briefly, a desired concentration of decellularized colon extracellular matrix suspended in phenol red-free high-glucose DMEM was added to non-treated 96-well plates (with half of the wells pre-coated with gelatin) to achieve 0 – 6 µg of extracellular matrix per well (50 µL of medium per well) and air-dried in a laminar airflow tissue culture hood at 20 °C for 48 h. The plate was then sterilized *via* 100 Gy of X-ray irradiation (*via* an XRD320 X-ray irradiator operated at 320 kV and 12.5 mA at a dose rate of 364 cGy/min) before being briefly washed twice with 1X PBS. Colon epithelial cells, either suspended in complete colon epithelial cell culture medium (supplemented with 2% FBS and growth factors) or suspended in colon epithelial cell culture medium with half of the supplements, were added to the extracellular matrix-coated plate at a cell density of 2,000 cells per well and cultured for 8 days. Representative optical microscope images were recorded at day 8 through a Nikon Digital Sight optical microscope to record the growth conditions. Cell proliferation was evaluated using the Promega CellTiter 96^TM^ AQueous One Solution Cell Proliferation Assay (MTS assay) according to the manufacturer’s protocol. The viability of colon epithelial cell growth in gelatin-coated wells in complete cell culture medium (in the absence of colon extracellular matrix) was used as a 100% viable reference.

### In vitro T cell assays

Naïve mouse CD8^+^ T cells were isolated from healthy C57BL/6 mice (female, 8-10 weeks old) using EasySep^TM^ Mouse CD8^+^ T Cell Isolation Kit (STEMCELL Technologies, Cambridge, MA) according to the manufacturer’s instructions. In the T cell proliferation study, naïve CD8^+^ T cells were further labeled with carboxyfluorescein succinimidyl ester (CFSE) *via* CellTracer CFSE (Invitrogen) according to the manufacturer’s protocol. TNP-specific (sensitized) CD8^+^ T cells were generated *via* an established method^51^, ^52^. Briefly, naïve mouse splenocytes were preserved from healthy C57BL/6 mice (female, 8 to 10 weeks old). To obtain trinitrophenol-sensitized splenocytes, trinitrophenol-modified splenocytes were first engineered by incubating naïve splenocytes with 2,4,6-trinitrobenzene sulfonic acid in HBBS buffer for 15 min according to the published protocol^51^, ^52^. Native splenocytes were sensitized by incubation with trinitrophenol-modified splenocytes at a 1:20 ratio for 4 days. trinitrophenol-sensitized splenocytes were used for *in vitro* studies, and sensitized CD8^+^ T cells were further isolated. trinitrophenol-modified colon epithelial cells for antigen-specific studies were generated by incubating 2,4,6-trinitrobenzene sulfonic acid in HBBS buffer for 15 min according to the published protocol^51^, ^52^. To quantify the viability of the colon epithelial cells after being co-cultured with splenocytes, the colon epithelial cells were loaded with calcein *via* calcein acetoxymethyl cell-permeant dye (Thermo Fisher), according to the manufacturer’s protocol, prior to the viability study.

To evaluate the inhabitation efficiencies of different immunosuppressive nanofibers and colon extracellular matrix in naïve CD8^+^ T cells, CFSE-labeled naïve CD8^+^ T cells (5×10^4^ cells per well in a 96-well plate) were stimulated using Dynbeads Mouse T-Activator CD3/CD28 (T cell activation beads; Gibco) at a 1:1 molar ratio with 1U/well of recombinant mouse IL-2 (Gibco) in complete T cell culture medium (phenol red-free RPMI 1640 culture medium supplemented with 10% FBS, 1X Anti-Anti, and 1X GlutaMAX Supplement) in the presence of 0.1 µg (total amount of immunoglobulin fusion proteins, e.g., 0.05 µg of PD-L1-Ig plus 0.05 µg of CD86- Ig)/well of free or nanofibers-conjugated PD-L1-Ig and/or CD86-Ig, 0.1 µg/well of small-molecule or nanofibers-encapsulated leflunomide, 0.5 µg/well of free/nanofibers-bound TGF-β1 mimetic peptide, and/or 5 µg/well of colon extracellular matrix. Further *in vitro* study was performed at one-fourth doses of combinational immunosuppressive nanofibers and colon extracellular matrix, i.e., each well contained the combination of PD-L1/CD86-functionalized nanofibers containing 12.5 ng of NF-conjugated PD-L1-Ig and 12.5 ng of NF-conjugated CD86-Ig, leflunomide-encapsulated nanofibers containing 25 ng of encapsulated leflunomide and TGF-β1 mimetic peptide-encapsulated nanofibers containing 125 ng of TGF-β1 mimetic peptide, and/or 1.25 µg/well of colon extracellular matrix. The cells were allowed to culture for 48 h. The stimulated T cells were then analyzed in a Calibur flow cytometer or LSRII flow cytometer at the University Flow and Mass Cytometry Facility at the UTSW. Each FACS analysis contained 20,000 singlet cells.

To evaluate the expression of inhibitory markers (exhaustion markers) on the surface of CD8^+^ T cells after being cultured with different ISM NFs, naïve CD8^+^ T cells (5×10^4^ cells per well in a 96-well plate) were stimulated using Dynbeads Mouse T-Activator CD3/CD28 (T cell activation beads; Gibco) at a 1:1 molar ratio with 1 U/well of recombinant mouse IL-2 (Gibco) in complete T cell culture medium (phenol red-free RPMI 1640 culture medium supplemented with 10% FBS, 1X Anti-Anti, and 1X GlutaMAX Supplement) in the presence of 0.1 µg (total amount of Ig, e.g., 0.05 µg of PD-L1-Ig plus 0.05 µg of CD86-Ig)/well of free or nanofiber-conjugated PD-L1-Ig and/or CD86-Ig, 0.1 µg/well of small-molecule or nanofiber-loaded leflunomide, 0.5 µg/well of free/nanofiber-bound TGF-β1 mimetic peptide, and/or 5 µg/well of colon extracellular matrix. The cells were allowed to culture for 48 h. The T cells were then stained with PE-labeled anti-mouse TIM-3 antibody (clone: RMT3-23; catalog number: 119704; BioLegend) and APC-labeled anti-mouse LAG-3 antibody (clone: C9B7W; catalog number: 125210; BioLegend) to quantify the inhibitory markers. Stained T cells were then analyzed in a Calibur flow cytometer or LSRII flow cytometer at the University Flow and Mass Cytometry Facility at UTSW. Each FACS analysis contained 20,000 singlet cells.

To demonstrate that the combinational immunosuppressive nanofibers and colon extracellular matrix inhibit colon antigen-specific CD8^+^ T cell activation, non-sensitized and TNP trinitrophenol-sensitized splenocytes (1.5×10^5^ cells per well in a 96-well plate) were cultured with pre-seeded trinitrophenol-modified colon epithelial cells (1×10^4^ cells per well in a 96-well plate) (or pre-seeded unmodified COL EPI cells as a control) in the presence of (i) combinational immunosuppressive nanofibers: 10 µg/well of PD-L1/CD86 nanofibers (containing 0.05 µg of PD- L1-Ig and 0.05 µg of CD86-Ig), 10 µg/well of leflunomide-encapsulated nanofibers (containing 0.1 µg of encapsulated leflunomide), and 10 µg/well TGF-β1 mimic nanofibers (containing 0.5 µg of TGF-β1 mimetic peptide), (ii) combinational immunosuppressants: 0.05 µg/well of unconjugated PD-L1-Ig, 0.05 µg/well of unconjugated CD86-Ig, 0.1 µg/well of small-molecule leflunomide, and 0.5 µg/well of TGF-β1 mimetic peptide, (iii) 5 µg/well of colon extracellular matrix, and (iv) the combination of combinational immunosuppressive nanofibers and colon extracellular matrix. The cells were cultured for 48 h and then stained first with A488-labeled anti-mouse CD8a antibody (clone: 53-6.7; catalog number: 100723; BioLegend), fixed with 10% neutral buffered formalin, and permeabilized with Intracellular Staining Permeabilization Wash Buffer (BioLegend) before being stained with PE-labeled anti-mouse IFN-gamma antibody (clone: XMG1.2; catalog number: 505808; BioLegend) and APC-labeled anti-human/mouse granzyme B recombinant antibody (clone: OA16A02; catalog number: 372204; BioLegend) to quantify the T cell activation markers. Alternatively, cells were stained with A488-labeled anti-mouse CD8a antibody (clone: 53-6.7; catalog number: 100723; BioLegend), PE-labeled anti-mouse TIM-3 antibody (clone: RMT3-23; catalog number: 119704; BioLegend), and APC-labeled anti-mouse LAG-3 antibody (clone: C9B7W; catalog number: 125210; BioLegend) to evaluate the T cell inhibitory markers on the surface of the sensitized CD8^+^ T cells. Stained cells were then analyzed in a BD LSR II flow cytometer at the University Flow and Mass Cytometry Facility at UTSW. Each FACS analysis contained 20,000 singlet cells.

The change in the number of IFN-gamma secreting cells cultured in the presence of COMBO ISM NFs was quantified *via* a mouse IFN-gamma ELISpot PLUSE Kit (ALP) (Mabtech, Pittsburgh, PA). In an initial study, non-sensitized and trinitrophenol-sensitized splenocytes (1.5×10^5^ cells per well in a 96-well ELISpot plate) were cultured with trinitrophenol-modified colon epithelial cells (1×10^4^ cells per well in a 96-well ELISpot plate) (or pre-seeded unmodified colon epithelial cells as a control) in the presence of (i) combinational nanofibers: 10 µg/well of PD-L1/CD86-functionalized nanofibers (containing 0.05 µg of PD-L1-Ig and 0.05 µg of CD86- Ig), 10 µg/well of leflunomide-encapsulated nanofibers (containing 0.1 µg of encapsulated LEF), and 10 µg/well TGF-β1mimetic peptide-encapsulated nanofibers (containing 0.5 µg of TGF-β1 mimetic peptide) or (ii) combinational immunosuppressive molecules: 0.05 µg/well of unconjugated PD-L1-Ig, 0.05 µg/well of unconjugated CD86-Ig, 0.1 µg/well of small-molecule leflunomide, and 0.5 µg/well of TGF-β1 mimetic peptide. The cells were cultured in the IFN-gamma ELISpot plate for 48 h. The number of IFN-gamma–positive spots were generated *via* the manufacturer’s protocol and counted through a Zeiss SteREO Discovery microscope. In further studies, trinitrophenol-sensitized splenocytes (1.5×10^5^ cells per well in a 96-well ELISpot plate) were cultured with trinitrophenol-modified colon epithelial cells (1×10^4^ cells per well in a 96-well ELISpot plate) in the presence of one-fourth of immunosuppressants, i.e., combinational immunosuppressive nanofibers containing PD-L1/CD86-encapsulated nanofibers containing 12.5 ng of nanofiber-conjugated PD-L1-Ig and 12.5 ng of nanofiber-conjugated CD86-Ig, leflunomide-encapsulated nanofibers containing 25 ng of encapsulated leflunomide and TGF-β1 mimic nanofibers containing 125 ng of TGF-β1 mimetic peptide, and/or 1.25 µg/well of colon extracellular matrix.

The viabilities of calcein-loaded trinitrophenol-modified colon epithelial cells (1×10^4^ cells per well in a black 96-well plate) after being co-cultured with naïve and trinitrophenol-specific splenocytes (1.5×10^5^ cells per well in a black 96-well plate) in the absence or presence of combinational nanofibers (or combinational immunosuppressive molecules), and/or colon extracellular matrix (5 µg/well) were determined by quantifying the calcein retained in the colon epithelial cells 48 h after co-treatment according to the manufacturer’s protocol. The specific cell killing of each treatment group was calculated according to a published method^76^. To demonstrate that the co-cultured combinational immunosuppressive nanofibers induced CD8^+^ T cell exhaustion, the co-cultured splenocytes were further stained with APC-labeled anti-mouse CD8a antibody (clone: 53-6.7; catalog number: 100712; BioLegend) and PE-labeled anti-mouse TIM-3 antibody (clone: RMT3-23; catalog number: 119704; BioLegend). The stained cells were then mounted with ProLong^TM^ Diamond Antifade Mountant with DAPI and imaged through a Zeiss LSM880 confocal microscope at the Quantitative Light Microscopy Core at UTSW.

### *In vitro* macrophage assays

Classically-activated M1 macrophages were obtained by culturing mouse non-differentiated M0 macrophages in complete macrophage culture medium supplemented with 50 ng/mL recombinant IFN-γ (catalog number: RP8617, Invitrogen) and 1 µg/mL lipopolysaccharides (catalog number: 497603, Invitrogen) for 36 h, as previously reported^77, 78^. The induced classically-activated M1 macrophages were washed three times with RPMI-1640 (Gibco) before detachment *via* the scraping method for further studies.

To demonstrate that combinational immunosuppressive nanofibers and colon extracellular matrix can directly polarize classically-activated M1 macrophages to alternatively-activated M2 macrophages, induced classically-activated M1 macrophages (1×10^4^ cells per well in a coverglass chamber (catalog number: GBL112358, Grace Bio-Lab), same surface area as in 96-well plate) were cultured with 0.1 µg (total amount of Ig-fusion protein(s), e.g., 0.05 µg of PD-L1-Ig plus 0.05 µg of CD86-Ig)/well of nanofiber-conjugated PD-L1-Ig and/or CD86-Ig, 0.1 µg/well of nanofiber-encapsulated leflunomide, 0.5 µg/well of nanofiber-bound TGF-β1 mimetic peptide, and/or 5 µg/well of colon extracellular matrix for 48 h. The cells were washed, and stained with PE-labeled anti-mouse CD86 antibody (clone: GL-1, catalog number: 105007). The washed and stained cells were then fixed with 10% neutral buffered formalin, and permeabilized with Intracellular Staining Permeabilization Wash Buffer (BioLegend) before being stained with Alexa Fluor 488-labeled anti-mouse Arginase 1 antibody (clone: 658922; catalog number: IC8026G; R&D Systems). The stained cells were then mounted with ProLong^TM^ Diamond Antifade Mountant with DAPI and imaged through a Zeiss LSM880 confocal microscope at the Quantitative Light Microscopy Core at UTSW.

The viabilities of calcein-loaded trinitrophenol-modified colon epithelial cells (1×10^4^ cells per well in a black 96-well plate) after being co-cultured with classically-activated M1 macrophage (1×10^4^ cells per well in a black 96-well plate) in the absence or presence of combinational immunosuppressive nanofibers (0.05 µg/well of nanofiber-conjugated PD-L1-Ig, 0.05 µg/well of nanofiber-conjugated CD86-Ig, 0.1 µg/well of nanofiber-loaded leflunomide and 0.5 µg/well of nanofiber-bound TGF-β1 mimetic peptide), colon extracellular matrix (5 µg/well), or their combination were determined by quantifying the calcein retained in the colon epithelial cells 48 h after co-treatment according to the manufacturer’s protocol. The cell killing of each treatment group was calculated according to a published method^76^.

## In vivo studies

Animals were maintained in the Animal Resource Center at UTSW. All procedures involving the experimental animals were performed following the protocol approved by UTSW Institutional Animal Care and Use Committee and conformed to the *Guide for the Care and Use of Laboratory Animals* (NIH publication no. 86-23, revised 1985). Wild-type C57BL/6 mice (nomenclature: C57BL/6NCrl) were purchased from Charles River Laboratories, Inc. (Houston, TX). IL10 KO mice (B6 background, nomenclature: B6.129P2-Il10tm1Cgn/J) and wild-type C57BL/6 mice (also known as B6 WT, nomenclature: C57BL/6J) were purchased from the Jackson Laboratory for the IL10 KO mouse study. Mice were randomized, ear tagged, and grouped on arrival. Grouped mice were housed in the conventional non-sterilized mouse facility for at least 1 week before all *in vivo* studies were conducted.

### Induction and assessment of colitis

Three mouse colitis models were used in this study. For the dextran sodium sulfate-induced model, colitis was induced according to an established protocol^53,54^. Mice were housed in groups of 4 or 5 and acclimatized for at least 7 days before the study. To induce colitis, C57BL/6 mice (7 to 8 weeks old) were fed 3% dextran sodium sulfate (molecular weight c.a. 40,000; catalog number: J63606.22; Thermo Scientific) supplemented with deionized water (catalog number: W12-4; Fisher Scientific) for 6 days. Mouse cages were changed before and after the induction of colitis. Unless specified, mouse cages were changed weekly afterward. The bodyweight and disease activity index scores were monitored before the induction of colitis and then daily from day 4 after the induction of colitis.

For the trinitrophenol-induced model, colitis was induced according to the published protocol^52^. Mice were housed in groups of 4 or 5 and acclimatized for at least 7 days before the study. To induce colitis, mice received two intrarectal injections of 100 µL of 2.5% (w/v) trinitrobenzene sulfonate in 1:1 water/ethanol at day 0 (sensitization) and day 14 (colitis induction). Mice were starved for 18 h prior to each intrarectal injection. Mouse cages were changed weekly or the day after the intrarectal injection. The bodyweight and disease activity index scores for the trinitrophenol-induced colitis mice were monitored every two days from day 0 and daily from day 13 of the study.

IL10 KO mice spontaneously develop colitis^55^. 4-week s old IL10 KO mice and B6 WT (C57BL/6, wide-type control) mice were purchased from the Jackson Laboratory. All mice were born within 3 days. Mice were housed in groups of 4 or 5 and were ear tagged for identification. Mouse cages were changed weekly. Once onset, colitis mice house with colitis mice. The bodyweight and disease activity index scores were monitored every 2 or 3 days after the study began. Whole blood (≈ 100 µL) was collected from the facial vein at 7, 9, 11, 13, 15 and 17 weeks of age for complete blood count. IL10 KO mice that showed colitis symptoms at 10 – 11 weeks of age were used in this study.

The disease activity index score was calculated by combining the score of percentage of bodyweight loss (0 = no weight loss from baseline, 1 = less than 5% weight loss, 2 = 5-10% weight loss, 3 = 10-20% weight loss, and 4 = more than 20% weight loss), diarrhea (0 = normal stool, 2 = semi-formed stool, 4 = liquid that adheres to the anus), and rectal bleeding (0 = no bleeding, 2 = positive hemoccult test, and 4 = gross bleeding). For statistical analysis, the disease activity index scores and bodyweights of dextran sodium sulfate colitis mice were compared based on data recorded up to day 10 (4 days after the therapeutic treatment). Unless specified, DSS colitis mice were euthanized at days 12-14, and colons were preserved by trained veterinary technicians at the UTSW ARC Diagnostic Laboratory without prior knowledge of the treatment. For the trinitrophenol-induced colitis mice, disease activity index scores were compared daily from day 16 to day 20. trinitrophenol-induced colitis mice were euthanized at day 20, and colons were preserved by trained veterinary technicians at the UTSW ARC Diagnostic Laboratory without prior knowledge of the treatment. For the IL10 KO mice, disease activity index scores were compared based on data collected up to 17 weeks of age (endpoint). IL10 KO mice were euthanized at 17 weeks of age, and colons were preserved by trained veterinary technicians at the UTSW ARC Diagnostic Laboratory without prior knowledge of the treatment. Unless specified, the lengths of preserved colons were measured, and fixed with 10% NBF for 24 h. Fixed specimens were kept in 70% ethanol and transferred to the UTSW Histo Pathology Core or IDEXX BioAnalytics (Columbia, MO) for paraffin embedding before further histological studies.

### *In vivo* evaluation of different immune niche formulations for amelioration of dextran sodium sulfate-induced colitis in a mouse model

Therapeutic colitis treatment was performed at day 6 after the induction of colitis. Colon-specific immune niches were engineered by gently mixing the combination of immunosuppressive nanofibers (1 mg/mouse PD-L1/CD86 dual-functionalized nanofibers, 1 mg/mouse leflunomide-encapsulated nanofibers and/or 1 mg/mouse TGF-β1 mimetic peptide-encapsulated nanofibers in 1X PBS), ball-milled decellularized colon extracellular matrix (0.5 mg/mouse in 1X PBS), and colon epithelial cells (1×10^6^ cells/mouse, in 1X PBS) together prior to conducting the *in vivo* studies. Suspensions of colon-specific immune niches (200 µL per mouse) were s.c. inoculated into the neck fat pocket *via* a 1 mL syringe equipped with a 21-gauge needle. In the control studies, dextran sodium sulfate-induced colitis mice received subcutaneous treatment with: (i) colon epithelial cells (1×10^6^ cells/mouse), colon (0.5 mg/mouse) and isotype control nanofibers (1 mg/mouse of IgG isotype control nanofibers, 1 mg/mouse of drug-free azide poly(ethylene glycol)-block-poly(lactide-co-glycolide) nanoparticle-functionalized nanofibers and 1 mg/mouse non-functionalized NFs); (ii) colon epithelial cells (1×10^6^ cells/mouse) and combinational immunosuppressive nanofibers (1 mg/mouse of PD- L1/CD86-functionalized nanofibers, 1 mg/mouse of leflunomide-encapsulated nanofibers and 1 mg/mouse TGF-β1 mimetic peptide-encapsulated nanofibers); (iii) colon extracellular matrix (0.5 mg/mouse); (iv) colon extracellular matrix (2 mg/mouse); (v) free immunosuppressants (5 µg of PD-L1-Ig, 5 µg of CD86-Ig, 120 µg/mouse of leflunomide-encapsulated nanoparticles and 50 µg of TGF-β1 mimetic peptide); (vi) MIM6 cells (1×10^6^ cells/mouse), pancreas extracellular matrix (0.5 mg/mouse) and combinational immunosuppressive nanofibers (1 mg/mouse of PD-L1/CD86- functionalized nanofibers, 1 mg/mouse of leflunomide-encapsulated nanofibers and 1 mg/mouse TGF-β1 mimetic peptide-encapsulated nanofibers). In another control study, DSS colitis mice received an i.p. treatment with combinational nanofibers (1 mg/mouse PD-L1/CD86- functionalized nanofibers, 1 mg/mouse leflunomide-encapsulated nanofibers and 1 mg/mouse TGF-β1 mimetic peptide-encapsulated nanofibers). The therapeutic effect was compared with that of subcutaneous treatment with combinational colon-specific immune niche.

Unless specified, dextran sodium sulfate-induced colitis mice were euthanized on days 13- 14, colons were preserved by trained veterinary technicians at the UTSW ARC Diagnostic Laboratory without prior knowledge of the treatment. Unless specified, the lengths of preserved colons were measured, and fixed with 10% NBF for 24 h. Fixed specimens were kept in 70% ethanol and transferred to the UTSW Histo Pathology Core or IDEXX BioAnalytics (Columbia, MO) for paraffin embedding before further histological studies. Subsequent H&E staining was performed by UTSW Histo Pathology Core or IDEXX BioAnalytics (Columbia, MO). Ki67 and TUNEL staining was performed by UTSW Histo Pathology Core. Further immunohistological studies were performed by the Pathology Services Core in the UNC Lineberger Comprehensive Cancer Center at the UNC School of Medicine or iHisto, Inc. (Salem, MA). Colon damage in each mouse was assessed from representative H&E-stained colon sections as previously reported^79^. Briefly, colon epithelial damage was assigned scores as follows: 0 = normal; 1 = hyperproliferation, irregular crypts; 2 = mild to moderate crypt loss (10–50%); 3 = severe crypt loss (50–90%); 4 = complete crypt loss, surface epithelium intact; 5 = small to medium size ulcer (<10 crypt widths); and 6 = large ulcer (>10 crypt widths). An inflammatory cell infiltration score was assigned separately for the mucosa (0 = normal, 1 = mild, 2 = modest, 3 = severe), submucosa (0 = normal, 1 = mild to modest, 2 = severe), and muscle/serosa (0 = normal, 1 = moderate to severe). Scores for epithelial damage and inflammatory cell infiltration were added, resulting in a total score ranging from 0 to 12. For selected experimental groups, feces were collected from mice 5 days after colitis treatment before euthanization. Preserved feces were submitted to the UTSW Microbiome Research Lab for 16S rRNA sequencing. Preserved colons were homogenized in (i) TrisHCl/Triton X-100-based lysis buffer^79^ in the presence of Thermo Scientific^TM^ Halt^TM^ Protease Inhibitor Cocktail (1X; ThermoFisher) for the quantification of mouse IL-6, mouse TNF-alpha, mouse IL-10, and mouse TGF-β1 expression in the preserved colon tissues in the Genomic and Microarray Core Facility at UTSW or (ii) hexadecyltrimethylammonium bromide buffer^79^ for the quantification of MPO expression through a Cayman Chemical Neutrophil Myeloperoxidase Activity Assay Kit according to the manufacturer’s protocol.

For the immune cell depletion study, three doses of anti-mouse CD8a antibody (clone: YTS169.4; catalog number: BE0117; BioXCell) and anti-mouse CD25 antibody (clone: PC- 61.5.3; catalog number: BE0012; BioXCell) were intraperitoneal administered on days 4, 6, and 8 to deplete CD8^+^ T cells and regulatory T cells, respectively. The antibody dose was 200 µg/mouse per injection. A single dose of clodrosome (140 µL, containing 700 µg of encapsulated clodronate; Encapsula NanoSciences, Brentwood, TN) was intravenous tail vein-administered at day 5 to deplete macrophages.

The chronic dextran sodium sulfate-induced colitis study was performed as previously reported^53^. Mice were subjected to 4 cycles of 3% dextran sodium sulfate-induced treatment for 5 to 6 days, followed by 14 days of normal drinking water. Mice in the single treatment group received a single subcutaneous treatment with the combo immune niche at day 6. Mice in the multiple (triple) treatment group received subcutaneous treatment of combinational colon-specific immune niche at days 6, 25, and 44. Each subcutaneous injection was composed of 1 mg of PD- L1/CD86-functionalized nanofibers, 1 mg of leflunomide-encapsulated nanofibers, 1 mg of TGF- β1 mimetic peptide-encapsulated nanofibers, 0.5 mg of decellularized colon extracellular matrix, and 1×10^6^ colon epithelial cells. Mice were euthanized at day 48. Colons were preserved for further histological study.

An *ex vivo* biodistribution study was performed to investigate the biodistribution of subcutaneously-administered combinational niche and intraperitoneal-administered combinational nanofibers in dextran sodium sulfate-induced colitis mice *via ex vivo* fluorescence imaging. Similar to the therapeutic study, rhodamine-labeled combinational niche (constructed from rhodamine-labeled combinational nanofibers) and rhodamine-labeled combinational nanofibers were administered to dextran sodium sulfate-induced colitis mice at day 6 after the initial study. Mice were euthanized by an overdose of carbon dioxide 3 days after colitis treatment (day 9 after the initial study). The biodistribution of rhodamine-labeled nanofibers was quantified by an AMI HTX optical imaging system (Spectra Instrument Imaging). Selected organs were preserved to quantify the uptake of rhodamine-labeled nanofibers.

Immune profiling studies were performed on colon and subcutaneous immune niches preserved 3 (for immune niche), 7 (for colon and immune niche), and 12 days (for colon) after colitis treatment *via* mass cytometry. For the preserved colons, immune cells infiltrated into lamina propria cells were isolated as previously reported^56^. Freshly isolated cells were transferred to the University Flow and Mass Cytometry Facility at UTSW for staining and subsequent flow mass cytometry study. For the preserved subcutaneous-inoculated combinational niche, the preserved niches were fixed for 24 h before transfer to the UTSW Histo Pathology Core and University Flow and Mass Cytometry Facility at UTSW for imaging mass cytometry study. The antibodies used in the mass cytometry studies are summarized in Supplementary Table 5.

*In vivo* evaluation of trinitrophenol-modified combinational colon-specific immune niche for amelioration of trinitrohpehnol-induced colitis in a mouse model. To demonstrate that the subcutaneously inoculated combinational niche can ameliorate trinitrophenol-induced colitis, therapeutic treatment was performed 1 day after the second intrarectal injection of trinitrophenol (15 days after the initial study). Briefly, trinitrophenol-modified combinational niche composed of trinitrophenol-modified colon epithelial cells (1x10^6^ cells per mouse), colon extracellular matrix (0.5 mg per mouse) and combinational niche (containing 1 mg/mouse PD-L1/CD86-functionalized nanofibers, 1 mg/mouse leflunomide-encapsulated nanofibers and 1 mg/mouse TGF-β1 mimetic peptide-encapsulated nanofibers) was subcutaneously administered to the neck fat pocket of the trinitrophenol-induced colitis mice. Mice were monitored daily after the treatment. Mice were euthanized on day 20. Colons were preserved by trained veterinary technicians at the UTSW ARC Diagnostic Laboratory without prior knowledge of the treatment. The lengths of preserved colons were measured, and fixed with 10% NBF for 24 h. Fixed specimens were kept in 70% ethanol and transferred to the UTSW Histo Pathology Core for histological study.

### *In vivo* evaluation of combinational colon-specific immune niche for amelioration of colitis in IL10 KO mice

To demonstrate that s.c administration of combinational niche can ameliorate colitis in IL10 KO mice that spontaneously develop colitis, IL10 KO mice that developed colitis at 10 – 11 weeks of age received repeated colitis treatment with combinational niche at 11.5, 13.5 and 15.5 weeks of age. Each treatment involved s.c. administration of combinational niche engineered from colon epithelial cells (1x10^6^ cells per mouse), colon extracellular matrix (0.5 mg per mouse) and combinational nanofibers (containing 1 mg/mouse PD-L1/CD86 NFs, 1 mg/mouse leflunomide-encapsulated nanofibers and 1 mg/mouse TGF-β1 mimetic peptide-encapsulated nanofibers). Bodyweight and disease activity index scores were monitored every 2 or 3 days after treatment. Whole blood samples were collected every 2 weeks for complete blood count at the UTSW ARC Diagnostic Laboratory. Mice were euthanized at 17 weeks of age. Colons were preserved by trained veterinary technicians at the UTSW ARC Diagnostic Laboratory without prior knowledge of the treatment. The lengths of preserved colons were measured, and fixed with 10% NBF for 24 h. Fixed specimens were kept in 70% ethanol and transferred to the UTSW Histo Pathology Core for histological study.

### *In vivo* evaluation of immune niches for ameliorating colitis-associated colorectal cancer in an azoxymethane/dextran sodium sulfate model of colitis-associated cancer

The ability of the immunosuppressive immune niche to ameliorate colitis-associated colorectal cancer was evaluated in the established azoxymethane/dextran sodium sulfate colitis mouse model as previously reported^62, 63^. At the beginning of the study, azoxymethane (Sigma) pre-dissolved in 1X PBS (Gibco) was intraperitoneal administered to a healthy group of C57BL/6 mice (female, 8 to 9 weeks old) at a dose of 10 mg/kg. Mice in the dextran sodium sulfate-induced colitis and azoxymethane/dextran sodium sulfate colitis groups were fed 3% dextran sodium sulfate drinking water at days 7, 27, and 47 for 6 days. Mice in the treatment group received subcutaneous therapeutic treatment with combinational niche at days 13, 32, and 52. Each immune niche treatment was composed of 1 mg of PD-L1/CD86-functionalized nanofibers, 1 mg of leflunomide-encapsulated nanofibers, 1 mg of TGF-β1 mimetic peptide-encapsulated nanofibers, 0.5 mg of ball-milled decellularized colon extracellular matrix, and 1×10^6^ colon epithelial cells. Bodyweight and disease activity index scores were monitored every 1 to 2 days for up to 65 days. Mice were then euthanized. Colons were preserved and washed with saline before being stained with 1% Alcian blue (Electron Microscopy Sciences) to label tumor lesions. The length of the colon was measured, and the number of tumors was counted. The colon was then washed 2 to 3 times with saline to remove the Alcian blue before being fixed with 10% NBF for further histological study at the Histo Pathology Core at UTSW.

### *In vivo* immune-mediated colitis treatment in syngeneic tumor models

B16F10, B16OVA, and MC38 xenograft tumors were inoculated into healthy C57BL/6 mice (female, 7 to 8 weeks old) through s.c. inoculation of B16F10 cells (2.5×10^5^ cells/mouse) in serum-free DMEM, B16OVA cells (2.5×10^5^ cells/mouse) in serum-free RPMI 1640 culture medium, and MC38 cells (5×10^5^ cells/mouse) in serum-free DMEM in the right flank. Tumor volume (volume = 0.5×a×b^2^, where *a* is the large diameter and *b* is the small diameter) was measured every 2 to 3 days from day 7 after the tumor was inoculated.

For the copanlisib-triggered colitis study in the B16OVA tumor model, copanlisib treatment was provided by intraperitoneal administration of copanlisib (25 mg/kg, in saline) at days 6, 8, 10, 12, 14, 16 and 18 after tumor inoculation. Colitis was induced by feeding the tumor-bearing mice 3% dextran sodium sulfate drinking water at day 14 for 4 days. For the colitis treatment group, mice received a single s.c. treatment with combinational colon-specific immune niche at day 18. Bodyweight and disease activity index scores were monitored every 2 or 3 days. Mice were euthanized at 22 days post-inoculation. Colons were preserved by trained veterinary technicians at the UTSW ARC Diagnostic Laboratory without prior knowledge of the treatment. The lengths of preserved colons were measured, and fixed with 10% NBF for 24 h. Fixed specimens were kept in 70% ethanol and transferred to the UTSW Histo Pathology Core for histological study (H&E, Ki67 and TUNEL stains).

For the B16F10 and B16OVA tumor models, immune checkpoint blockade therapy was provided by i.p. administration of 100 µg of anti-mouse CTLA-4 antibody (clone: 9H10; catalog number: BE0131; BioXCell) and 100 µg of anti-mouse PD-1 antibody (clone: RMP1-14; catalog number: BP0146; BioXCell) at days 7 and 9 after the tumor was inoculated. For the MC38 tumor model, mice received immune checkpoint blockade treatments at days 7 and 9, followed by maintenance treatments at days 19, 22, and 25. Colitis was induced by feeding the tumor-bearing mice 3% dextran sodium sulfate drinking water at day 7 for 6 days. For the colitis treatment group, mice received a single subcutaneous treatment with combinational niche or intraperitoneal treatment with combinational nanofibers at day 13. Each immune niche treatment was composed of 1 mg of PD-L1/CD86-functionalized nanofibers, 1 mg of leflunomide-encapsulated nanofibers, 1 mg of TGF-β1 mimetic peptide-encapsulated nanofibers, 0.5 mg of ball-milled decellularized colon extracellular matrix, and 1×10^6^ colon epithelial cells.

For the B16F10 tumor model, mice were euthanized when the large diameter of the tumor was more than 2 cm, if they had suffered 10% weight loss in 2 days, or 21 days after tumor inoculation. Colons were preserved for further histological study. For the B16OVA tumor model, mice were euthanized at day 16 (3 days after colitis treatment). Xenograft tumors were preserved for mechanistic study to investigate the phenotypes of the tumor-infiltrated T cells. Briefly, a preserved tumor was digested with 2 mg/mL collagenase D (Roche) and 0.2 mg/mL DNase I (Roche) in HBBS buffer (10 mL per tumor). The dead cells were first stained with Live/Dead Fixable Green Dead Cell Stain Kit (catalog number: L23101; Invitrogen) before being incubated with PE-labeled OVA(257-264)-specific tetramer (catalog number: TB50011; MBL International Corporation) at 20 °C for 1 h. The stained cells were then divided into two portions before being stained with (i) PE/Cy5-labeled anti-mouse CD3 antibody (clone: 17A2; catalog number: 100274; BioLegend) and APC-labeled anti-mouse CD8a antibody (clone: 53-6.7; catalog number: 100712; BioLegend) or (ii) APC-labeled anti-mouse CD8a antibody (clone: 53-6.7; catalog number: 100712; BioLegend) and PE/Cy5-labeled anti-TIM-3 antibody (clone: F38-2E2; catalog number: 345052; BioLegend). Stained cells were analyzed in a Calibur flow cytometer at the University Flow and Mass Cytometry Facility at UTSW. For the MC38 tumor model, mice were euthanized when the large diameter of the tumor was greater than 2 cm, if they suffered 10% weight loss in 2 days, or 42 days after the tumor inoculation.

In a control study, B16F10 tumor-bearing mice received immune checkpoint blockade treatments at days 7 and 9 and were fed 3% dextran sodium sulfate drinking water from days 7 to 13. Mice were euthanized at day 11 or 13. Spleens were preserved. The splenocytes were then stained with FITC-labeled PD-L1-Ig, rhodamine-labeled CD86, and APC-labeled anti-mouse CD8a antibody (clone: 53-6.7; catalog number: 100712; BioLegend) to confirm that the CD8^+^ T cells can bind to PD-L1 and CD86. The stained cells were analyzed in a Calibur flow cytometer at the University Flow and Mass Cytometry Facility at UTSW.

### Statistical analysis

No statistical methods were used to pre-determine the sample size of the experiment. The sample sizes were based on similar published *in vitro* and *in vivo* studies. No collected experimental data were excluded from the quantitative analysis. Quantitative data are expressed as the mean ± standard error of the mean (SEM). One-way or two-way analysis of variance (ANOVA), followed by Tukey’s honestly significant difference (HSD) multiple comparison post hoc test was used to test differences among groups. Statistical analyses were performed in Graph Pad Prism 6 software pack. A p-value < 0.05 was considered statistically significant and a p-value > 0.05 was considered statistically insignificant.

### Reporting Summary

Further information on the research design is available in the Nature Research Summary linked to this article.

## Data availability

The data supporting the findings of this study are available within the article and its Supplementary Information files. All relevant data can be provided by the authors upon reasonable request.

## Extended Data

### Supplementary Figures

### Supplementary Tables

**Supplementary Fig. 1.**
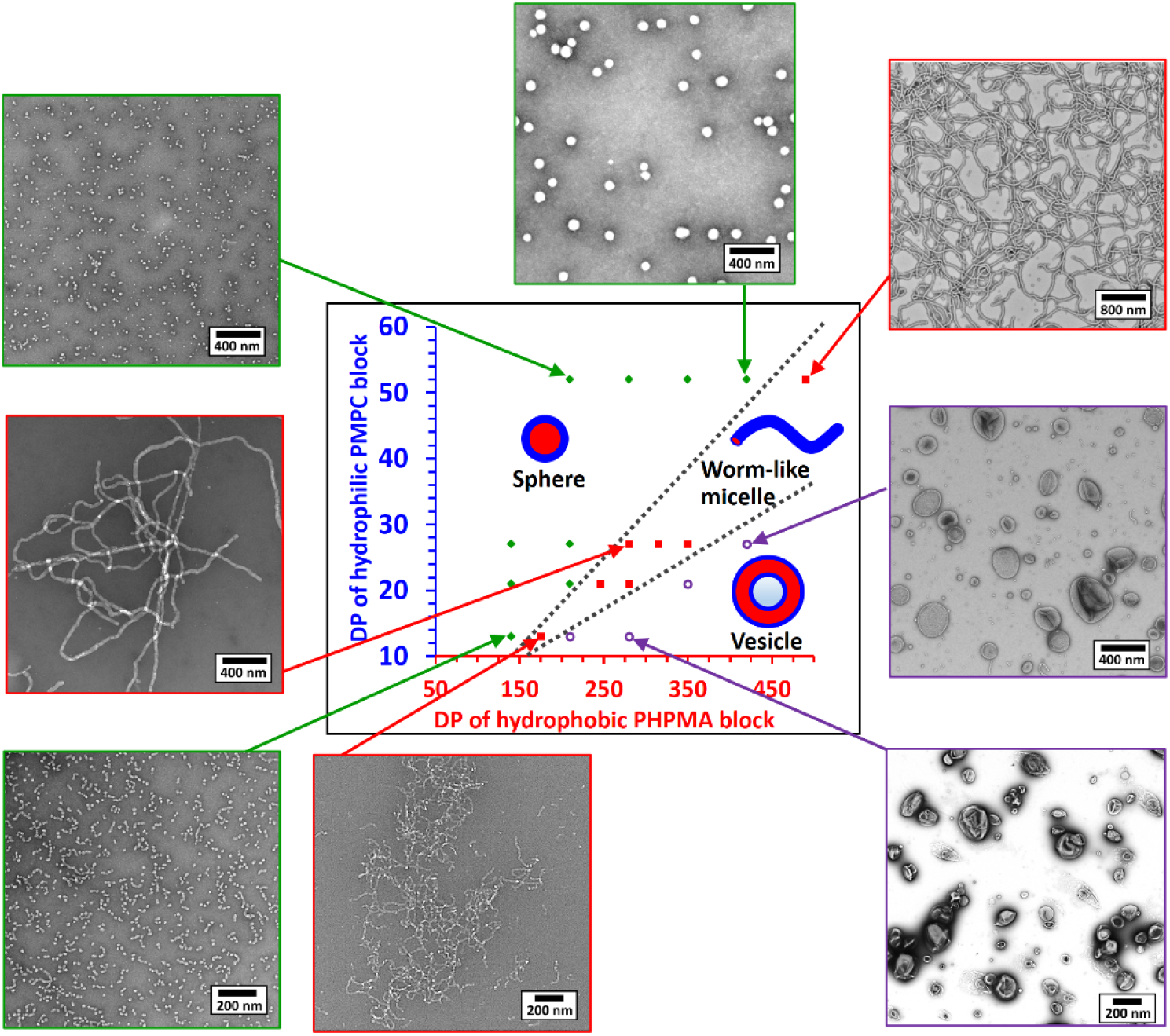
Self-assembly of poly(2-(methacryloyloxy)ethyl phosphorylcholine)- poly(2-hydroxypropyl methacrylate) diblock copolymers. Phase diagram of PMPC-PHPMA diblock copolymers with different poly(2-(methacryloyloxy)ethyl phosphorylcholine) and poly(2- hydroxypropyl methacrylate) block lengths. All diblock copolymers were prepared *via* aqueous reversible addition-fragmentation chain transfer polymerization at 25 wt/wt% solid content. High solid content facilitates the diblock copolymers self-assembly into different nano-objects. PMPC, poly(2-(methacryloyloxy)ethyl phosphorylcholine); PHMPA, poly(2-hydroxypropyl methacrylate).

**Supplementary Fig. 2.**
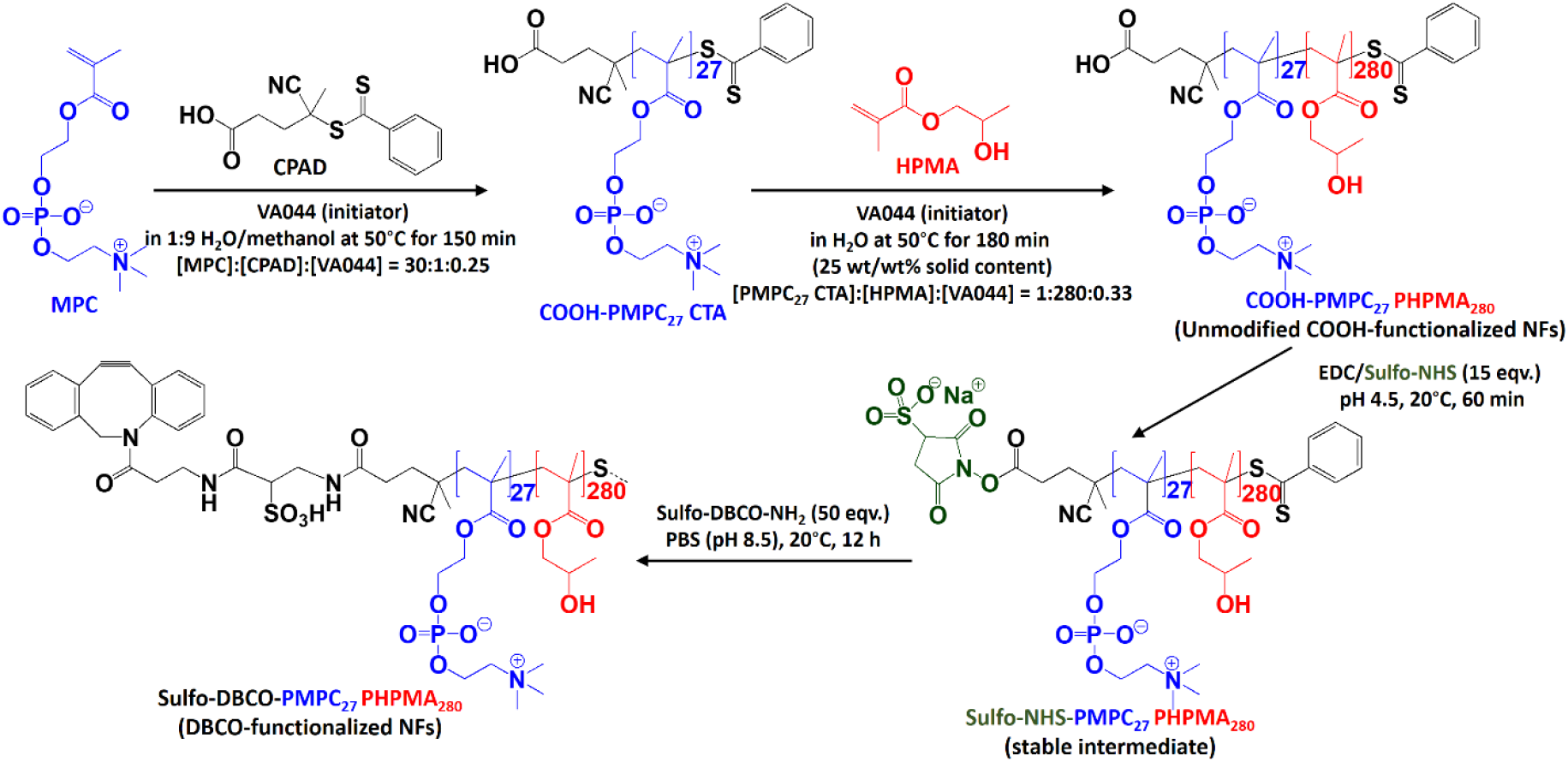
Synthesis of poly(2-(methacryloyloxy)ethyl phosphorylcholine)- poly(2-hydroxypropyl methacrylate) diblock copolymer nanofibers through an optimized two-step reversible addition-fragmentation chain transfer polymerization. Carboxylic acid α- end-functionalized PMPC27 macro-chain transfer agent (macro-CTA) synthesized through reversible addition-fragmentation chain transfer polymerization. Chain extension of the PMPC27 macro-chain transfer agent with HPMA at 25 wt/wt% solid content yielded worm-like PMPC27- PHPMA280 (carboxylic acid (COOH)-functionalized) nanofibers. The as-prepared carboxylic acid-functionalized nanofibers were then activated with EDC/sulfo-NHS before being functionalized with sulfo-dibenzocyclooctyne amine (sulfo-DBCO amine) to produce DBCO-functionalized nanofibers. MPC = 2-methacryloyloxyethyl phosphorylcholine; CPAD = 4-cyano-4-(phenylcarbonothioylthio) pentanoic acid (chain transfer agent); VA044 = 2,2’-azobis [2-(2- imidazolin-2-yl) propane] dihydrochloride (thermal initiator); HPMPA = 2-hydroxypropyl methacrylate (monomer); EDC = 1-ethyl-3-(3-dimethylaminopropyl) carbodiimide; sulfo-NHS = N-hydroxysulfosuccinimide. A family of PMPC27-PHPMAx (where x = 140, 210, 350, and 420) diblock polymers were prepared using the same method, except that different PMPC27 macro-chain transfer agent and HPMA (monomer) molar ratios were used while the solid content was maintained at 25 wt/wt%.

**Supplementary Fig. 3.**
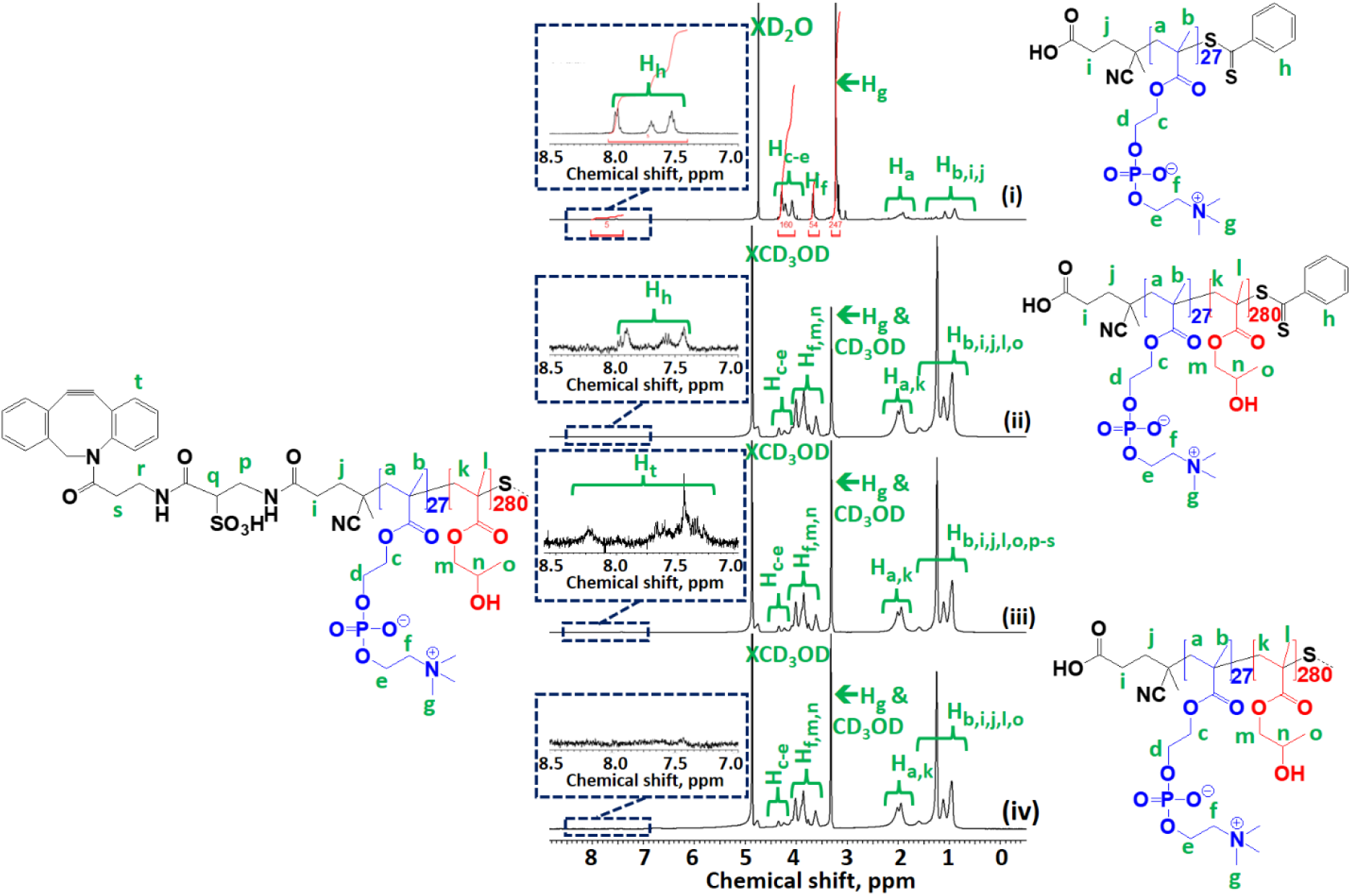
^1^H NMR spectroscopy of purified diblock copolymer nanofibers. ^1^H NMR (400 MHz, in CD3OD) spectra of (i) purified PMPC27 macro-chain transfer agent; (ii) unmodified COOH-functionalized PMPC27-PHPMA280 diblock copolymer; (iii) purified DBCO-functionalized PMPC27-PHPMA280 diblock copolymer; and (iv) unmodified COOH-functionalized PMPC27-PHPMA280 diblock copolymer after adjusted to pH 8.5 (in 1X PBS) and dialysis against water. The COOH-functionalized PMPC27-PHPMA280 diblock copolymer was lyophilized before being redissolved in CD3OD for the ^1^H NMR study. The DBCO-functionalized PMPC27-PHPMA280 diblock copolymer was purified *via* equilibrium dialysis against deionized water and lyophilized before being redissolved in CD3OD for the ^1^H NMR study. The control sample, dialyzed unmodified COOH-functionalized PMPC27-PHPMA280 diblock copolymer, was first incubated in 1X PBS at pH 8.5 for 12 h before dialysis against deionized water. The control diblock polymer was lyophilized before being redissolved in CD3OD for the ^1^H NMR study. The thiocarbonylthio reversible addition-fragmentation chain transfer end group was hydrolyzed under basic conditions (pH 8.5) during amine-N-hydroxysulfosuccinimide coupling. Sulfo-NHS, N-hydroxysulfosuccinimide; EDC, 1-ethyl-3-(3-dimethylaminopropyl)carbodiimide.

**Supplementary Fig. 4.**
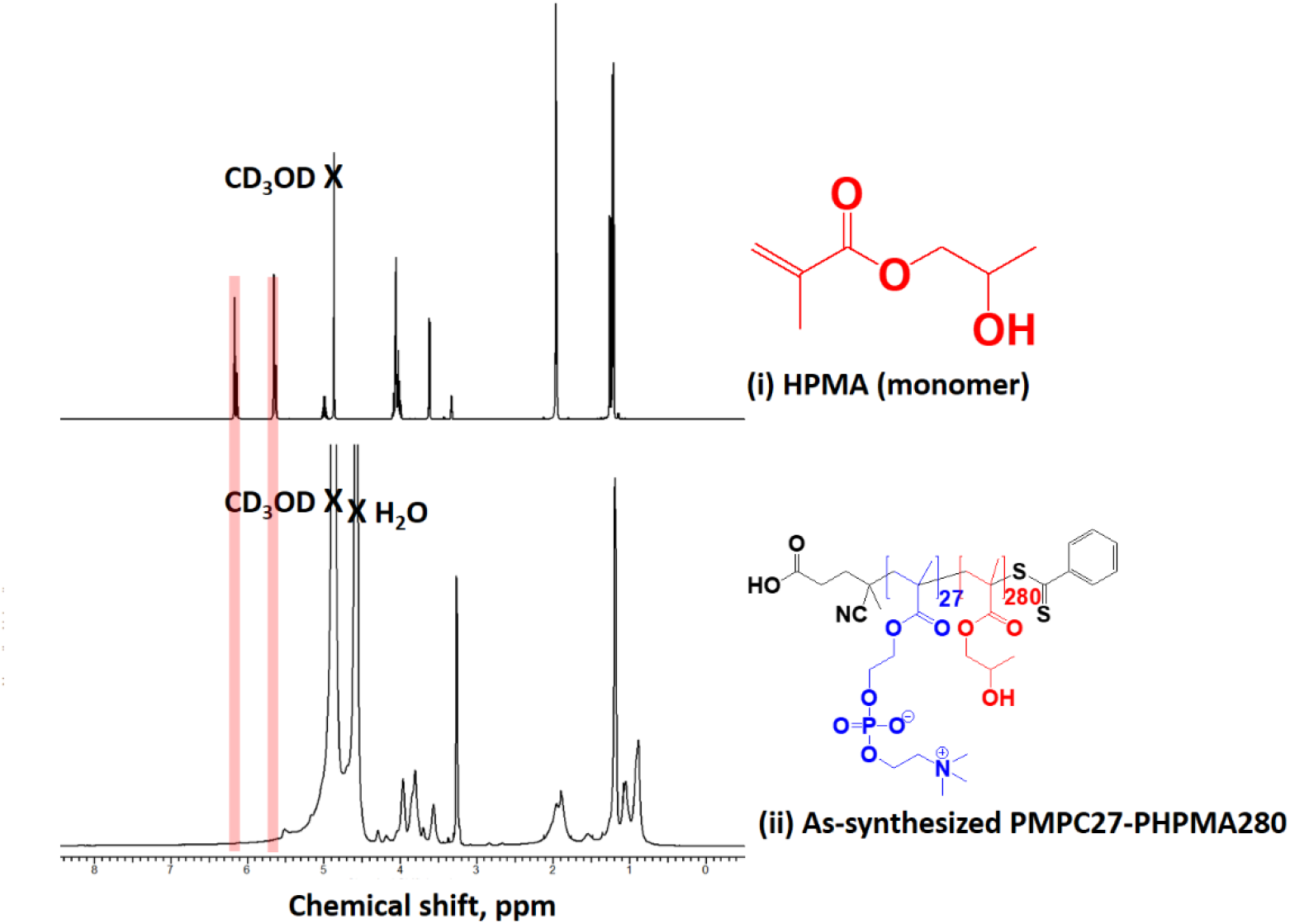
^1^H NMR spectroscopy of as-polymerized (non-purified) poly(2- (methacryloyloxy)ethyl phosphorylcholine)-poly(2-hydroxypropyl methacrylate) diblock copolymers. ^1^H NMR (400 MHz) spectra of (i) HPMA (monomer, recorded in CD3OD); and (ii) the as-synthesized PMPC27-PHPMA280 diblock copolymer. The diblock polymer was prepared at 25 wt/wt% solid content. The spectrum of the as-prepared diblock copolymer was recorded in 1:10 H2O/CD3OD. The disappearance of methacrylate protons (C<u>H2</u>=C(CH3)C(=O)O-R) at 5.7 and 6.2 ppm (highlighted red regions) in all diblock copolymers confirmed the complete polymerization of HPMA.

**Supplementary Fig. 5.**
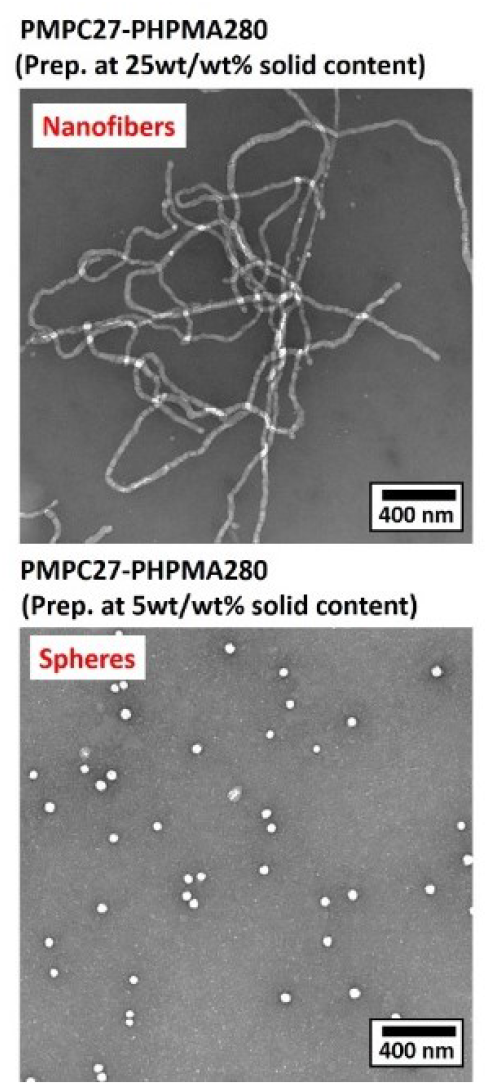
Representative transmission electron microscopy images of the PMPC27-PHPMA280 diblock copolymer prepared at different solid contents. The preparation of the PMPC27-PHPMA280 diblock polymer at 25 wt/wt % solid content yielded the formation of monophasic rod-like nanofibers and thus suggested further use for *in vitro* and *in vivo* studies.

**Supplementary Fig. 6.**
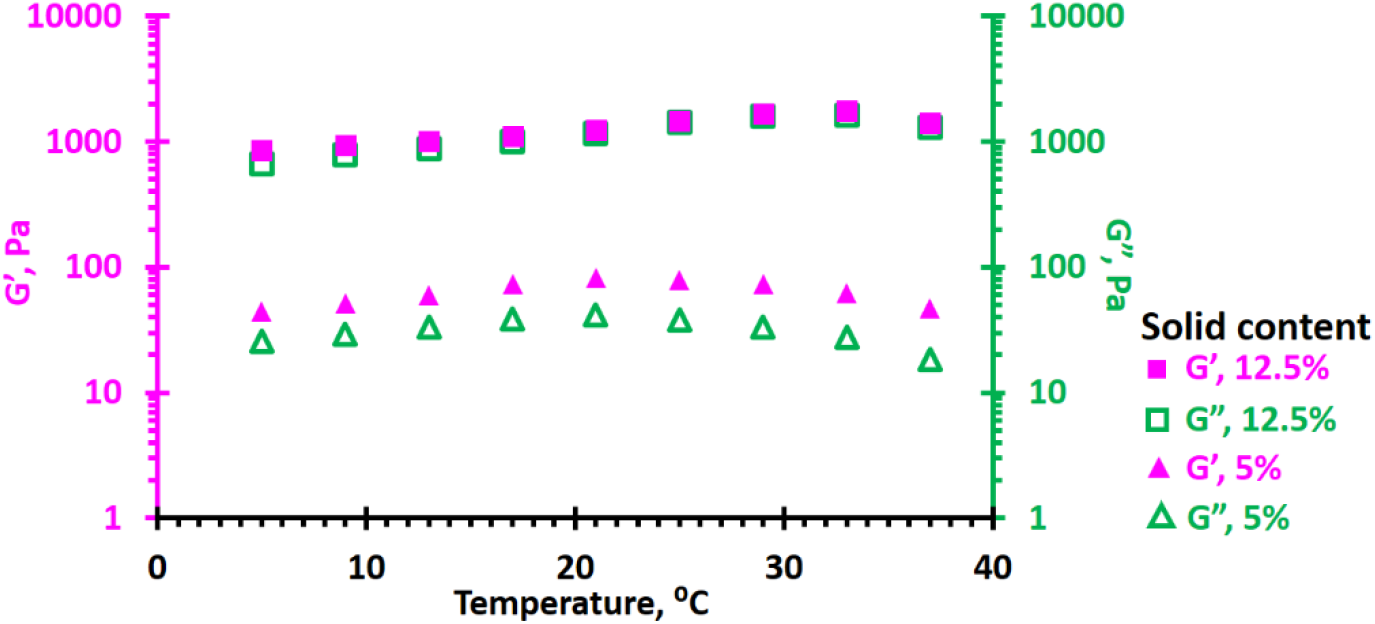
Physiological properties of PMPC27-PHPMA280 diblock copolymer nanofibers. Variable-temperature oscillatory rheology data were recorded for 12.5 wt/wt% and 5 wt/wt% of unmodified PMPC27-PHPMA280 diblock copolymer nanofibers. The oscillatory rheology study confirmed that the diblock copolymer is thermally stable between 5 °C and 37 °C.

**Supplementary Fig. 7.**
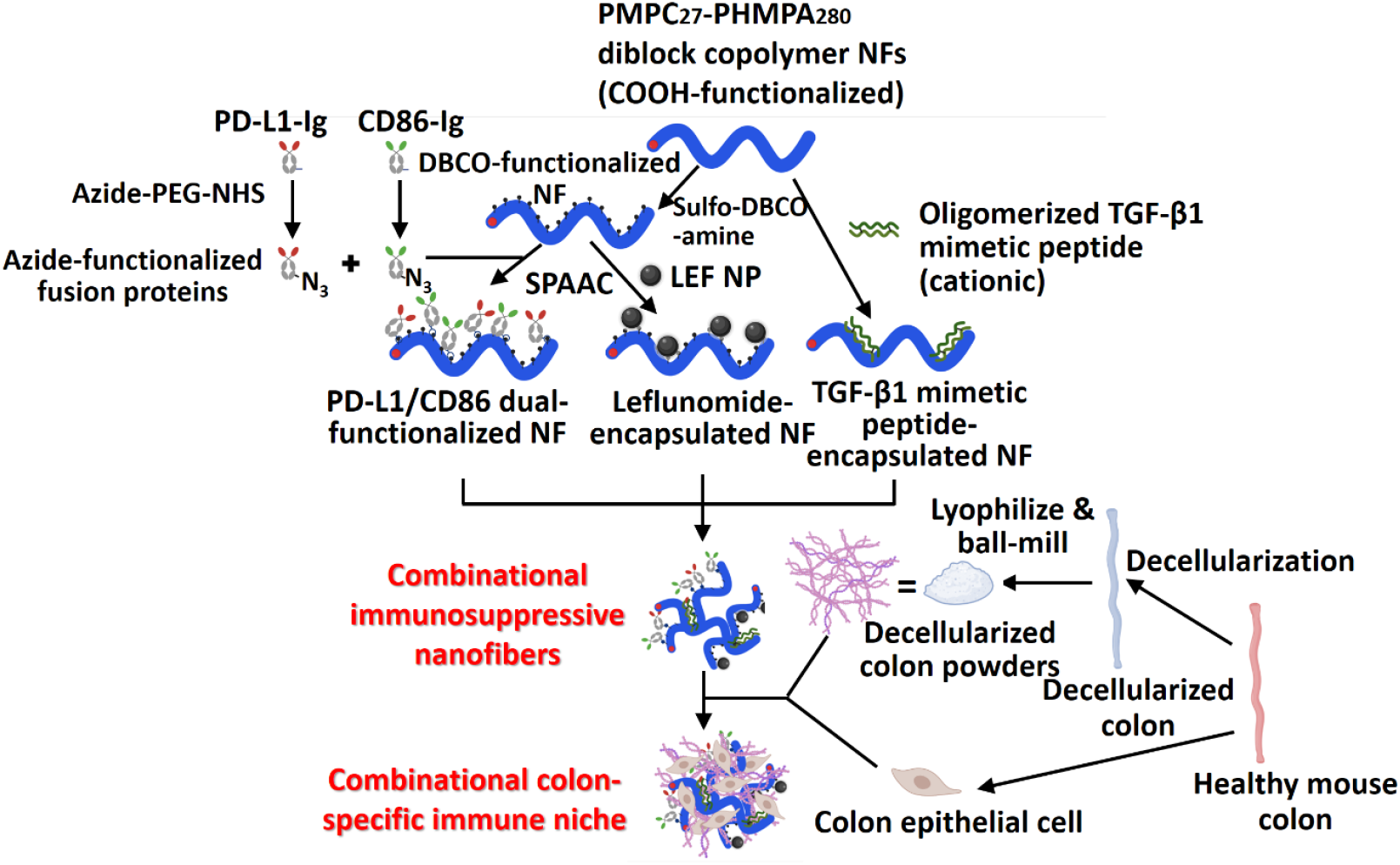
Bioengineering of combinational immunosuppressive nanofibers and combinational colon-specific immune niche. Bioengineering of the immune niche for ulcerative colitis treatment. The colon-specific immune niche was constructed from immunosuppressive nanofibers, colon epithelial cells, and a decellularized colon biomatrix. Different immunomodulator-functionalized nanofibers were constructed from COOH alpha-end-functionalized PMPC-PHPMA diblock copolymer nanofibers. COOH-functionalized nanofibers were converted into DBCO-functionalized nanofibers through sulfo-NHS–EDC coupling with sulfo DBCO-amine. PD-L1 immunoglobulin (PD-L1-Ig) and CD86 immunoglobulin (CD86-Ig) were converted into azide-functionalized PD-L1-Ig and CD86-Ig through amine-NHS ester coupling. The azide-functionalized fusion proteins were then conjugated to the DBCO-functionalized NFs through strain-promoted azide-alkyne cycloaddition. Azide-functionalized leflunomide-encapsulated nanoparticles were conjugated to DBCO-functionalized nanofibers *via* strain-promoted azide-alkyne cycloaddition. Cationic thiol-rich TGF-β1 mimetic peptide was first oligomerized by incubation at 20 °C for 4 h before physiosorbed onto the anionic COOH-functionalized NFs to give TGF-β1 mimetic peptide-encapsulated nanofibers. In this proof-of-concept study, decellularized colon biomatrix and colon epithelial cells were isolated from healthy C57BL/6 mice. In practice, healthy colon tissue and colon epithelial cells can be isolated from healthy colon tissue collected through biopsy or obtained from deceased donors. The decellularized colon was lyophilized and ball-milled to fine powder.

**Supplementary Fig. 8.**
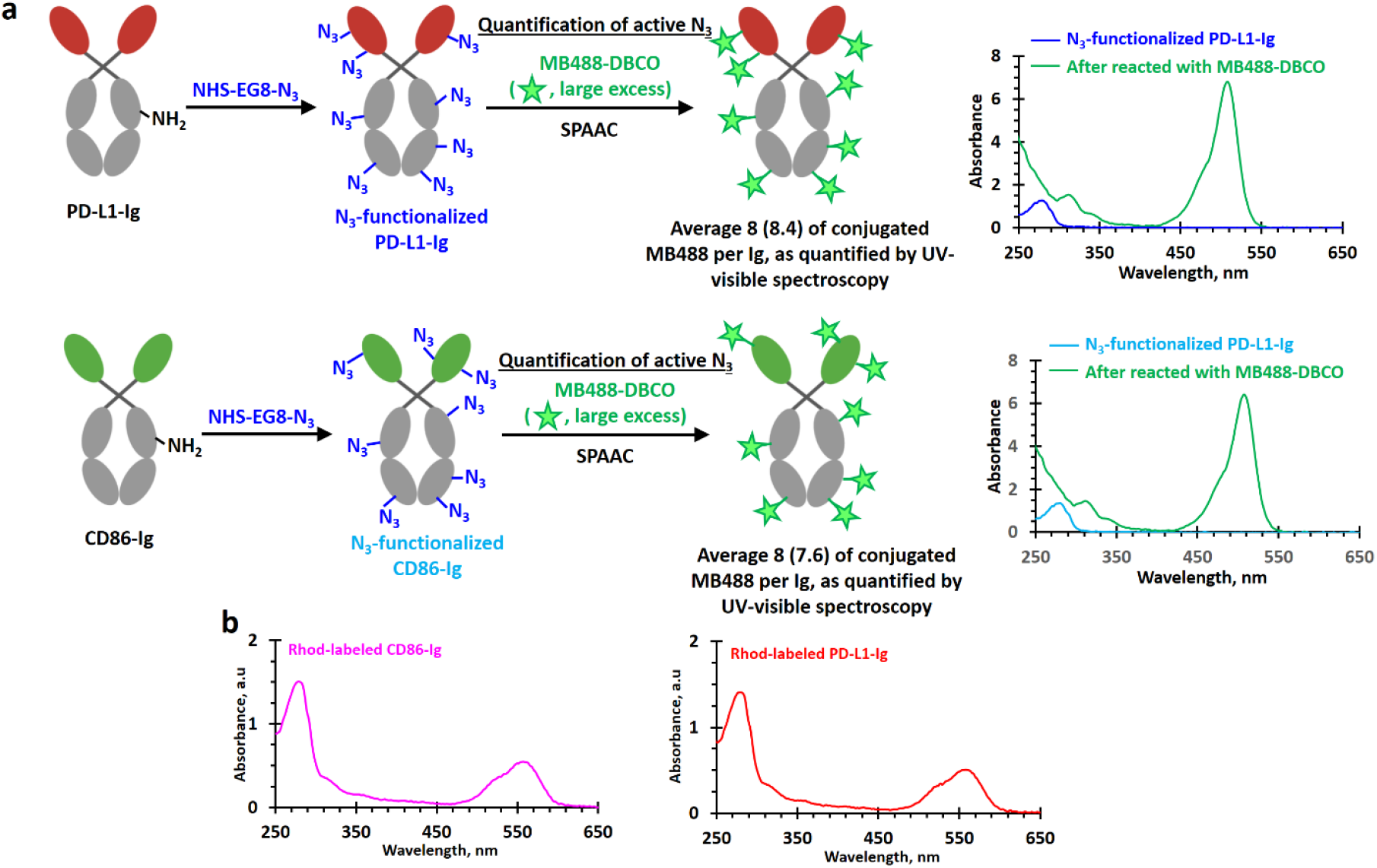
Characterization of azide (N3)-functionalized PD-L1-Ig and CD86-Ig. **a**, UV-visible spectra of 1 mg/mL azide-functionalized PD-L1-Ig and CD86-Ig. The number of conjugated azide ligands was quantified by reacting with a large excess (50 molar equivalents per Ig) of MB488-DBCO dye at 37 °C for 1 h. Unreacted MB488-DBCO was removed by desalting columns (3 times). The target degree of conjugation was 40, and the actual degree of conjugation was approximately 8 for both proteins. The azide ligands were randomly conjugated to both fusion proteins. **b,** UV-visible spectra of 1 mg/mL rhodamine B/azide dual-functionalized PD-L1-Ig and CD86-Ig. Both proteins were prepared using the same amine/NHS ester coupling (target degree of functionalization = 50) in the presence of 4 molar equivalents of rhodamine B isocyanate. It was calculated that each protein was labeled with an average of 1 rhodamine B molecule. SPAAC, strain-promoted azide-alkyne cycloaddition.

**Supplementary Fig. 9.**
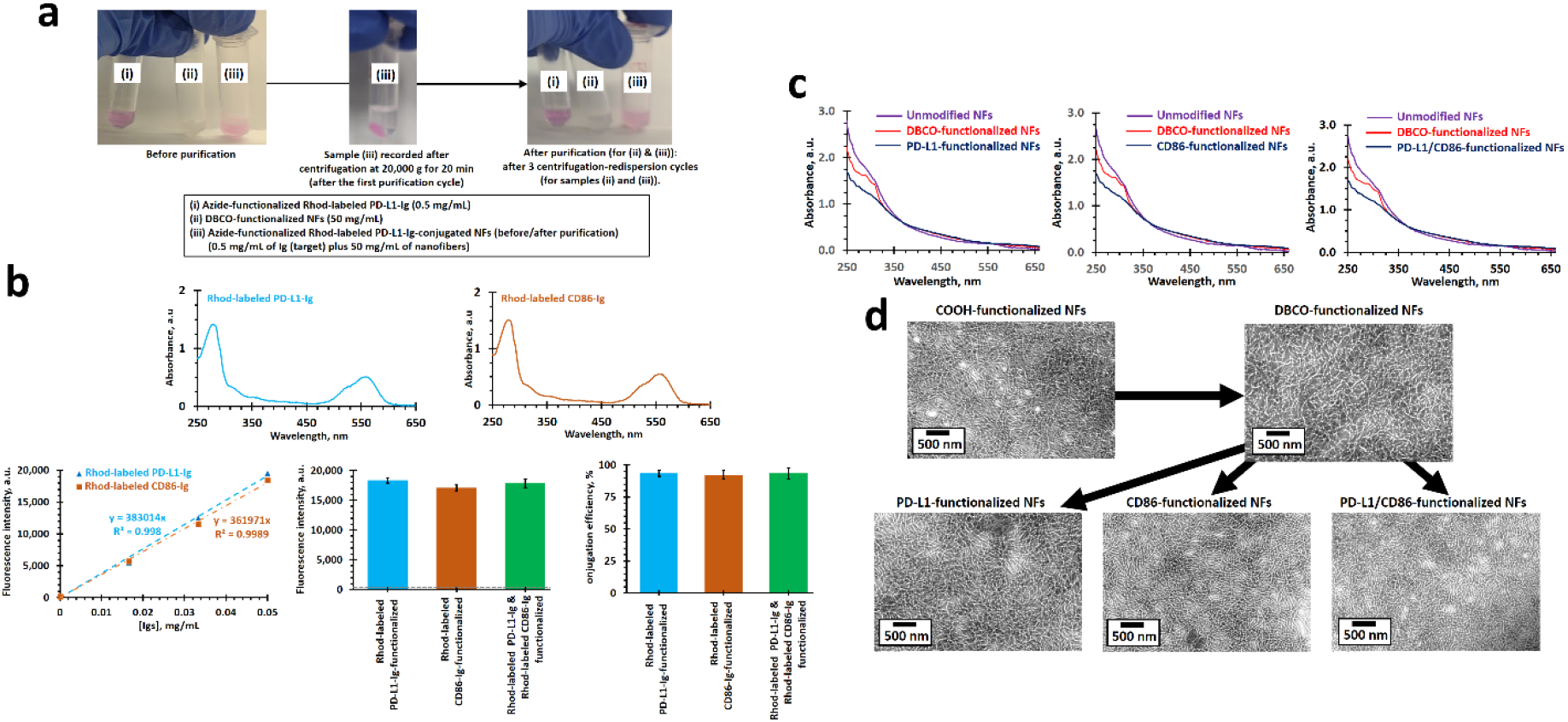
Characterization of PD-L1-functionalized nanofibers, CD86- functionalized nanofibers and PD-L1/CD86 dual-functionalized nanofibers. Azide-functionalized PD-L1-Ig and/or CD86-Ig were conjugated to the DBCO-functionalized nanofibers through strain-promoted azide-alkyne cycloaddition at a target loading of 10 µg of protein per 1 mg of DBCO-functionalized nanofibers, e.g., 5 µg of PD-L1 plus 5 µg of CD86 per 1 mg of nanofibers for the dual-functionalized nanofibers. **a**, Digital photographs of rhodamine B-labeled azide-functionalized PD-L1-Ig after conjugation to DBCO-functionalized nanofibers recorded before and after purification. **b**, Conjugation efficiencies for the preparation of rhodamine B-labeled PD-L1 and/or CD86 mono-/dual-functionalized nanofibers determined by fluorescence spectroscopy (to quantify the amount of conjugated rhodamine B-labeled PD-L1 and/or CD86). The conjugation efficiencies of mono-and/or dual-functionalized nanofibers were above 92%. **c**, UV-visible absorption spectra of 1 mg/mL non-fluorescently labeled PD-L1 and/or CD86 mono-/dual-functionalized nanofibers. The reduction in absorbance at 310 nm is a result of the change of cyclooctyne into triazole. **d**, Representative TEM images of DBCO-functionalized nanofibers, PD-L1-functionalized nanofibers, CD86-functionalized nanofibers, and PD-L1/CD86 dual-functionalized nanofibers.

**Supplementary Fig. 10.**
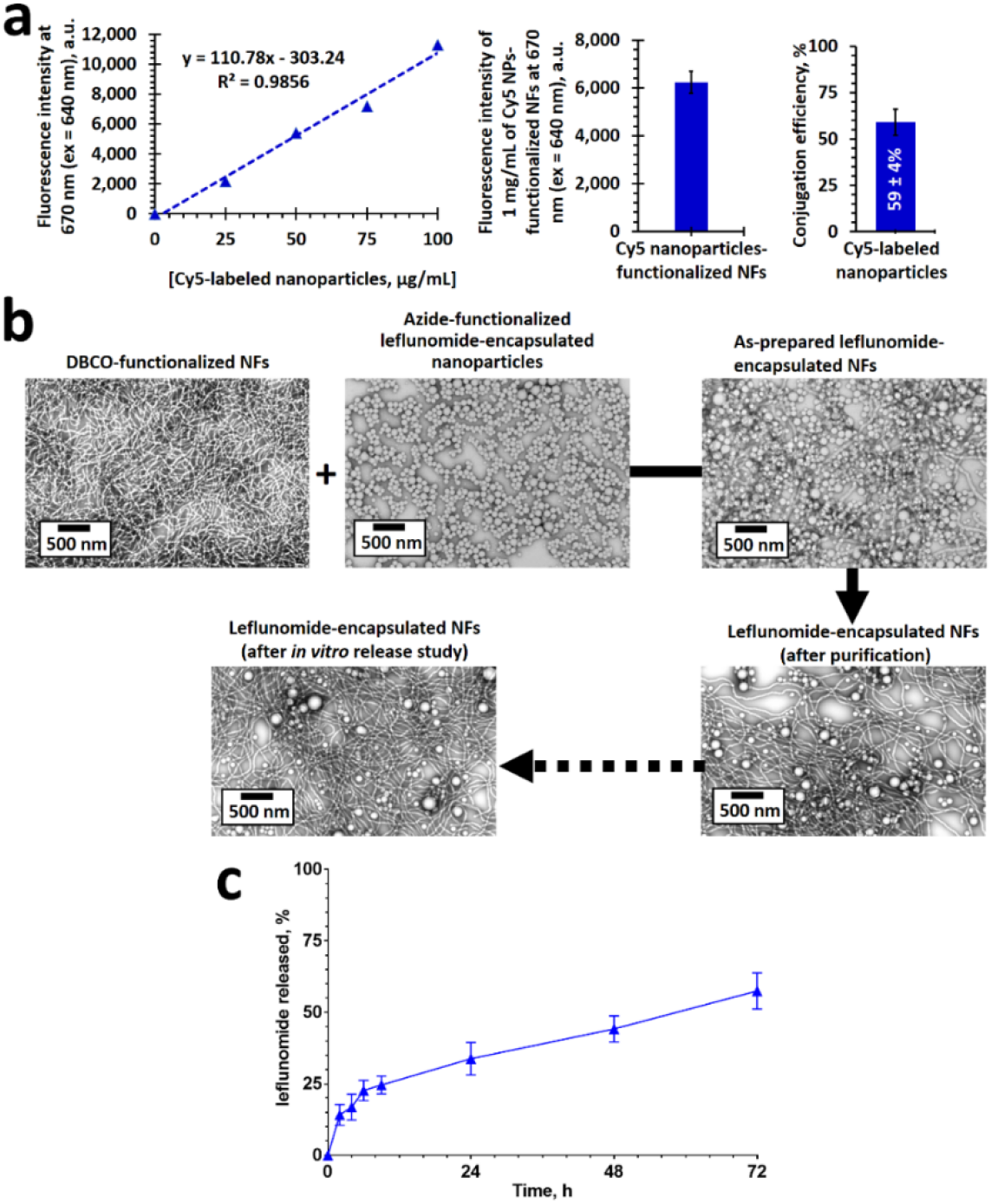
Characterization of leflunomide-encapsulated nanofibers. Azide-functionalized leflunomide-encapsulated nanoparticles were conjugated to the DBCO-functionalized nanofibers at a target loading of 0.1 mg of leflunomide-encapsulated nanoparticles or each 1 mg of DBCO-functionalized nanofibers. **a,** Quantification of Cy5-labeled leflunomide-encapsulated nanoparticles conjugated to DBCO-functionalized nanofibers *via* fluorescence spectroscopy. It was calculated that the conjugation efficiency was 59±4% (target loading: 200 µg of nanoparticles per 1 mg of nanofibers); i.e., approximately 118 µg of leflunomide-encapsulated nanoparticles were conjugated to 1 mg of DBCO-functionalized nanofibers. Since each 1 mg of the leflunomide-encapsulated nanoparticles contained 8.7±1.5 wt/wt% of leflunomide (target loading = 10 wt/wt%), each 1 mg of the leflunomide-encapsulated nanofibers encapsulated approximately 10 µg of the encapsulated leflunomide. **b**, Representative transmission electron microscopy images of leflunomide-encapsulated nanoparticles, DBCO-functionalized nanofibers, leflunomide-encapsulated nanofibers (before and after purification), and leflunomide-encapsulated nanofibers after the *in vitro* drug release study. **c**, *In vitro* leflunomide release kinetics under physiological conditions.

**Supplementary Fig. 11.**
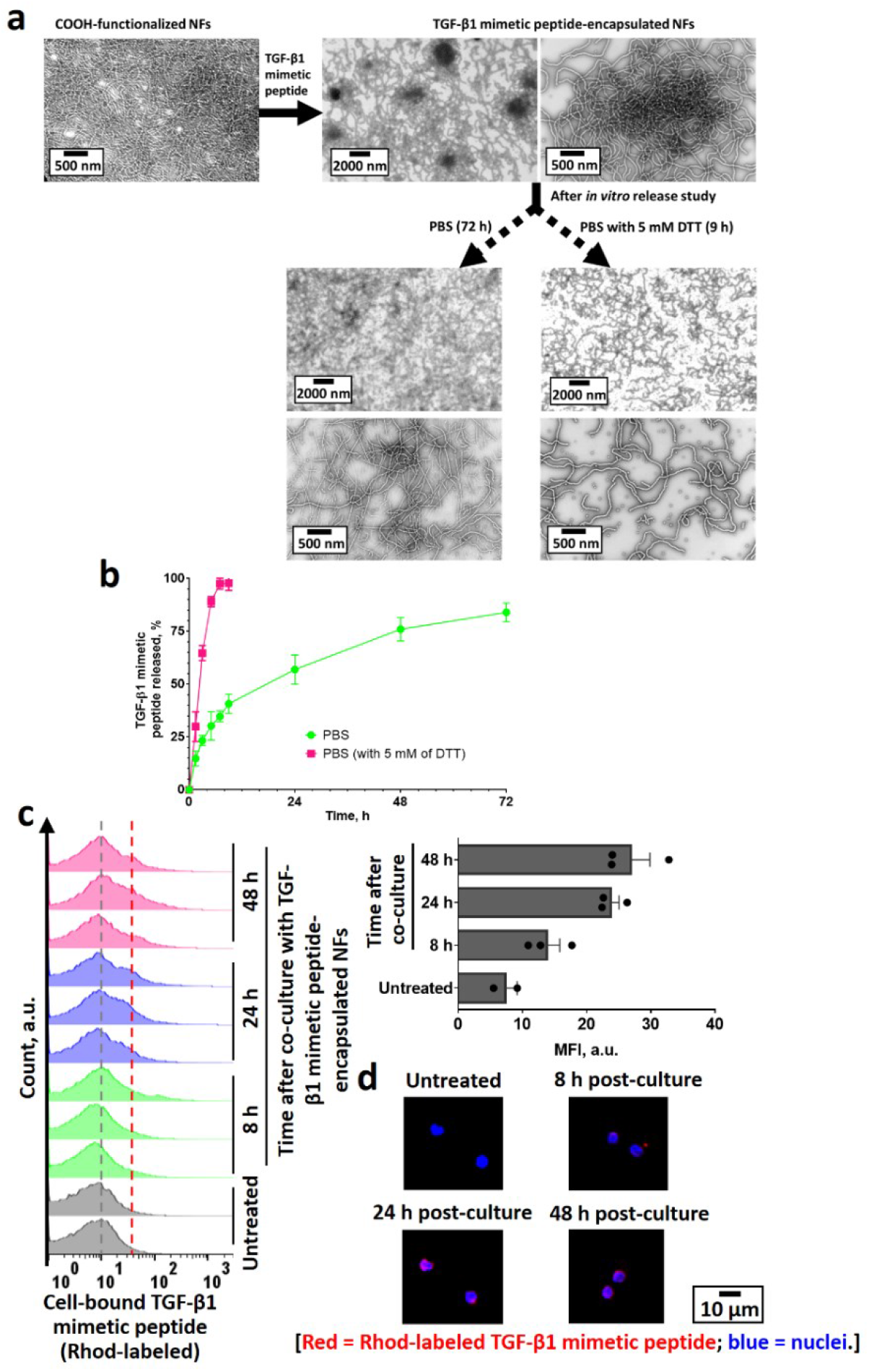
Characterization of TGF-β1 mimetic peptide-encapsulated. Cationic TGF-β1 mimetic peptide was adsorbed onto the anionic COOH-functionalized NFs through electrostatic interactions at a loading of 50 µg of peptide per 1 mg of nanofibers. Representative transmission electron microscopy images of unmodified COOH-functionalized nanofibers, TGF- β1 mimetic peptide-encapsulated nanofibers, and nanofibers after an *in vitro* release study under different physiological conditions. Physiosorbed rhodamine-labeled TGF-β1 mimetic peptide was gradually released from nanofibers and taken up by CD8^+^ T cells *in vitro*. **b**, *In vitro* TGF-β1 mimetic peptide release kinetics under reducing and non-reducing physiological conditions. **c**, Time-dependent fluorescence-activated cell sorting histograms (rhodamine channel) of CD8^+^ T cells after being cultured with rhodamine-labeled TGF-β1 mimetic peptide-encapsulated nanofibers for up to 48 h. **d**, Representative confocal laser scanning microscopy images of CD8^+^ T cells after being cultured with rhodamine-labeled TGF-β1 mimetic peptide-encapsulated nanofibers for up to 48 h.

**Supplementary Fig. 12.**
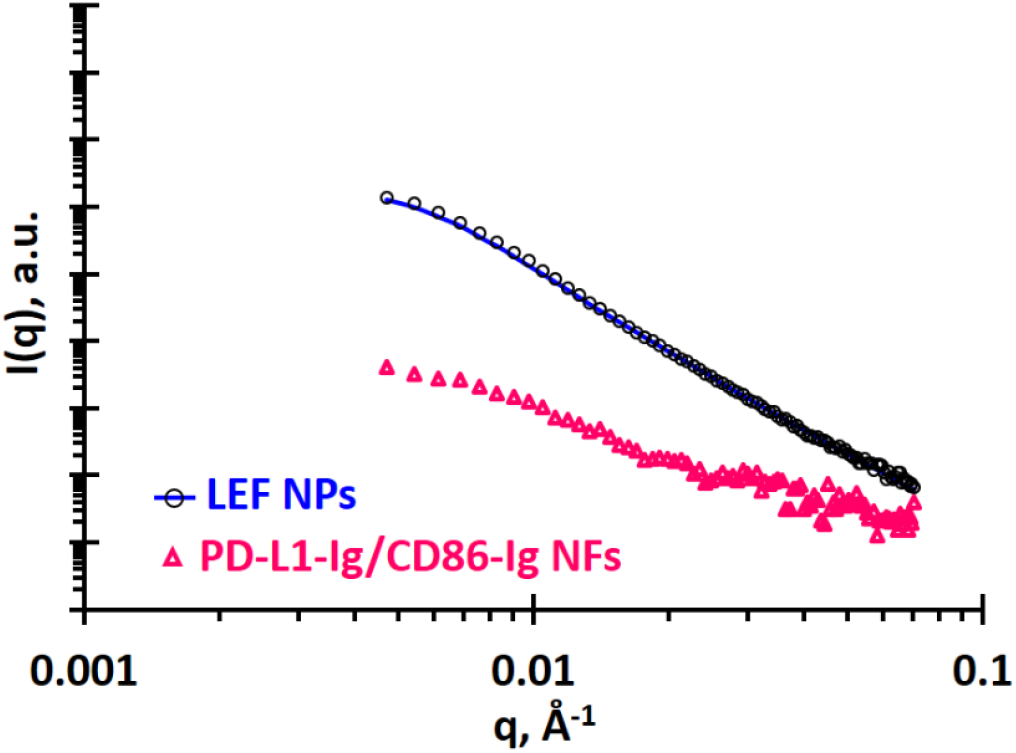
Solution small-angle X-ray scattering patterns recorded for leflunomide-encapsulated nanoparticles and a 1:1 mixture of PD-L1-Ig/CD86-Ig.

**Supplementary Fig. 13.**
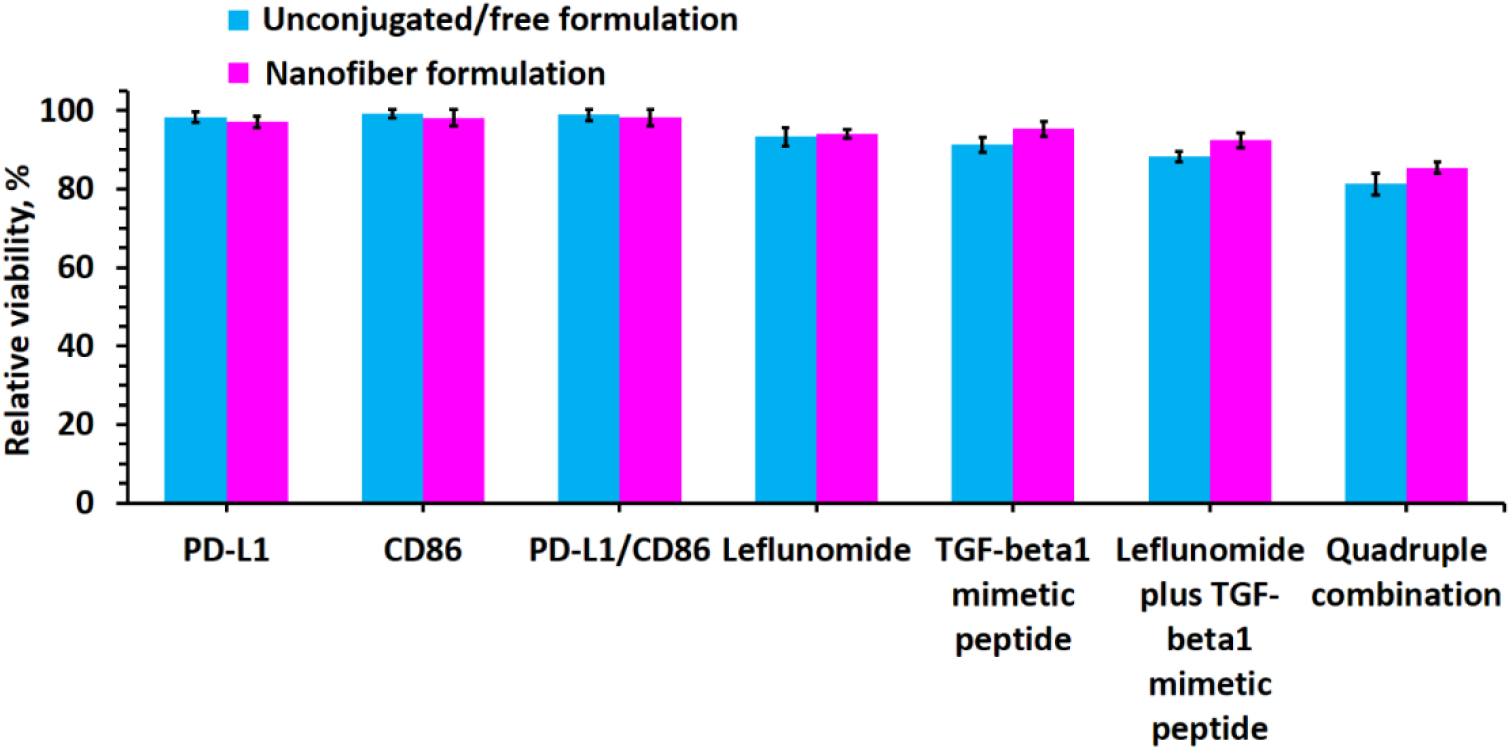
Biofunctionalized nanofibers show low *in vitro* toxicities when co-cultured with colon epithelial cells. *In vitro* toxicities of different functionalized nanofibers against colon epithelial cells. The viabilities of the colon epithelial cells (1×10^4^ cells per well, 1/100 times the dose used in the *in vivo* studies) were quantified by MTS assay after being cultured with 10 µg of nanofibers per well or 10 µg of PD-L1/CD86-encapsualated nanofibers plus 10 µg of leflunomide-encapsulated nanofibers and 10 µg of TGF-β1 mimetic peptide-encapsulated nanofibers per well for 4 days.

**Supplementary Fig. 14.**
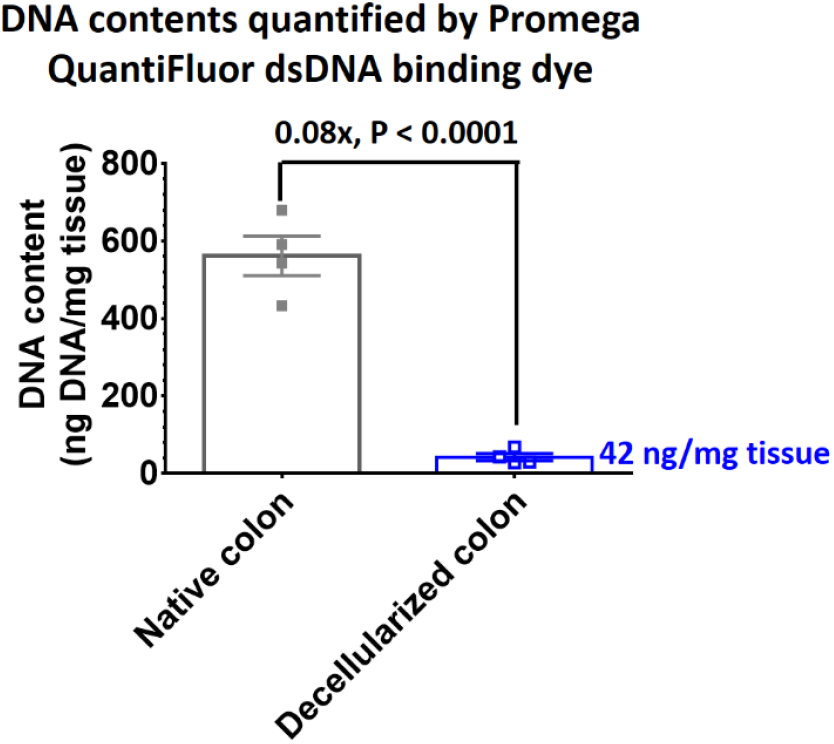
Spin-decellularization effectively decellularized the mouse colon. Quantification of DNA content in fresh and spin-decellularized mouse colon. Data are presented as the mean ± s.e.m. P value was analyzed using one-way ANOVA with Tukey’s HSD multiple comparisons post-hoc test.

**Supplementary Fig. 15.**
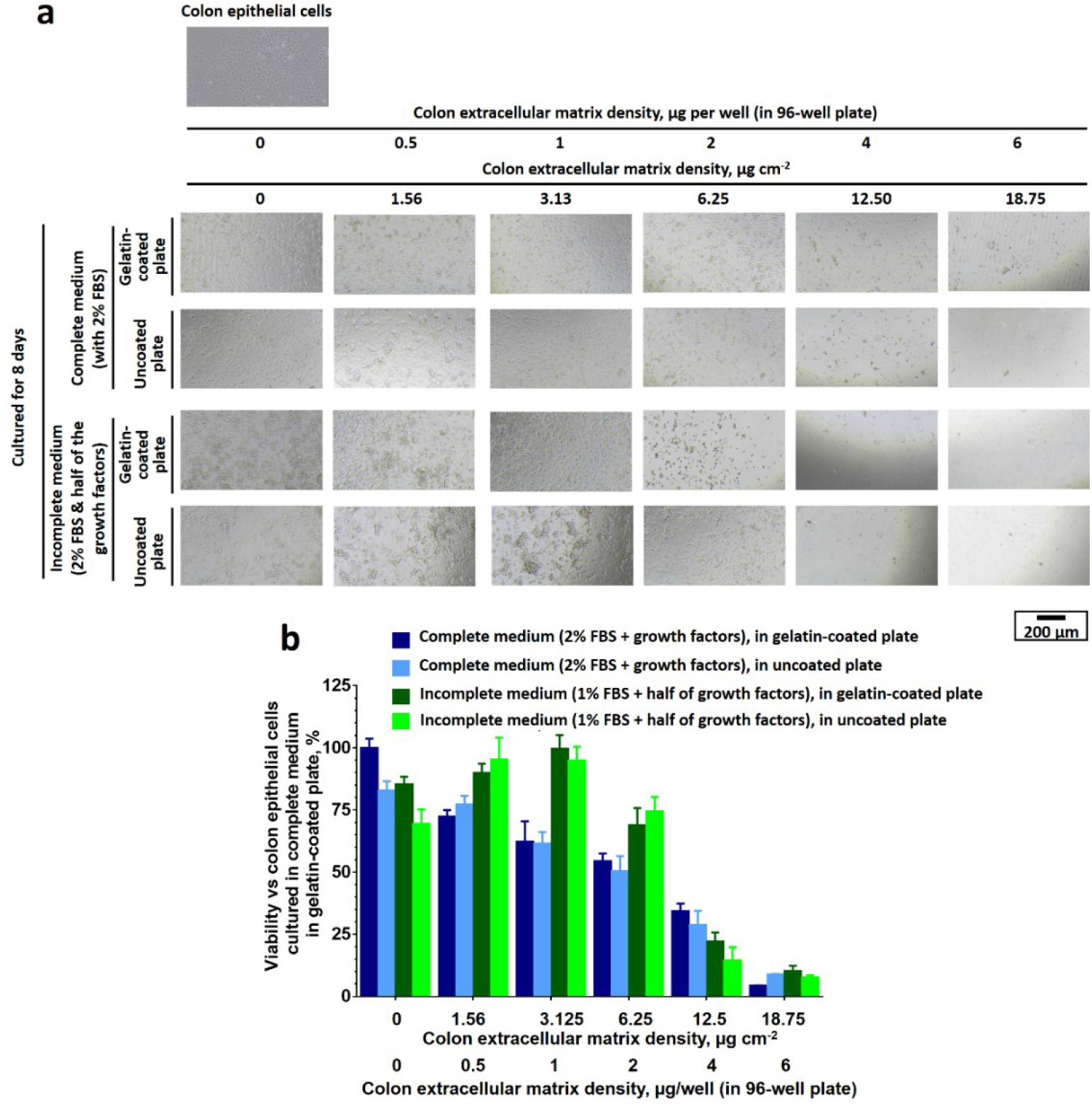
Decellularized colon extracellular matrix provides nutrients for the proliferation of colon epithelial cells in low growth factor growth media. The proliferation of colon epithelial cells after being co-cultured with different concentrations of decellularized colon extracellular matrix in complete tissue culture medium supplemented with 2% fetal bovine serum and 5 ng/mL recombinant epithelial growth factorand/or incomplete tissue culture medium supplemented with 1% fetal bovine serum and 2.5 ng/mL recombinant epithelial growth factor. **a,** Representative optical microscope images of colon epithelial cells after being cultured *in vitro* in the presence of different concentrations of decellularized colon extracellular matrix for 8 days. **b,** Relative viabilities of colon epithelial cells (*vs* colon epithelial cells cultured in extracellular matrix-free complete medium in gelatin-coated plates) after being cultured with different concentrations of decellularized colon extracellular matrix for 8 days. The *in vitro* study was performed in a non-adhesive 96-well plate. Unless specified, the wells were coated with gelatin before the *in vivo* study. The relative viabilities were quantified *via* MTS proliferation assay.

**Supplementary Fig. 16.**
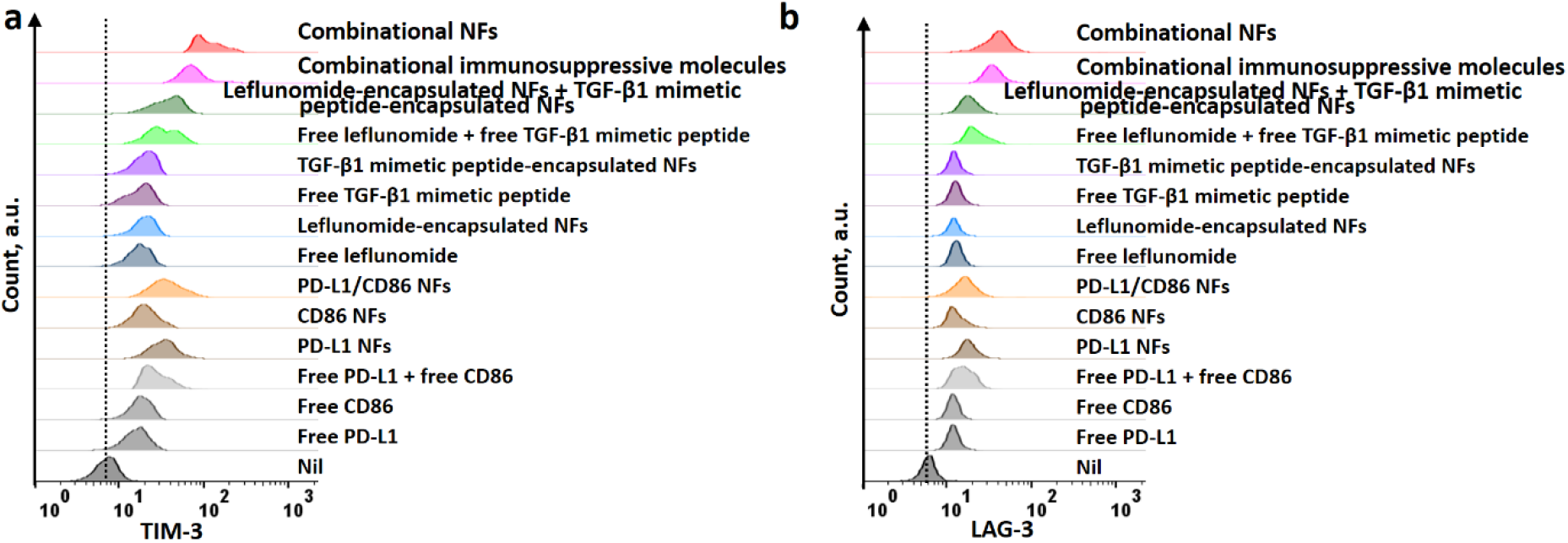
Immunosuppressive NFs induce CD8^+^ T cell exhaustion. **a-b**, Representative flow histograms show TIM-3 (**a**) and LAG-3 (**b**) expressions on stimulated CD8^+^ T cells after incubation with different immunosuppressive nanofibers.

**Supplementary Fig. 17.**
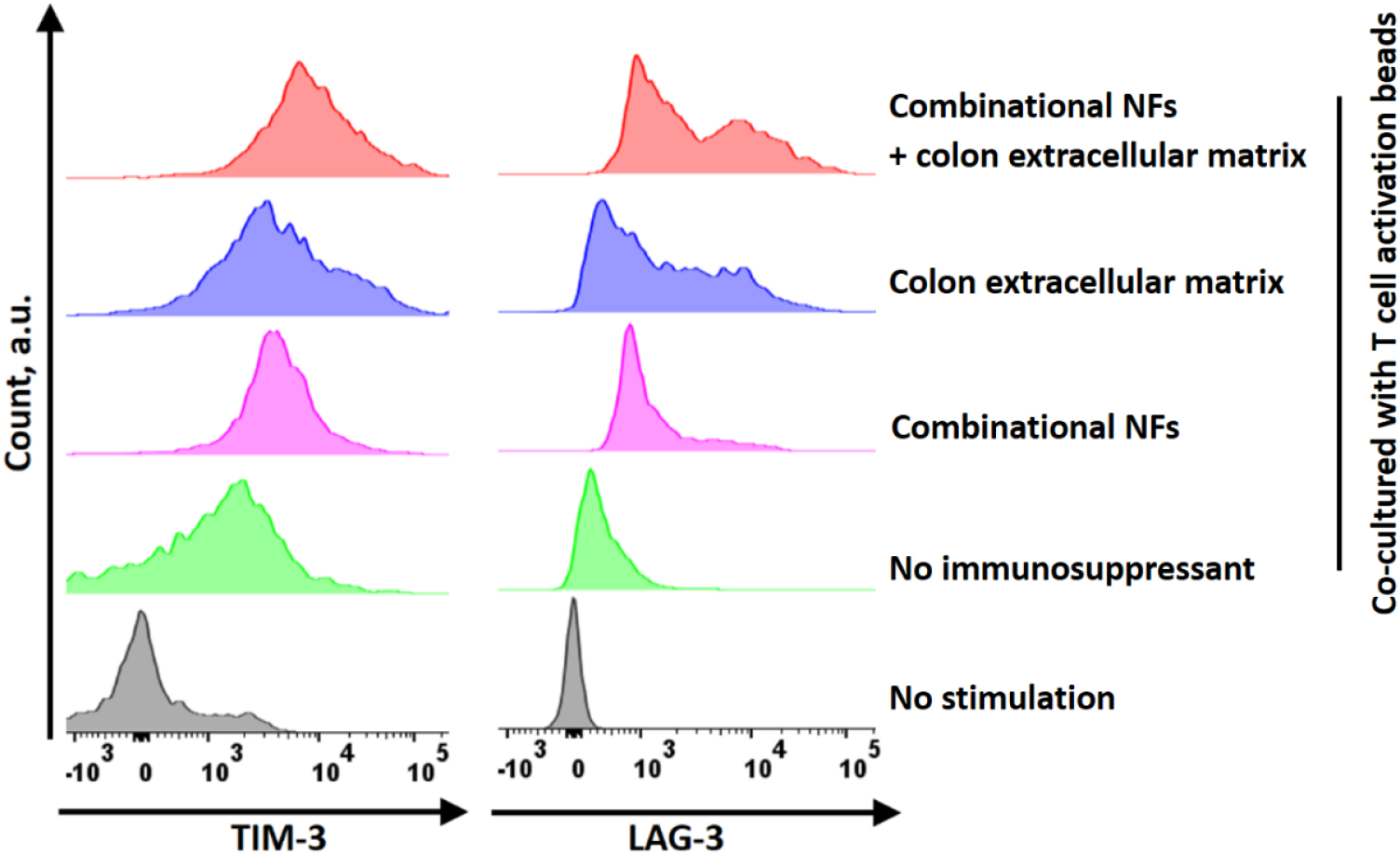
The combination of combinational immunosuppressive nanofibers and colon extracellular matrix is more effective than combinational immunosuppressive nanofibers alone in exhausting stimulated CD8^+^ T cells. **a-b**, Representative flow histograms show the TIM-3 (**a**) and LAG-3 (**b**) expressions on stimulated CD8^+^ T cells after incubation with different immunosuppressive nanofibers.

**Supplementary Fig. 18.**
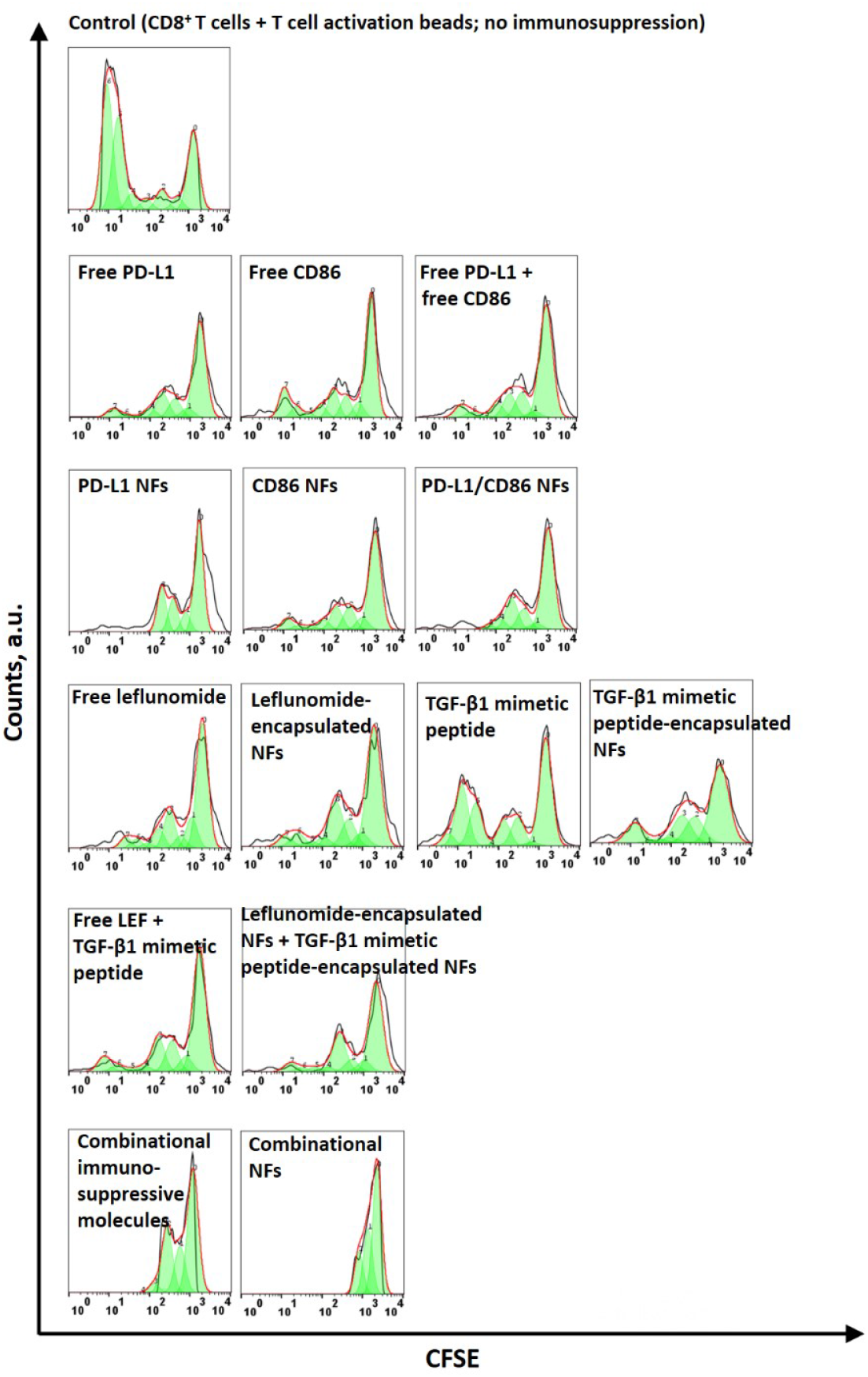
Immunosuppressive nanofibers inhibit the proliferation of stimulated CD8^+^ T cells. Representative CFSE profiles (and their best fit to the proliferation model) recorded for the CFSE-labeled CD8^+^ T cells cultured in the presence of different immunosuppressive NFs under stimulated conditions for 72 h. The CFSE-labeled CD8^+^ T cells were stimulated using anti-CD3/anti-CD28 T cell activation beads (Dynbeads) at a 1:1 ratio.

**Supplementary Fig. 19.**
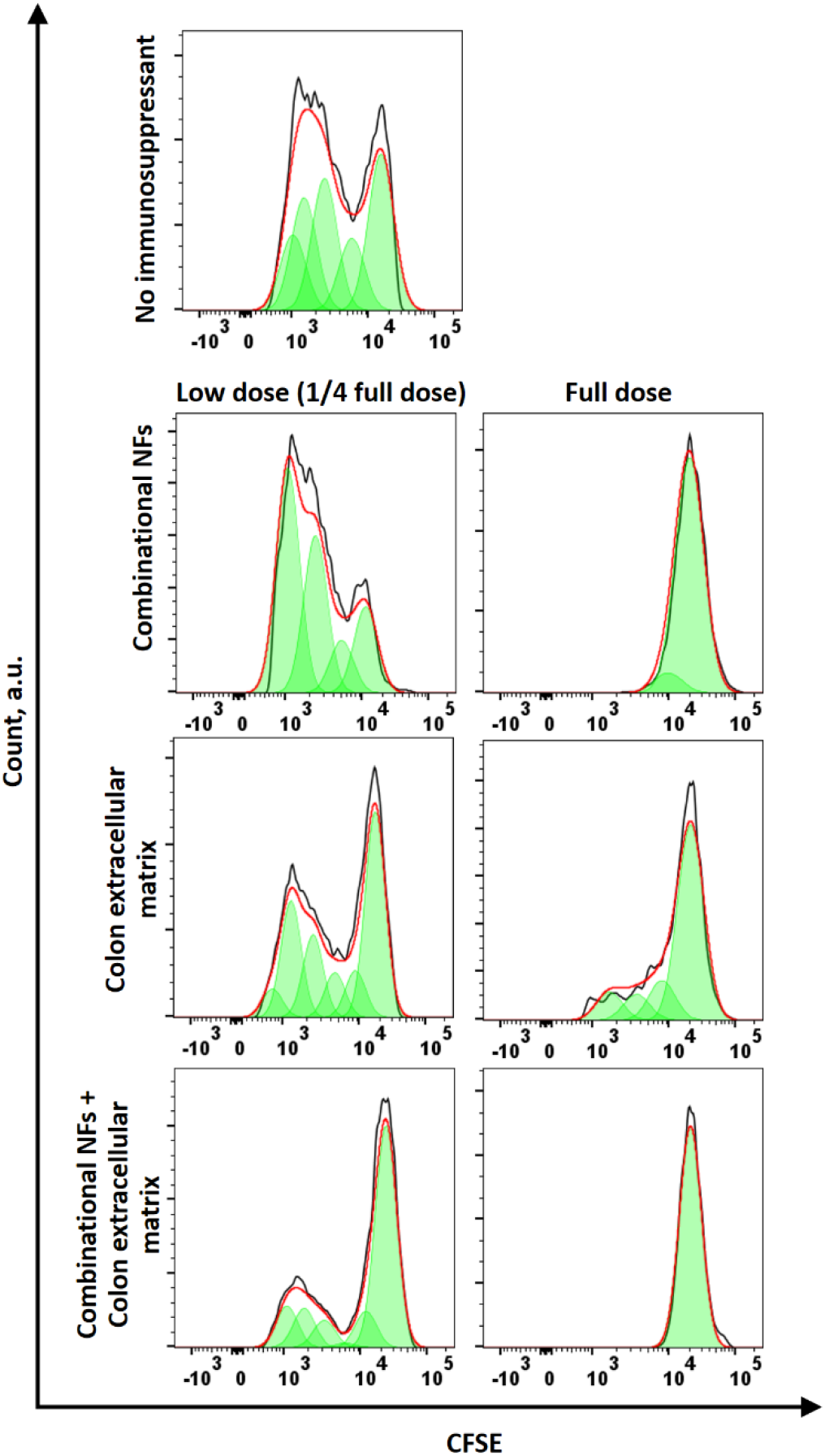
The combination of combinational immunosuppressive nanofibers and colon extracellular matrix is more effective than combinational immunosuppressive nanofibers in inhibiting the proliferation of stimulated CD8^+^ T cells. Representative CFSE profiles (and their best fit to the proliferation model) recorded for the CFSE-labeled CD8^+^ T cells cultured in the presence of different immunosuppressive NFs under stimulated conditions for 72h. The CFSE-labeled CD8^+^ T cells were stimulated using anti-CD3/anti-CD28 T cell activation beads (Dynbeads) at a 1:1 ratio.

**Supplementary Fig. 20.**
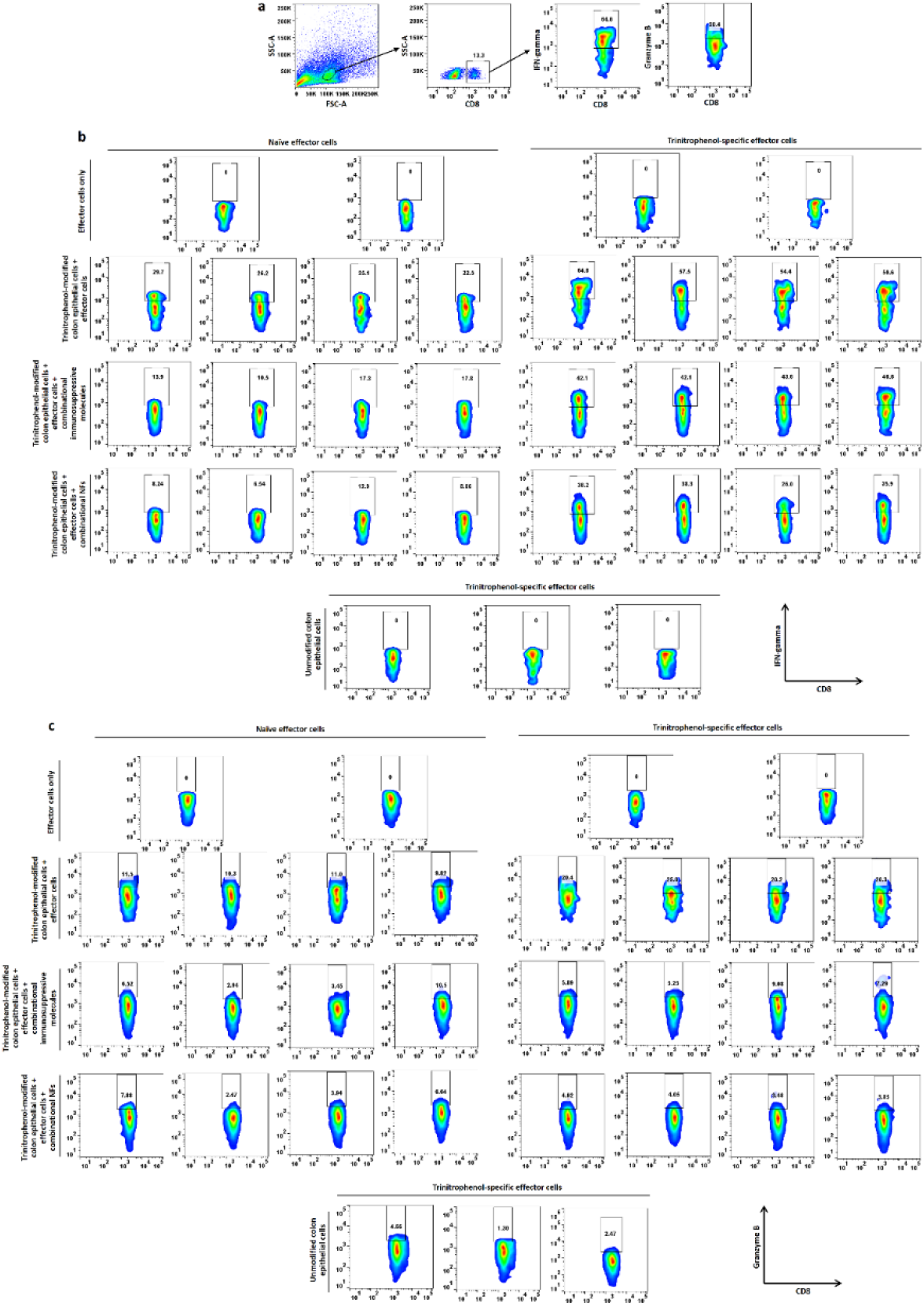
Combinational immunosuppressive nanofibers inhibit antigen-specific CD8^+^ T cell activation against trinitrophenol-modified colon epithelial cells *in vitro*. **a**, Gating strategy for the quantification of intracellular IFN-gamma and granzyme B expression in CD8^+^ T cells (among cocultured splenocytes) *via* fluorescence-activated cell sorting. **b-c**, Fluorescence-activated cell sorting dot plots showing the IFN-gamma **(b)** and granzyme B (**c**) expressions of trinitrophenol-sensitized and non-sensitized (naïve) CD8^+^ T cells after incubation with trinitrophenol-modified colon epithelial cells (or unmodified colon epithelial cells, as a control) in the presence or absence of combinational immunosuppressive nanofibers (or combinational immunosuppressive molecules) for 48 h. The effector cells are trinitrophenol-sensitized or non-sensitized splenocytes. The effector-to-target ratio was 20:1 (2×10^5^ splenocytes:1×10^4^ colon epithelial cells).

**Supplementary Fig. 21.**
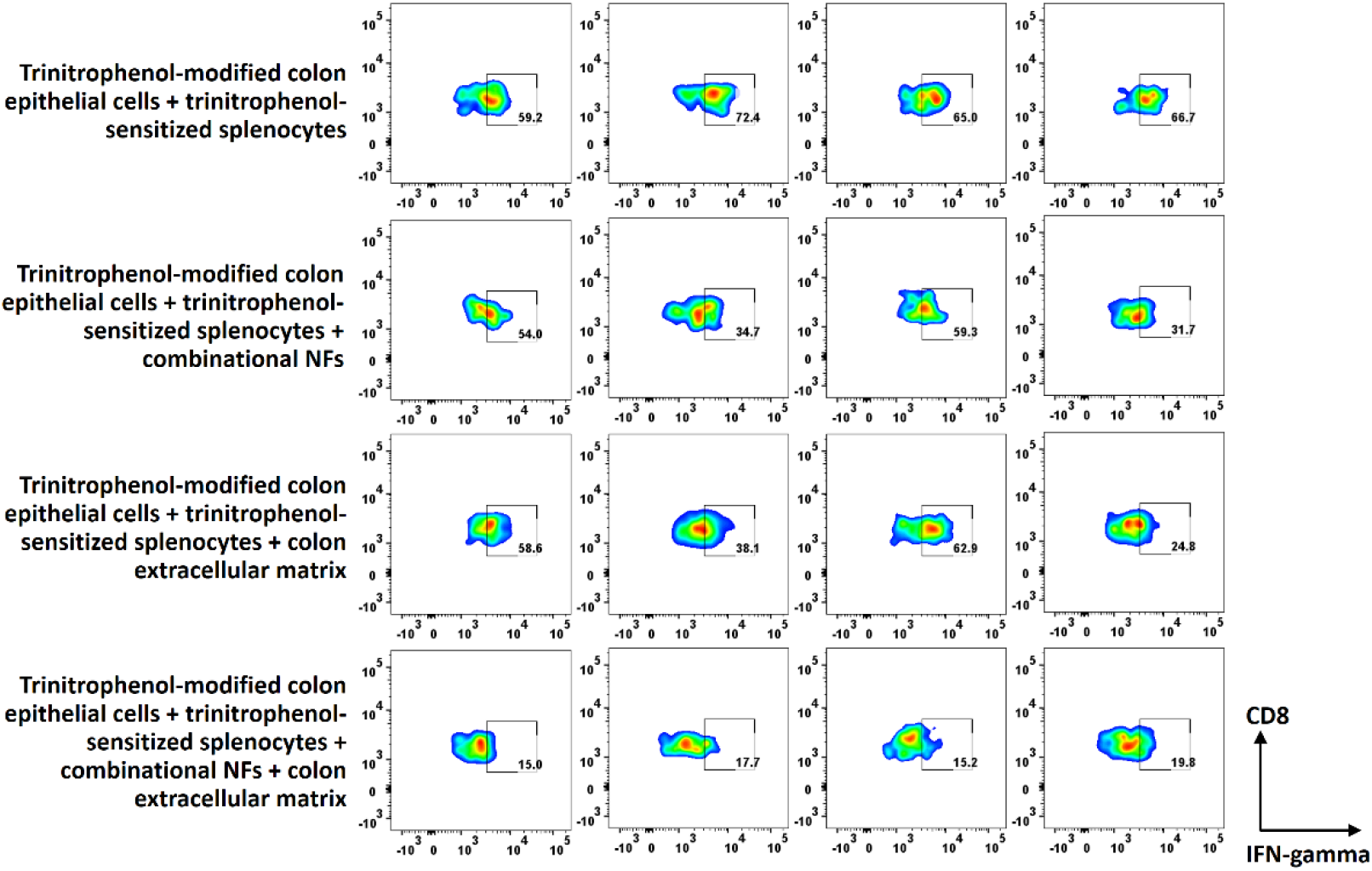
The combination of combinational immunosuppressive nanofibers and colon extracellular matrix is more effective than combinational immunosuppressive nanofibers in inhibiting antigen-specific CD8^+^ T cell activation against trinitrophenol-modified colon epithelial cells *in vitro*. fluorescence-activated cell sorting dot plots showing the IFN- expression of trinitrophenol-sensitized and non-sensitized (naïve) CD8^+^ T cells after incubation with trinitrophenol-modified colon epithelial cells (or unmodified COL EPI cells, as a control) in the presence or absence of combinational immunosuppressive nanofibers and colon extracellular matrix for 48 h. The effector cells are trinitrophenol-sensitized or non-sensitized splenocytes. The effector-to-target ratio was 20:1 (2×10^5^ splenocytes:1×10^4^ colon epithelial cells).

**Supplementary Fig. 22.**
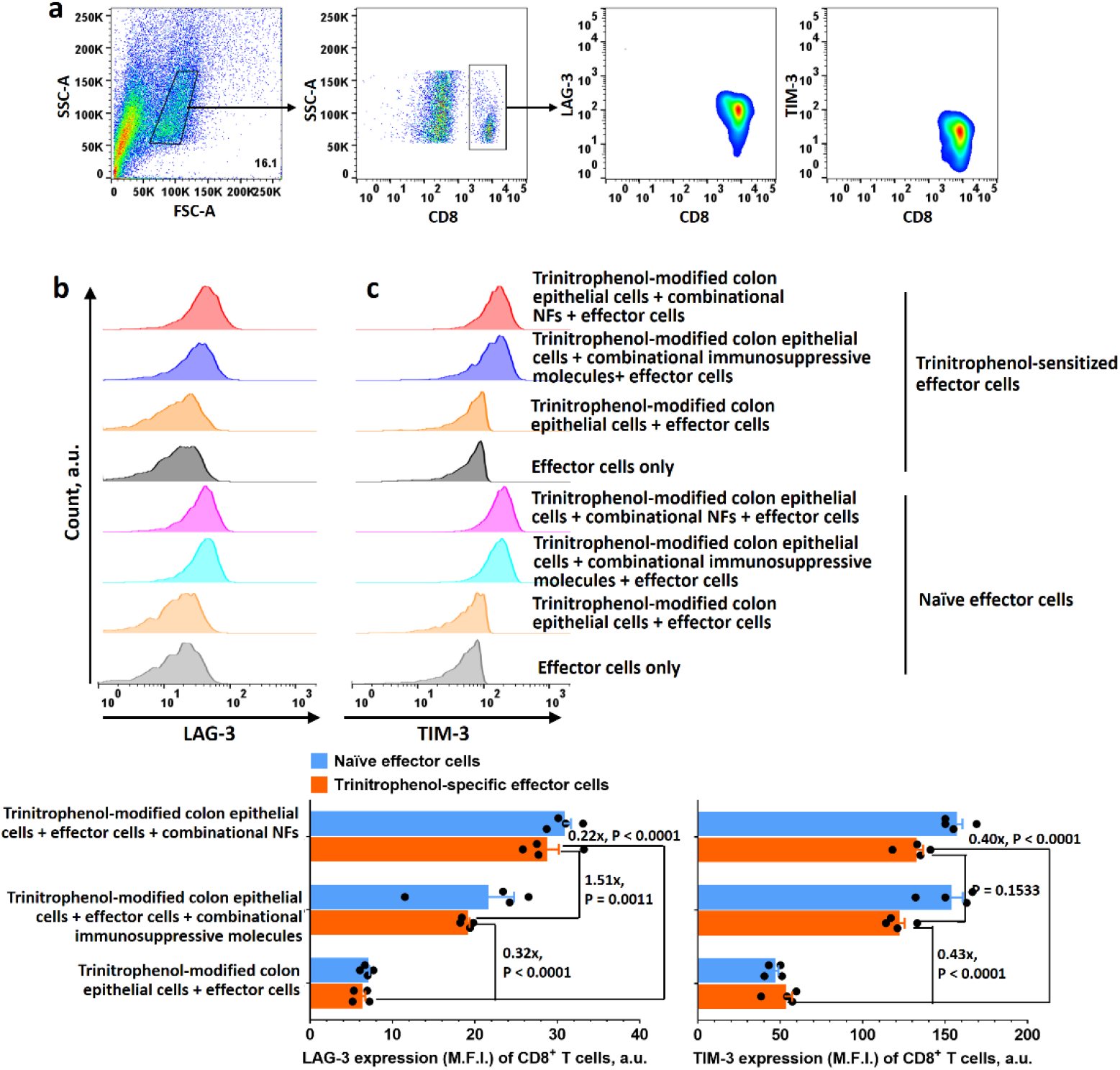
Combinational immunosuppressive nanofibers inhibit antigen-specific CD8^+^ T cell activation triggered by trinitrophenol-modified colon epithelial cells *in vitro* by inducing T cell exhaustion. **a**, Gating strategy for the quantification of T cell inhibition markers on CD8^+^ T cells (among the co-cultured splenocytes) *via* fluorescence-activated cell sorting. **b-c**, Fluorescence-activated cell sorting dot plots showing the LAG-3 (**b**) and TIM-3 (**c**) expressions of trinitrophenol-sensitized and non-sensitized (naïve) CD8^+^ T cells after incubation with trinitrophenol-modified colon epithelial cells in the presence or absence of combinational immunosuppressive nanofibers (or combinational immunosuppressive molecules) for 48 h. The effector cells are trinitrophenol-sensitized or non-sensitized splenocytes. The effector-to-target ratio was 20:1 (2×10^5^ splenocytes:1×10^4^ colon epithelial cells). Data are presented as the mean ± standard error of the mean (s.e.m.). All P values were analyzed using two-way ANOVA with Tukey’s HSD multiple comparisons post-hoc test.

**Supplementary Fig. 23.**
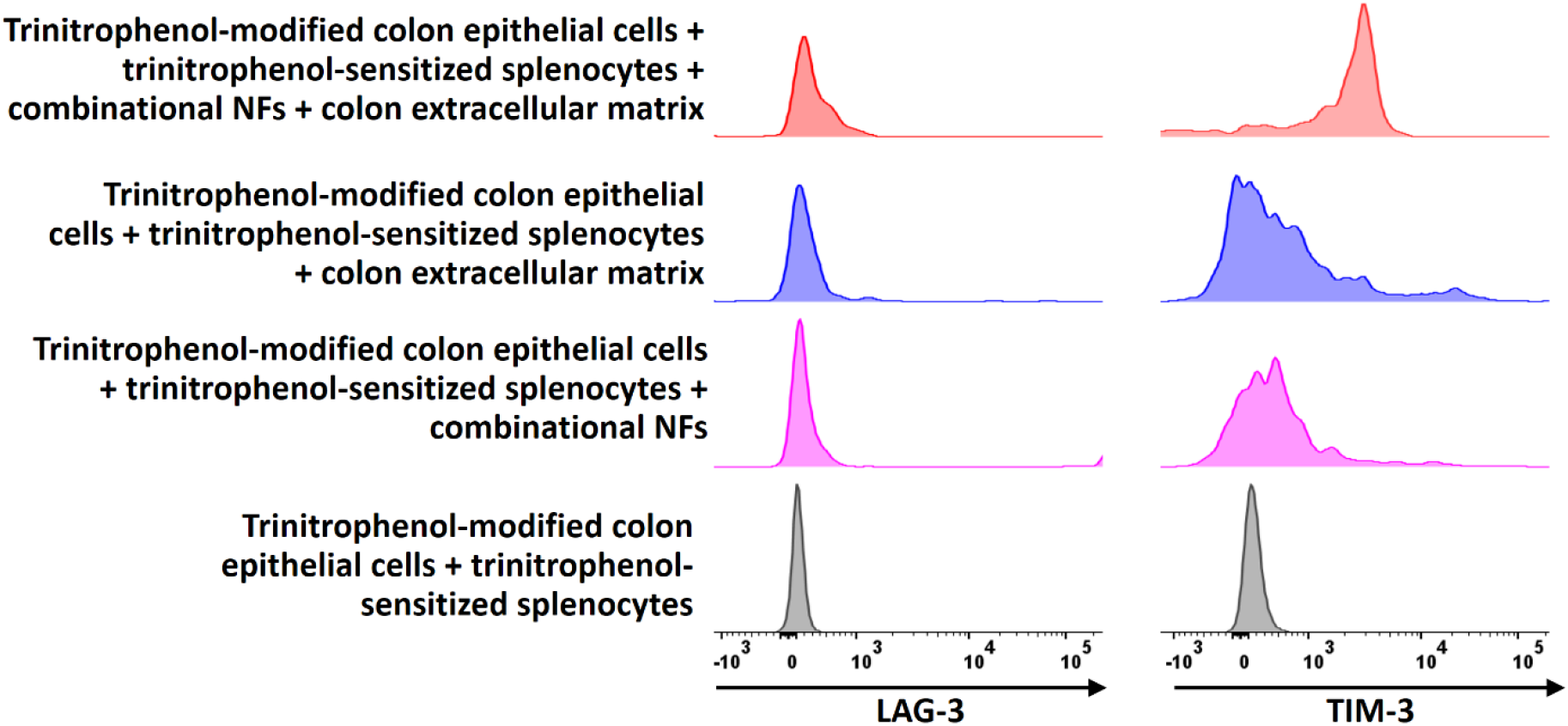
The combination of combinational immunosuppressive nanofibers and colon extracellular matrix is more effective in inducing antigen-specific T cell exhaustion *in vitro*. Fluorescence-activated cell sorting dot plots showing the LAG-3 and TIM-3 expression in trinitrophenol-sensitized and non-sensitized (naïve) CD8^+^ T cells after incubation with trinitrophenol-modified colon epithelial cells in the presence or absence of combinational immunosuppressive nanofibers and colon extracellular matrix for 48 h. The effector cells are trinitrophenol-sensitized or non-sensitized splenocytes. The effector-to-target ratio was 20:1 (2×10^5^ splenocytes:1×10^4^ colon epithelial cells). Data are presented as the mean ± standard error of the mean (s.e.m.). All P values were analyzed using two-way ANOVA with Tukey’s HSD multiple comparisons post-hoc test.

**Supplementary Fig. 24.**
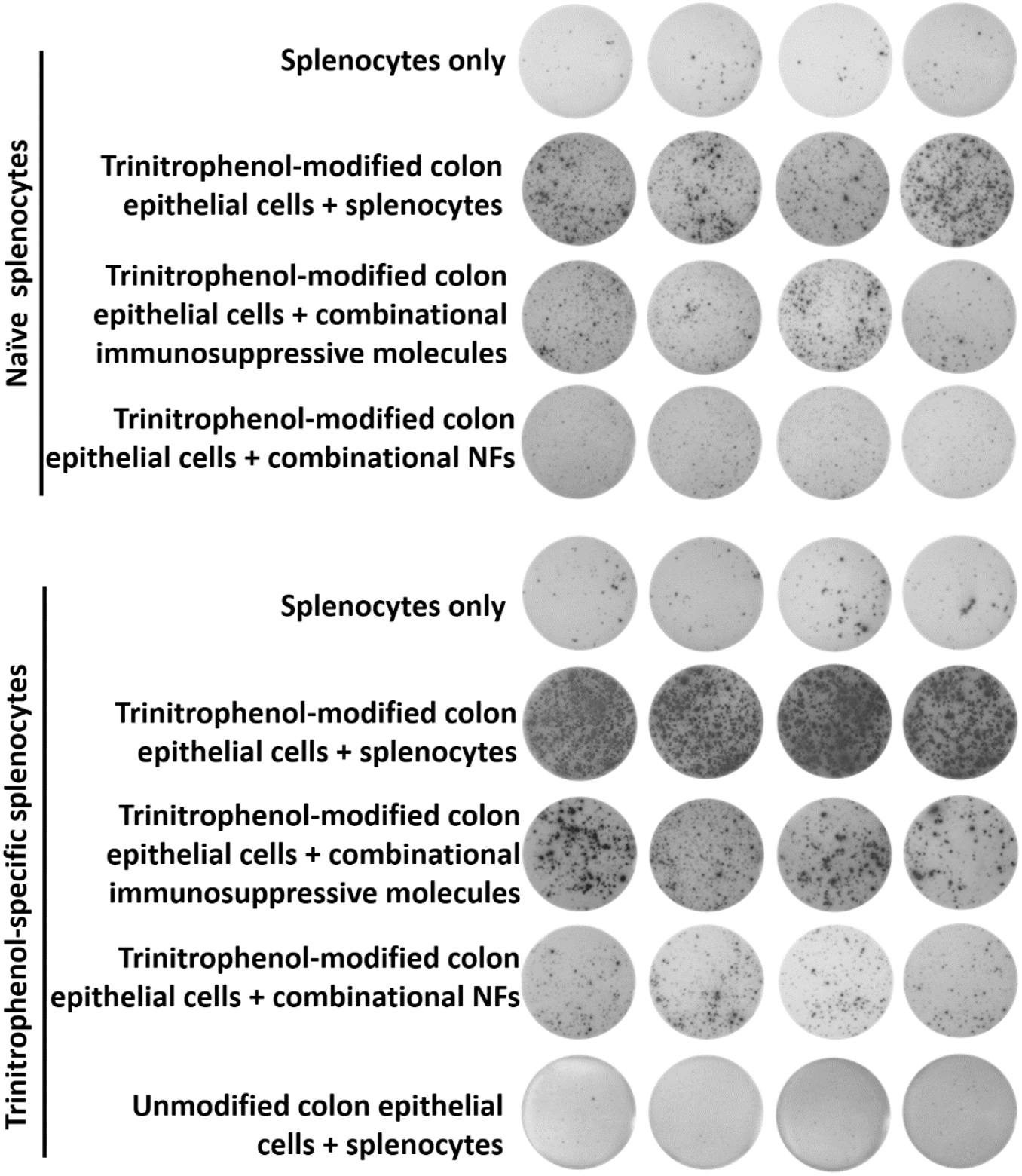
Combinational immunosuppressive nanofibers inhibit the secretion of proinflammatory IFN-gamma from antigen-specific splenocytes when co-cultured with trinitrophenol-modified colon epithelial cells *in vitro*. Digital photograph of IFN-gamma ELISpot wells after TNP-modified colon epithelial cells were co-cultured with trinitrophenol-sensitized or naïve splenocytes in the presence or absence of combinational immunosuppressive nanofibers or combinational immunosuppressive molecules for 48 h.

**Supplementary Fig. 25.**
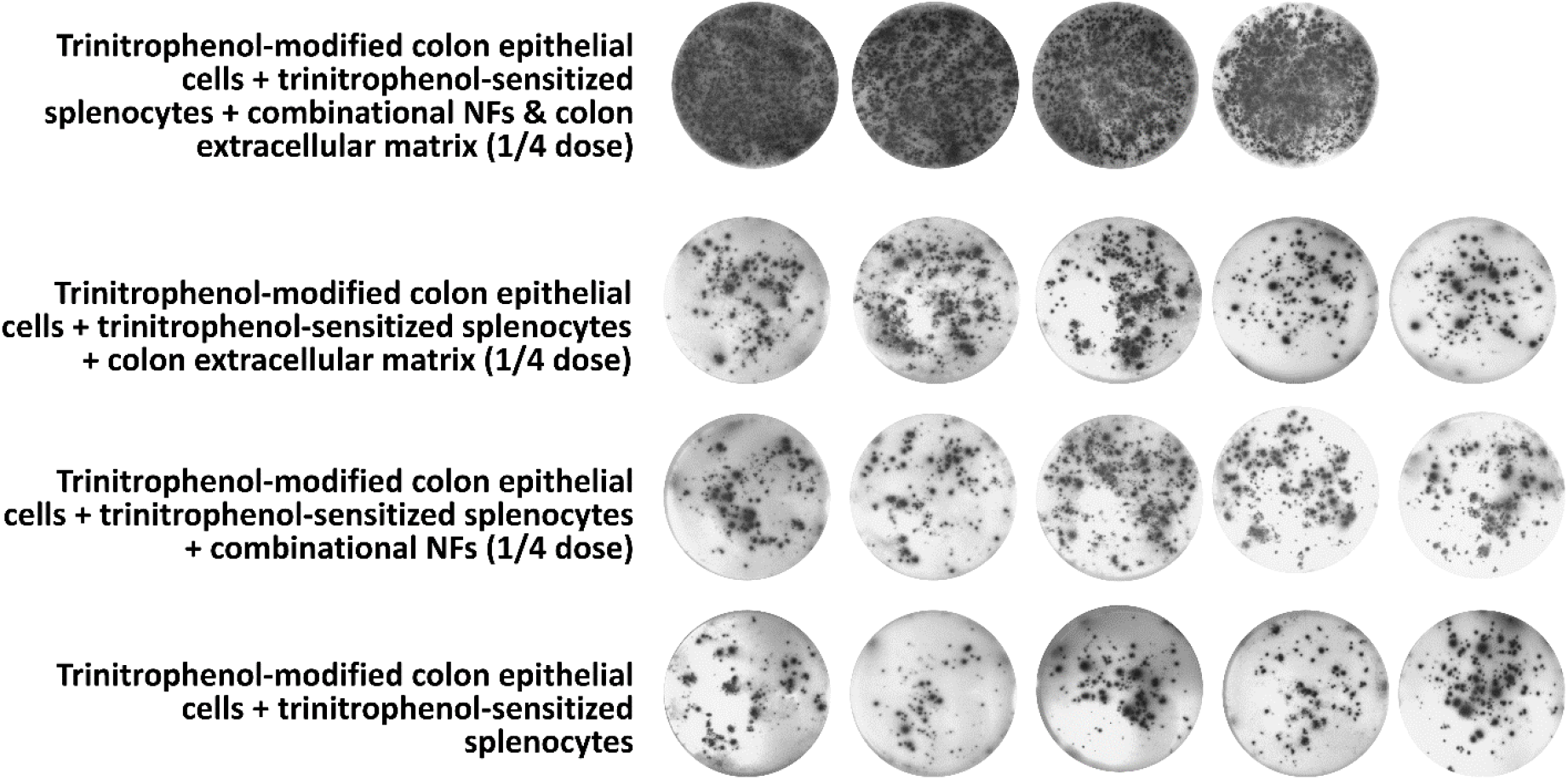
The combination of combinational immunosuppressive nanofibers and colon extracellular matrix effectively inhibits the secretion of proinflammatory IFN-gamma from antigen-specific splenocytes when co-cultured with trinitrophenol-modified colon epithelial cells *in vitro*. Digital photograph of IFN-gamma ELISpot wells after TNP- modified colon epithelial cells were co-cultured with TNP-sensitized or naïve splenocytes in the presence combinational immunosuppressive nanofibers and/or colon extracellular matrix for 48 h.

**Supplementary Fig. 26.**
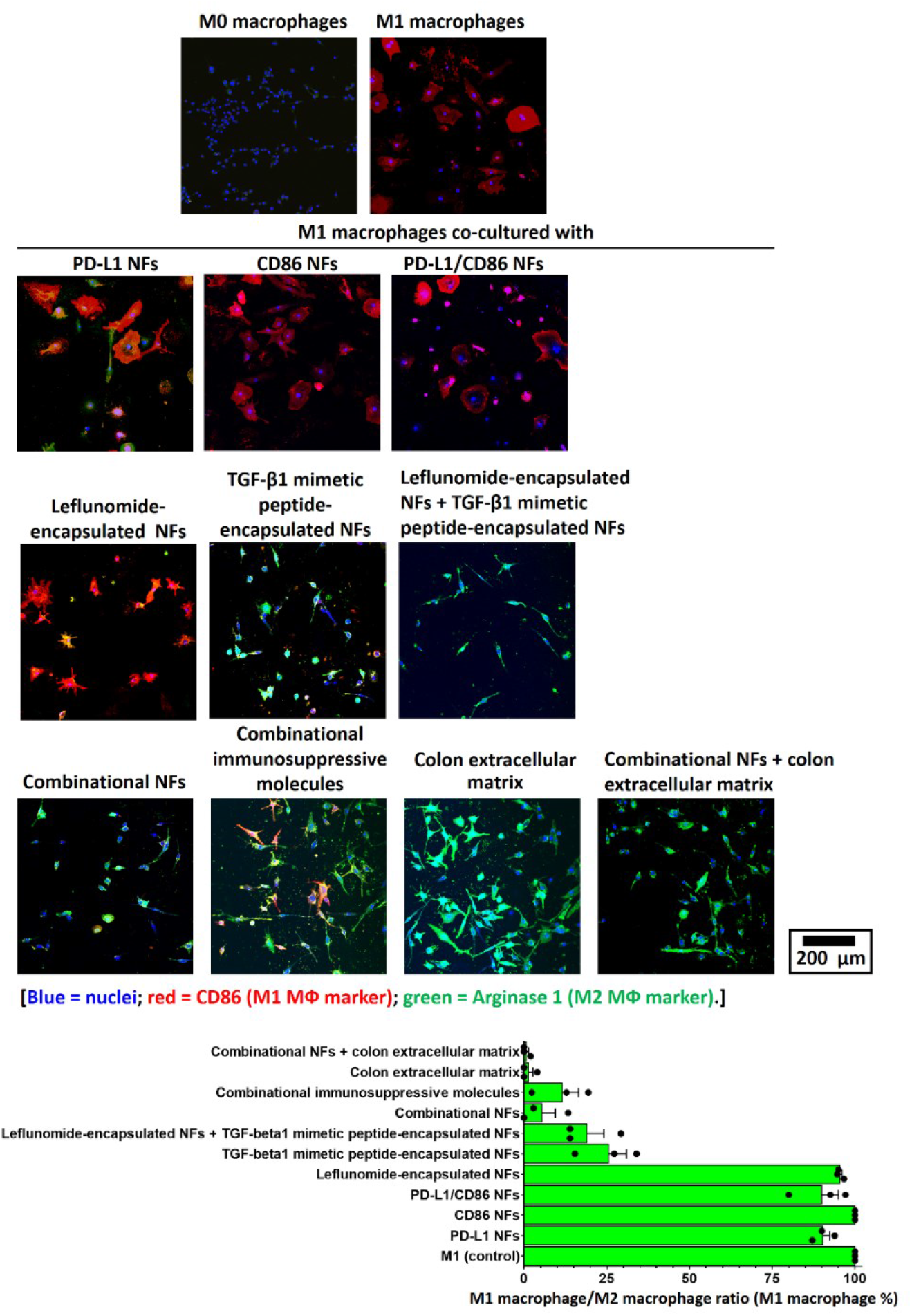
The combination of combinational immunosuppressive nanofibers and colon extracellular matrix induces classically-activated M1 macrophages differentiation into alternatively-activated M2 macrophages *in vitro*. Representative CLSM images of IFN-γ- and LPS-induced classically-activated M1 macrophages after cultured with different immunosuppressive nanofibers, colon extracellular matrix and their combination for 48 h. The round-shape M1 MФs (with cytoplasmic extension on the cellular surface) differentiated into elongated alternatively-activated M2 macrophages with cytoplasmic extension on the apical ends after co-cultured with combinational immunosuppressive nanofibers and colon extracellular matrix.

**Supplementary Fig. 27.**
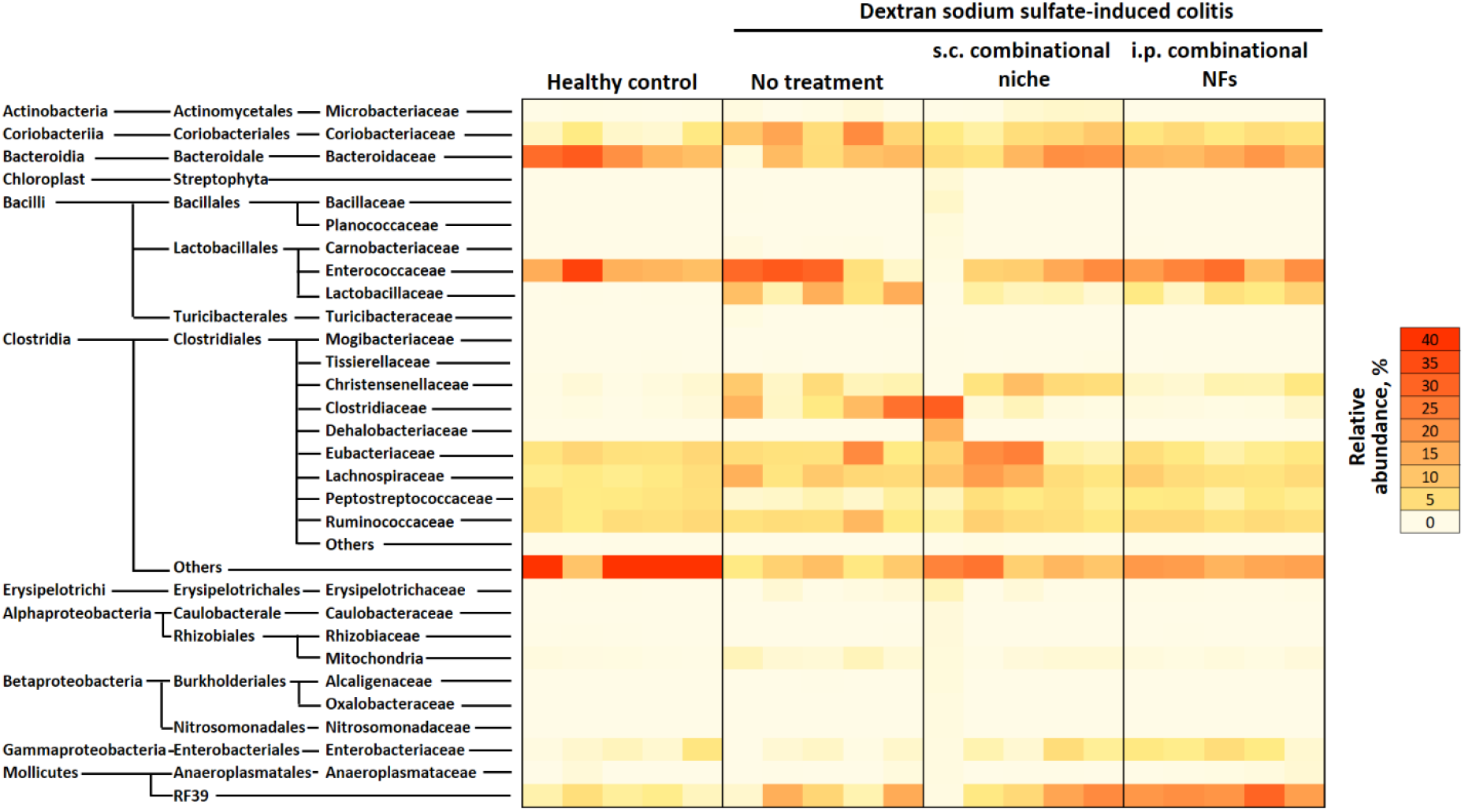
Colitis treatment with subcutaneously-administered colon-specific immune niches or intraperitoneally-administered combinational immunosuppressive nanofibers resolved imbalance of the pathogenic and commensal fecal microbiome in dextran sodium sulfate-induced colitis mice. Heatmap of the relative abundance of the microbiome (rows) for each dextran sodium sulfate-induced colitis mouse (column) after different treatments. The abundance is shown as a relative percentage. Gut microbiome analysis (by 16S rRNA sequencing) was performed on feces collected 5 days after colitis treatment.

**Supplementary Fig. 28.**
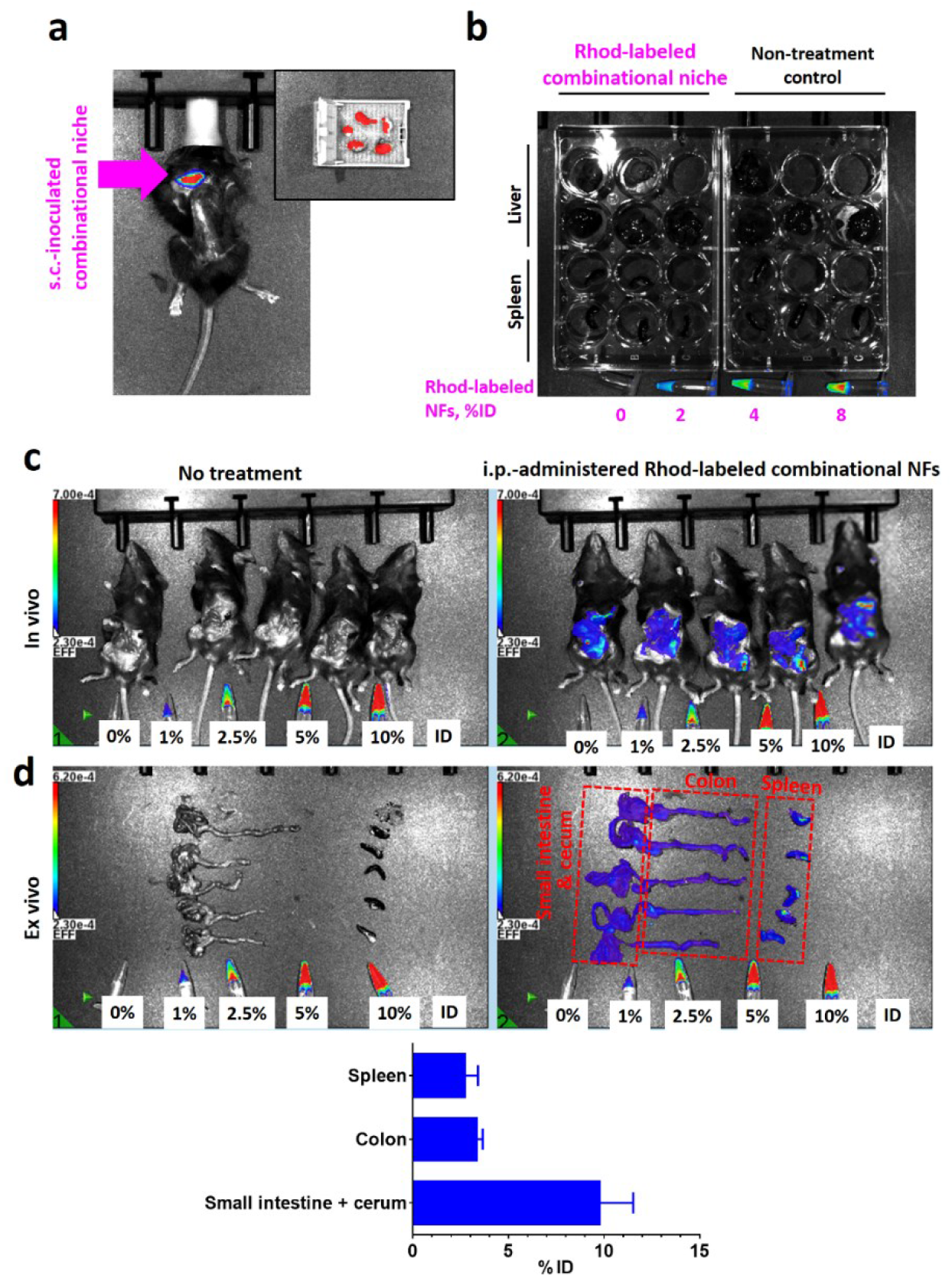
Subcutaneously-inoculated combinational colon-specific immune niche is retained at the inoculation site, but intraperitoneal-administered combinational immunosuppressive nanofibers diffuse throughout the abnormal cavity after administration. **a,** Ex vivo fluorescent images of inoculation site and grafts preserved 3 days after subcutaneous inoculation of combinational colon-specific immune niche constructed from rhodamine-labeled nanofibers in dextran sodium sulfate-induced colitis mice. **b,** *Ex vivo* fluorescent images of spleen and liver preserved 3 days after subcutaneous inoculation of combinational colon-specific immune niche. There is no indication for the clearance of nanofibers by the immune cells that would accumulate in the liver and spleen. **c-d,** *In vivo* and *ex vivo* fluorescence images recorded for dextran sodium sulfate-induced colitis mice 3 days after therapeutic treatment with i.p.- administered rhodamine-labeled combinational immunosuppressive nanofibers.

**Supplementary Fig. 29.**
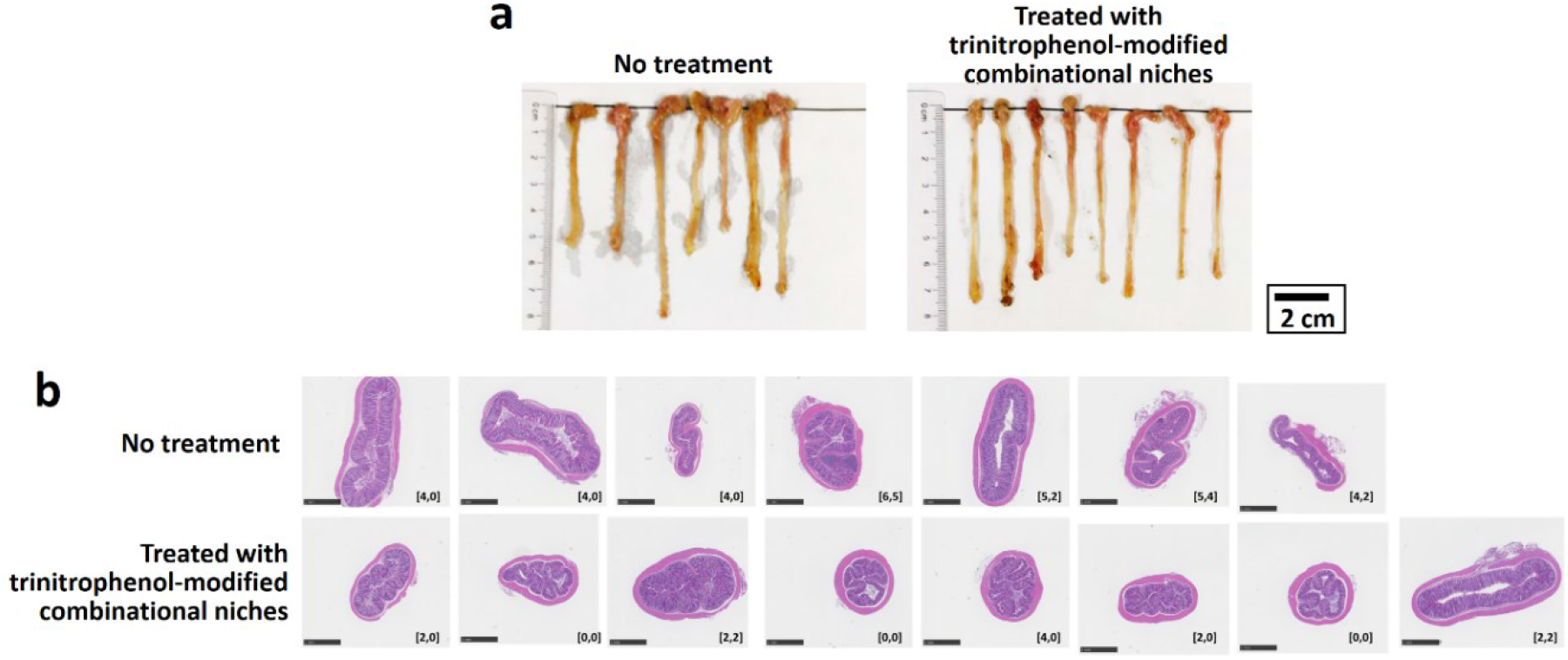
Trinitrophenol-modified combinational colon-specific immune niche effectively ameliorates trinitrophenol-induced colitis. **a,** Digital photographs of colons preserved at the study endpoint. **b,** Representative H&E-stained sections of each colon and colon damage scores after different colitis treatments. The inserted label in each image represents the corresponding colon epithelial damage score and inflammatory cell infiltration score.

**Supplementary Fig. 30.**
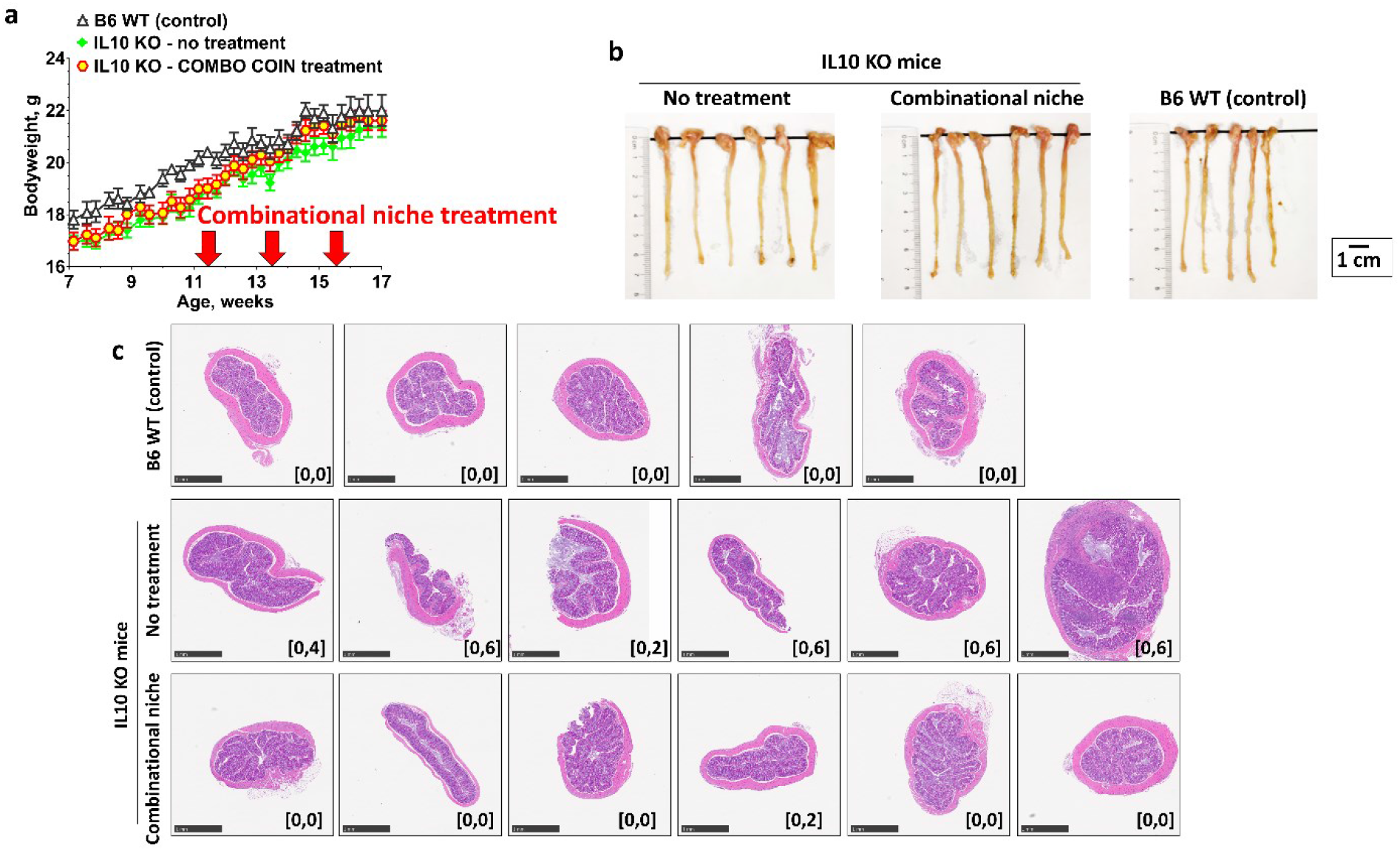
Combinational colon-specific immune niche effectively ameliorates colitis in IL10 KO mice that spontaneously develop colitis. **a,** Bodyweight change. **b,** Digital photographs of colons preserved at the study endpoint. **c,** Representative H&E-stained sections of each colon and colon damage scores after different colitis treatments. The inserted label in each image represents the corresponding colon epithelial damage score and inflammatory cell infiltration score. (n = 6)

**Supplementary Fig. 31.**
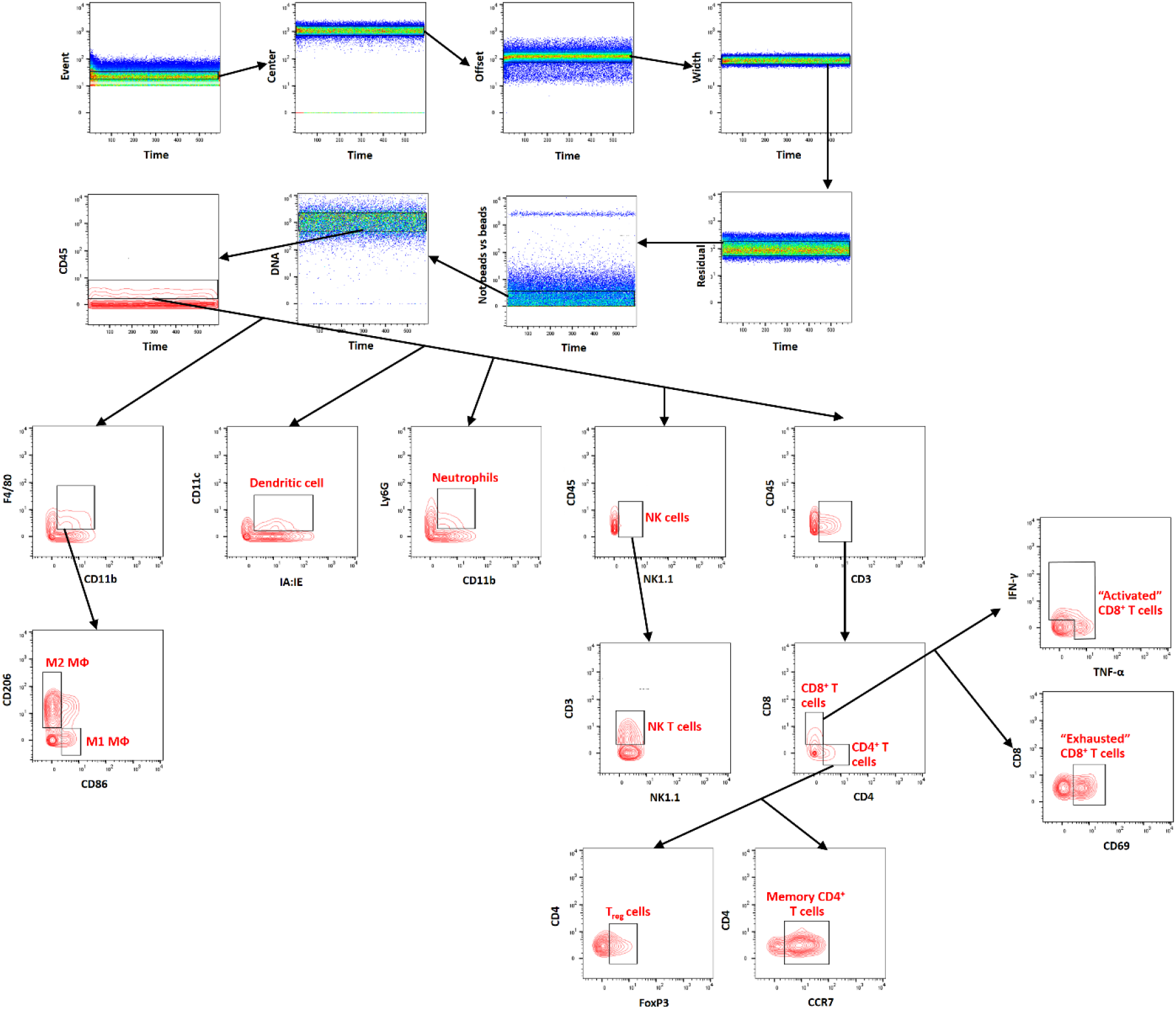
Gating strategy for immune profiling of immune cells at the colonic lamina propria through flow mass cytometry.

**Supplementary Fig. 32.**
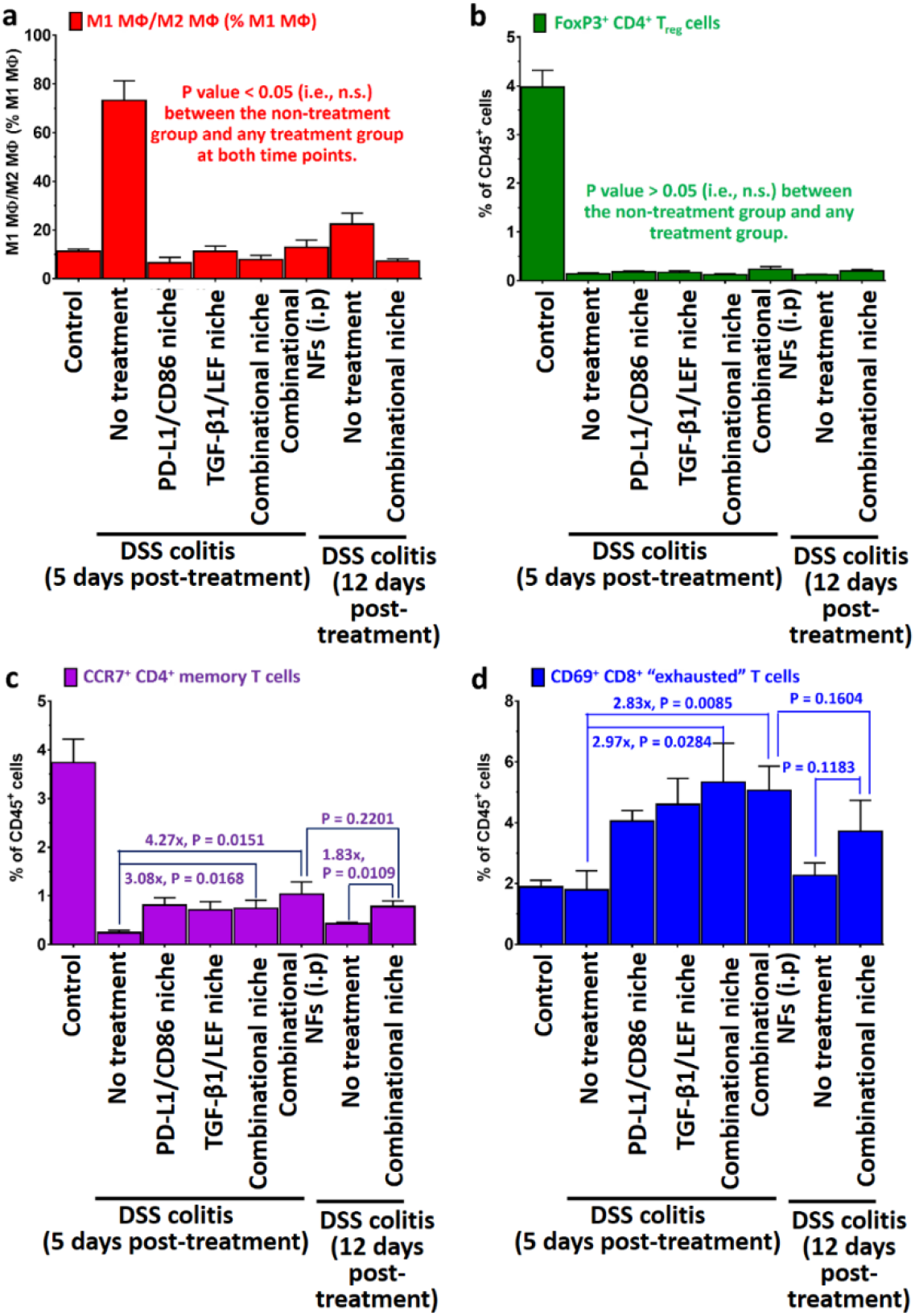
Combinational colon-specific immune niche and combinational immunosuppressive nanofibers treatment significantly increased the anti-inflammatory alternatively-activated M2 macrophages, CCR7^+^ CD4^+^ memory T cells and exhausted CD8^+^ T cells at the lamina propria. **a,** Ratio of colonic proinflammatory classically-activated M1 macrophages to anti-inflammatory alternatively-activated M2 macrophages after different colitis treatments. **b,** Frequency of pro-regulatory colonic regulatory T cells after different colitis treatments. **c,** Frequency of memory T cells after different colitis treatments. **d,** Frequency of exhausted colonic CD8^+^ T cells after different colitis treatments. (n=5) All P values were analyzed using two-way (**a,b,c,d**) ANOVA with Tukey’s HSD multiple comparisons post-hoc test.

**Supplementary Fig. 33.**
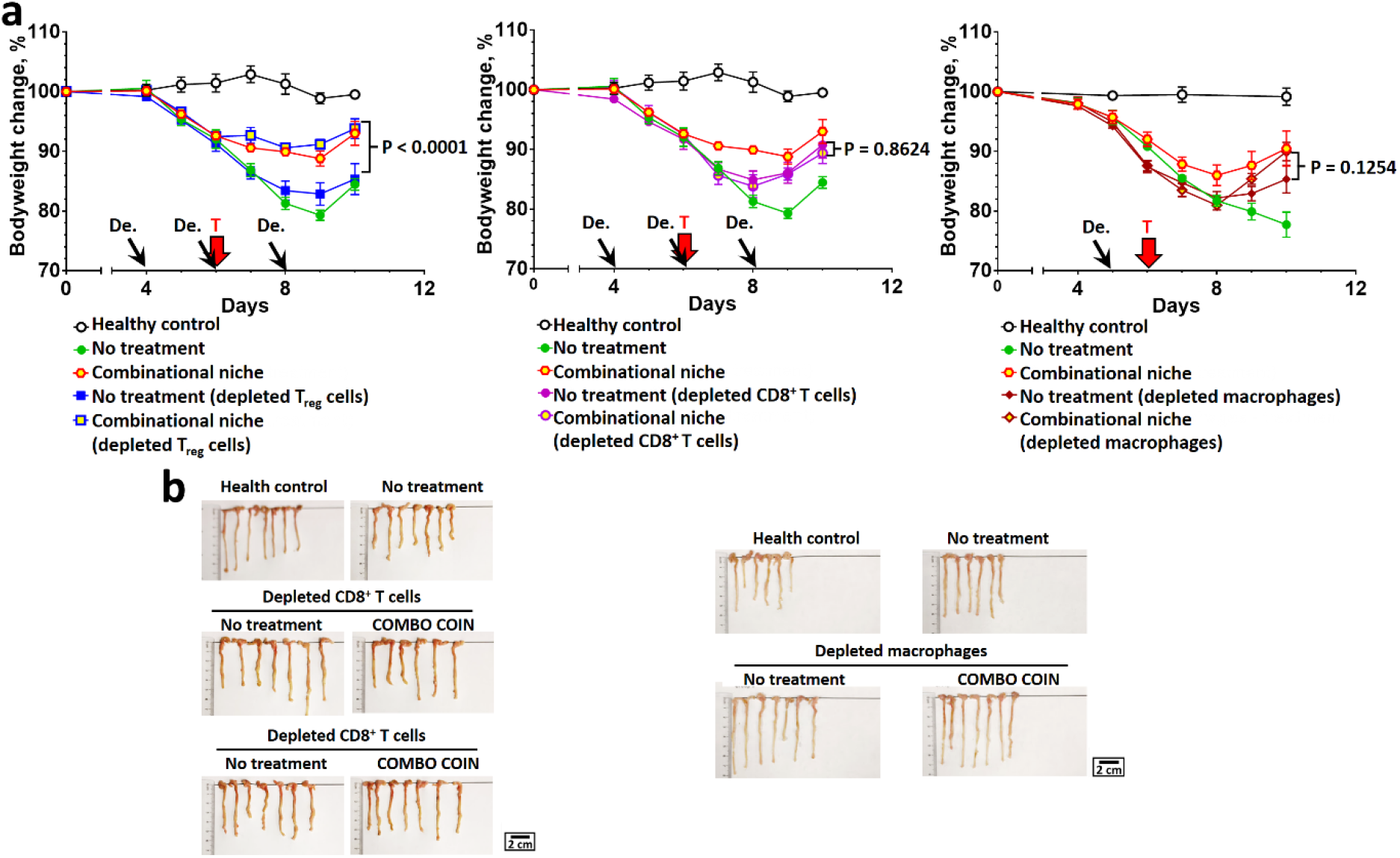
Treatment with combinational colon-specific immune niches relieves colitis symptoms through the inhibition of CD8^+^ T cell activation and macrophage polarization. **a,** Bodyweight change after treatment with combinational colon-specific immune niches in CD8^+^ T cell, regulatory T cell-, and macrophage-depleted DSS colitis mice. **b**, Digital photographs of colons preserved at the study endpoint after treatment with combinational colon-specific immune niches in CD8^+^ T cell or regulatory T cell, and macrophage-depleted colitis mice. All P values were analyzed using two-way ANOVA with Tukey’s HSD multiple comparisons post-hoc test.

**Supplementary Fig. 34.**
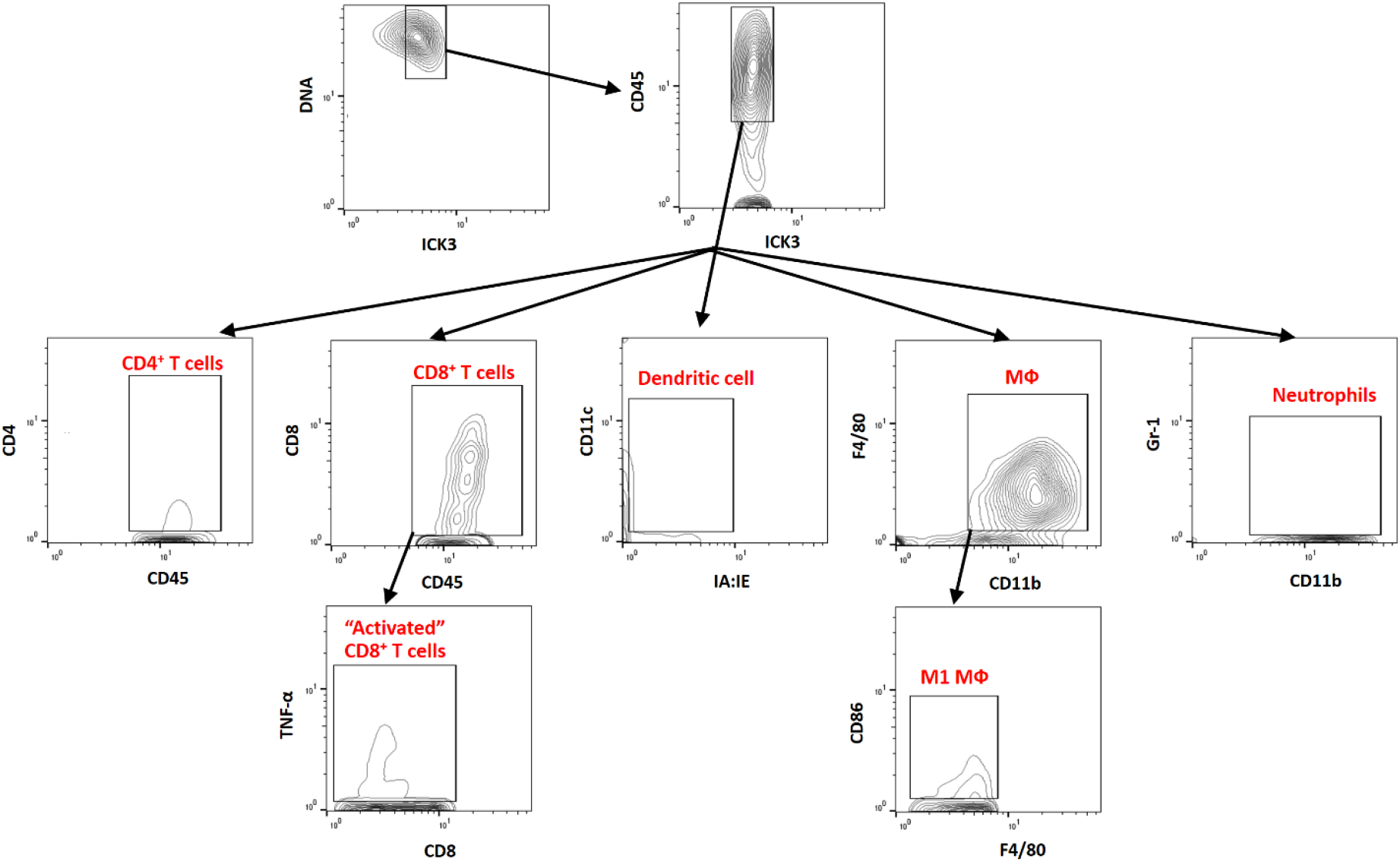
Gating strategy for immune profiling of immune cells at the subcutaneously-inoculated combinational colon-specific immune niche through imaging mass cytometry.

**Supplementary Fig. 35.**
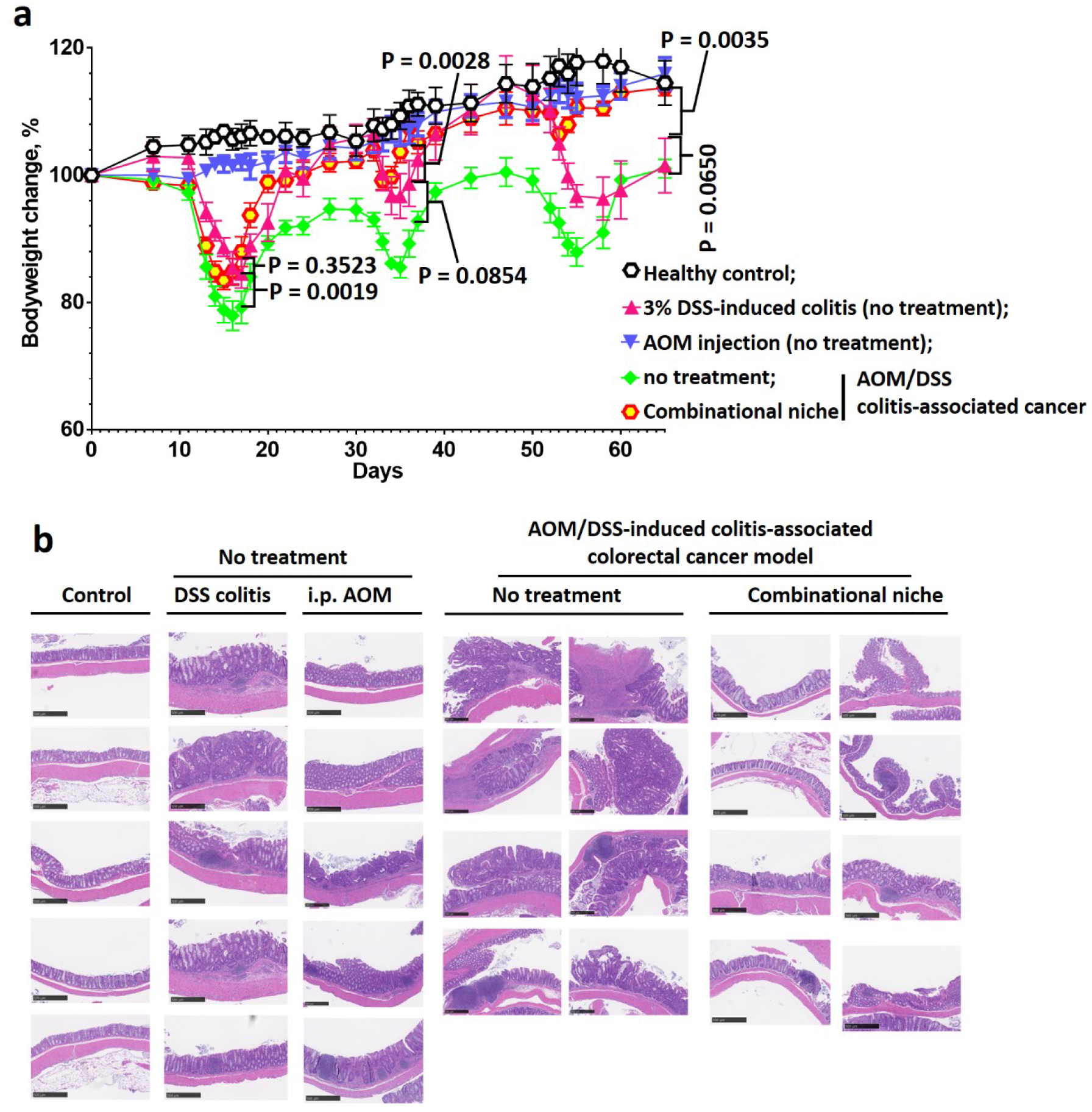
Therapeutic treatment with combinational colon-specific immune niches prevents the development of colorectal cancer. **a,** Bodyweight change after treatment with combinational colon-specific immune niches in the AOM/DSS colitis-induced colorectal cancer tumor model. **b,** H&E-stained colon sections preserved at the study endpoint (65 days after the initial study) after therapeutic treatment with combinational colon-specific immune niches in the azoxymethane/dextran sodium sulfate colitis-induced colorectal cancer tumor model.

**Supplementary Fig. 36.**
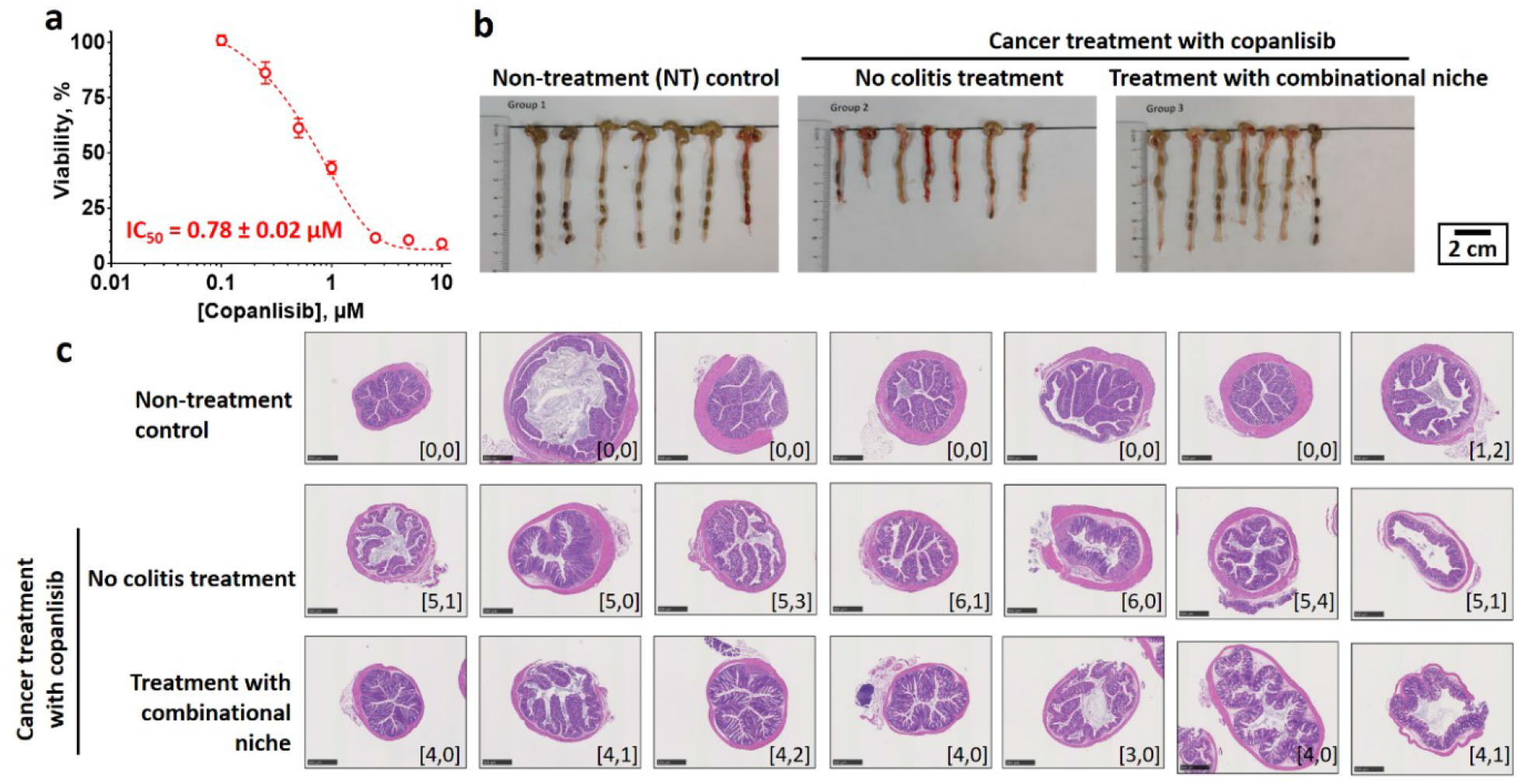
Combinational colon-specific immune niche effectively ameliorates copanlisib-induced colitis in B16OVA murine tumor model. **a,** *In vitro* toxicity of copanlisib against B16OVA cells, as determined by MTS assay. **b,** Digital photographs of colons preserved at the study endpoint. **c,** Representative H&E-stained sections of each colon and colon damage scores after different colitis treatments. The inserted label in each image represents the corresponding colon epithelial damage score and inflammatory cell infiltration score. (n = 7)

**Supplementary Fig. 37.**
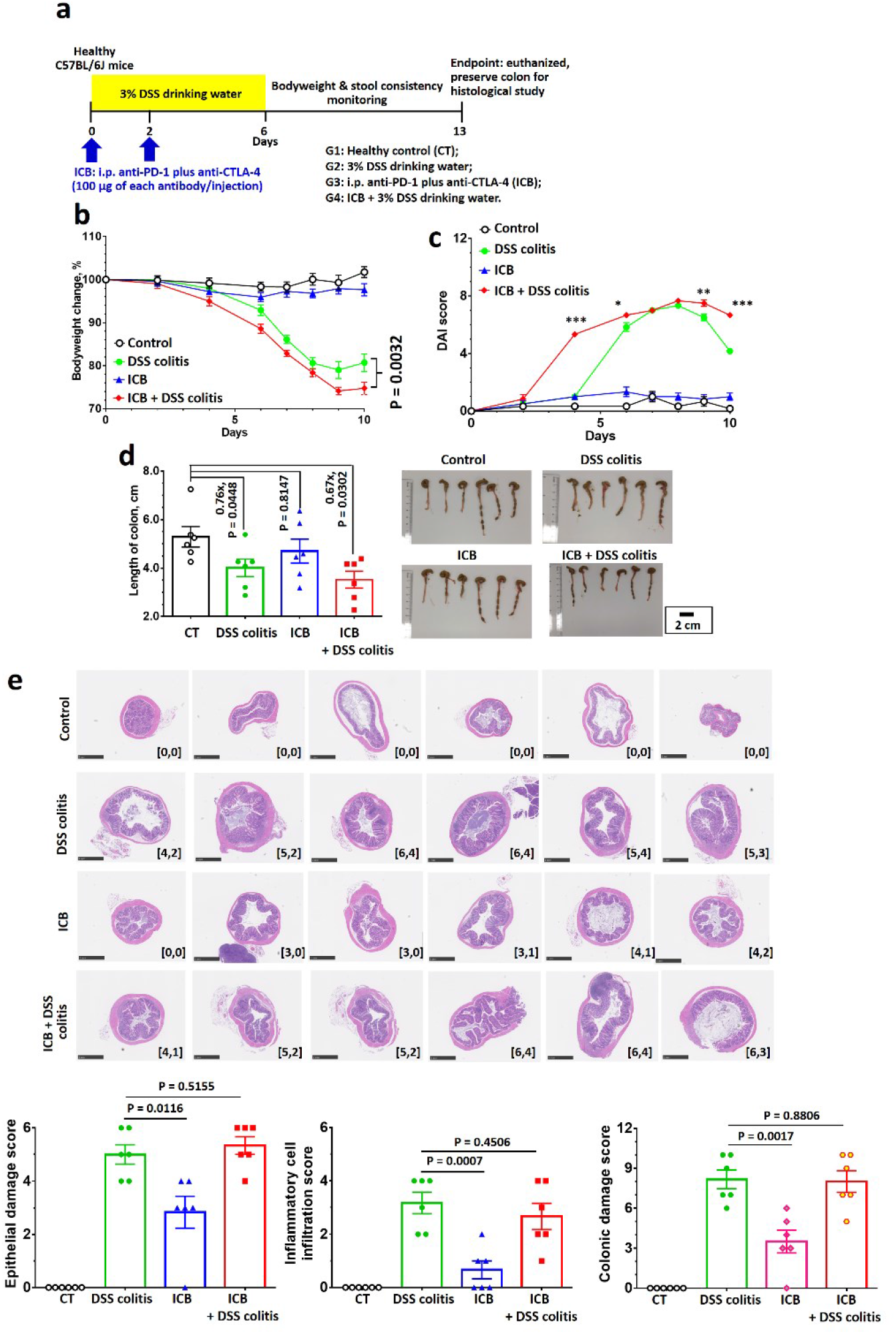
Dual immune checkpoint blockade worsens colitis symptoms in a dextran sodium sulfate-induced colitis model. **a**, Colitis induction schedule. **b-c**, Bodyweight change (**b**) and disease activity index score (**c**) recorded after colitis induction with and without immune checkpoint blockade. **d**, Digital photographs of colons and their corresponding lengths preserved at the study endpoint. **e**, H&E images of preserved colons and corresponding disease scores. Data are presented as the mean ± standard error of the mean (s.e.m.). All P values were analyzed using one-way (**d,e**) or two-way (**b,c**) ANOVA with Tukey’s HSD multiple comparisons post-hoc test.

**Supplementary Fig. 38.**
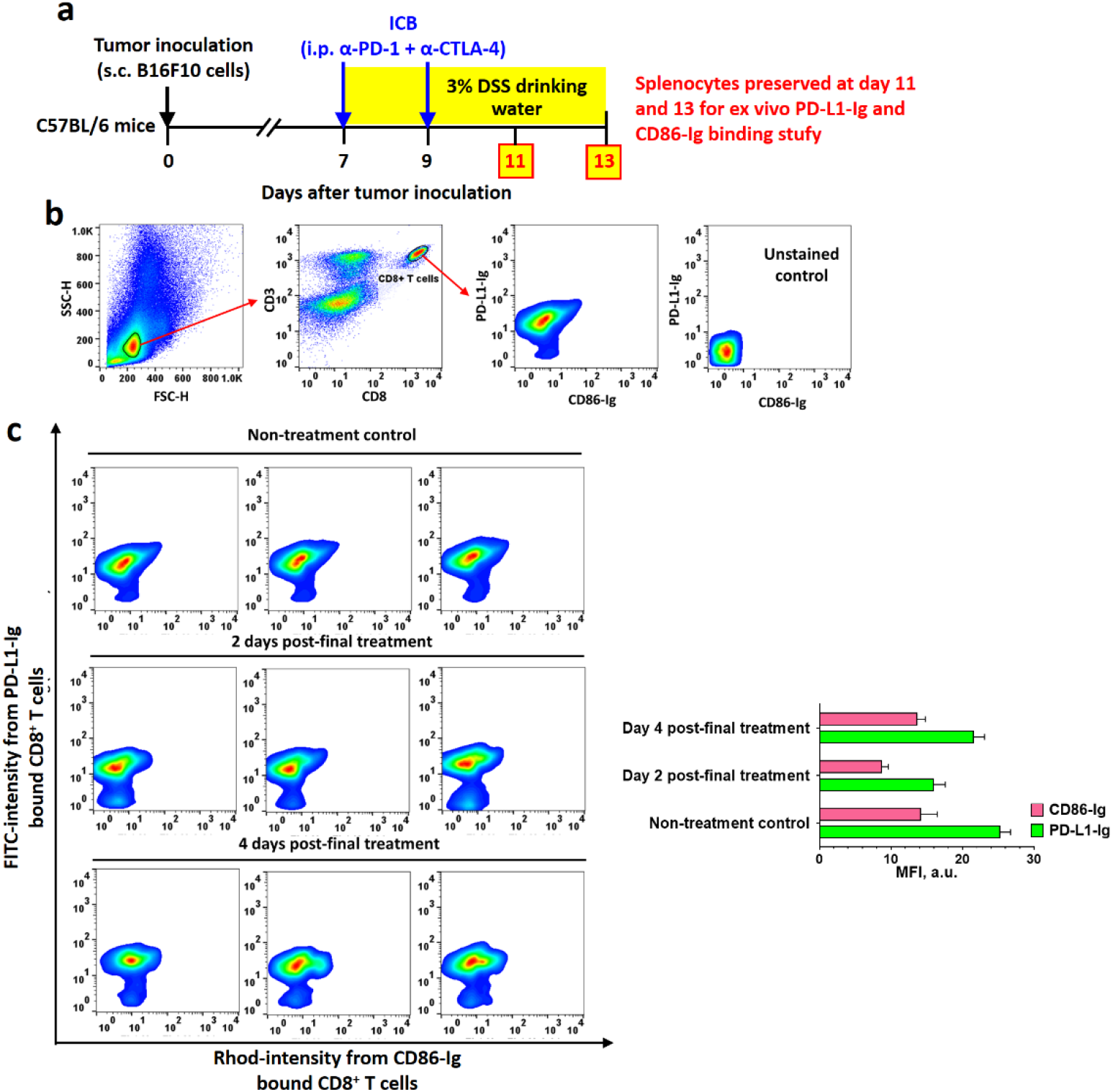
Anti-PD-1 and anti-CTLA-4 slowly detach from CD8^+^ T cells after immune checkpoint blockade treatment, allowing PD-L1 and CD86 binding to CD8^+^ T cells to induce colon-specific immunotolerance. **a**, *Ex vivo* binding study treatment schedule. **b**, fluorescence-activated cell sorting gating strategy. **c**, Individual dot scatter plots of CD8^+^ T cells after binding to FITC-labeled PD-L1-Ig and Rhod-labeled CD86-Ig.

**Supplementary Fig. 39.**
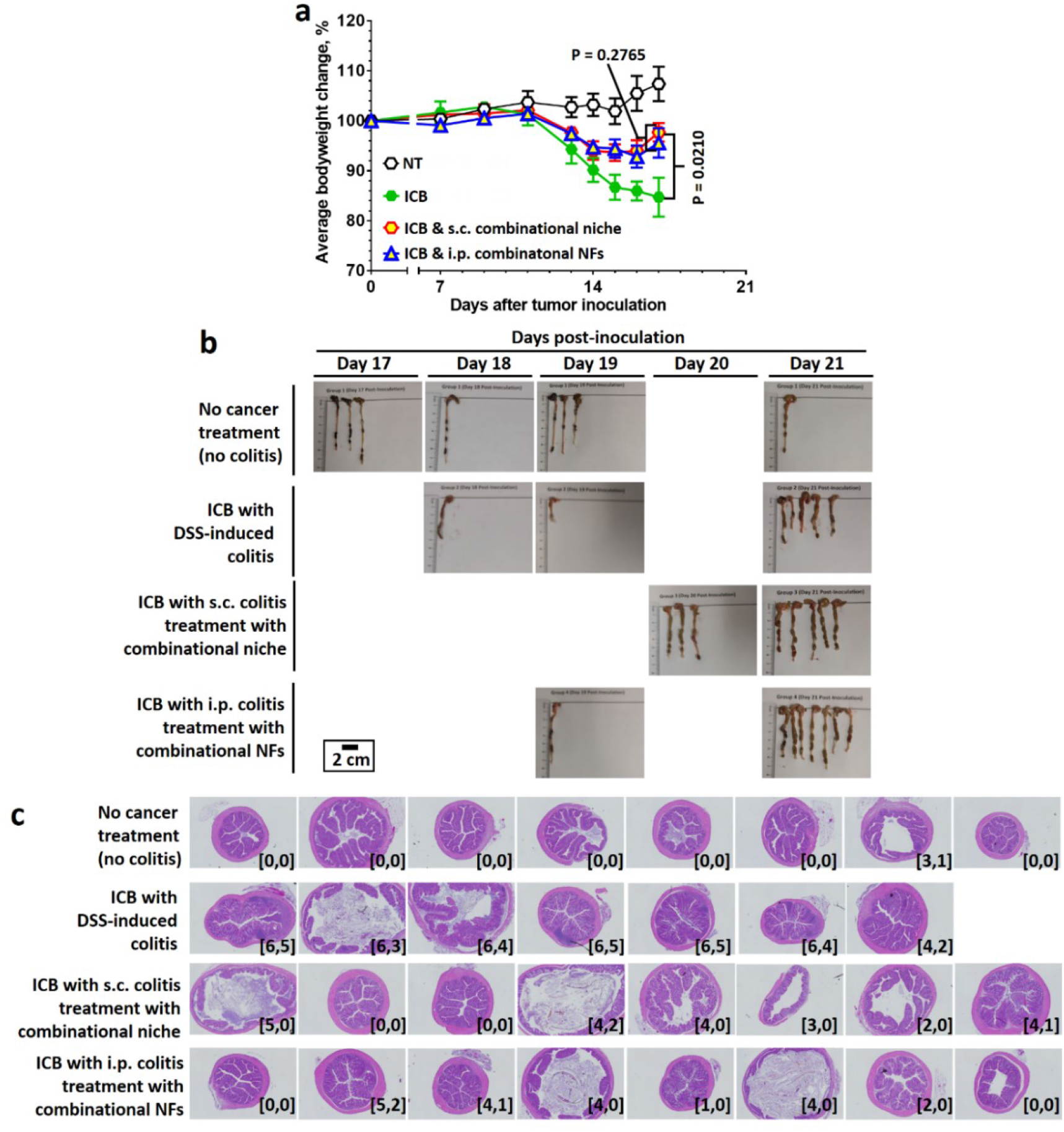
Colitis treatment with subcutaneously-inoculated combinational colon-specific immune niches or intraperitoneally-administered combinational immunosuppressive nanofibers ameliorates severe colitis symptoms triggered by dual immune checkpoint blockade in B16F10 tumor-bearing mice by reducing immune cell infiltration into the colon epithelium. **a,** Bodyweight change. **b,** Digital photographs of colons preserved at the study endpoint (23 days after the inoculation of B16F10 xenograft tumor/10 days after colitis treatment). **c,** Representative H&E-stained section of each colon preserved at the study endpoint after different treatments. The inserted label in each image represents the corresponding colon epithelial damage score and inflammatory cell infiltration score. All P values were analyzed using two-way (**a**) ANOVA with Tukey’s HSD multiple comparisons post-hoc test.

**Supplementary Fig. 40.**
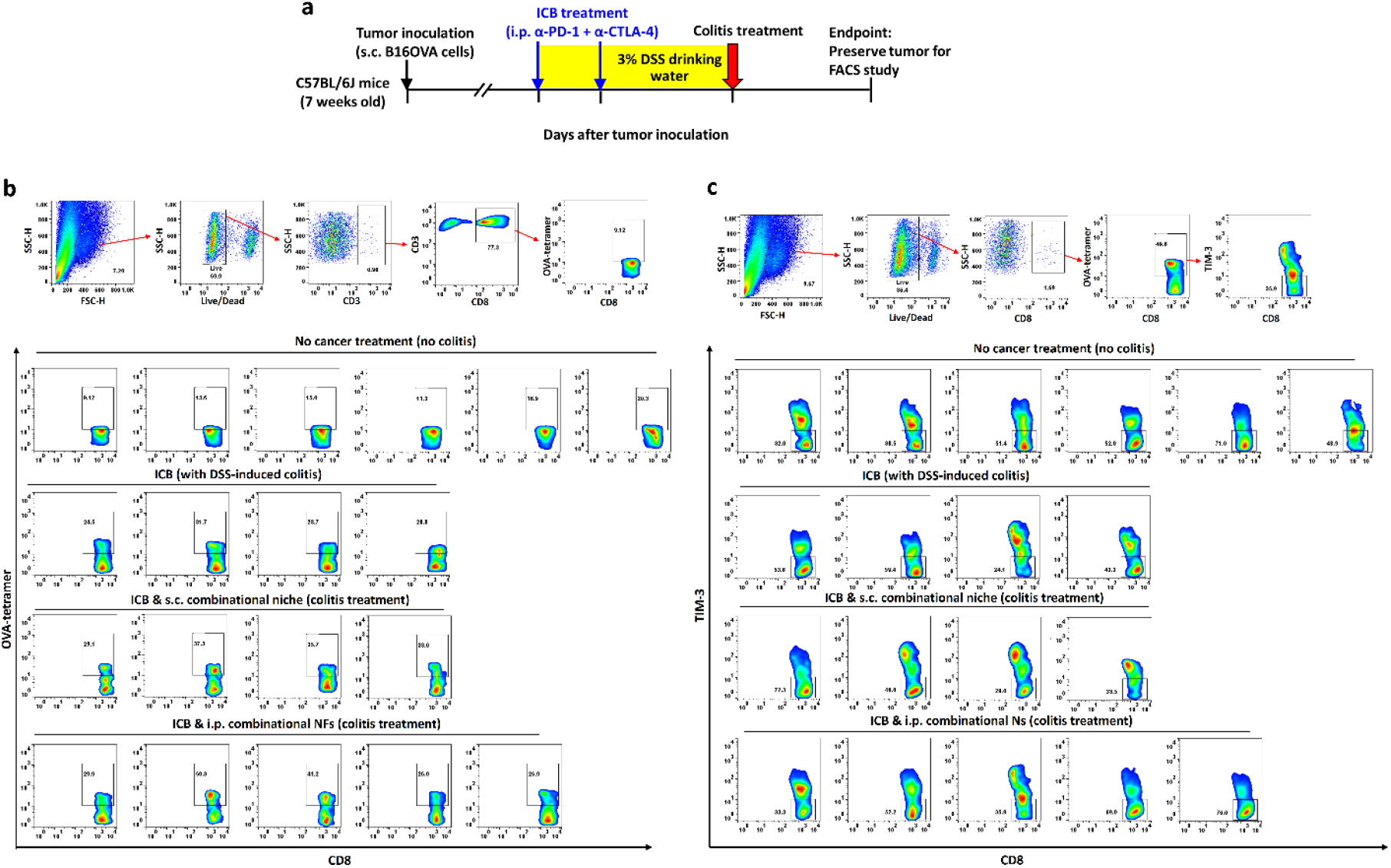
Colitis treatment with subcutaneously-inoculated combinational colon-specific immune niches or intraperitoneally-administered combinational immunosuppressive nanofibers does not exhaust tumor-specific antigen-specific CD8^+^ T cells in the B16OVA tumor model. **a**, *In vivo* treatment schedule. **b**, gating strategy and individual dot-scatter plots showing the populations of OVA-specific CD8^+^ T cells. **c**, fluorescence-activated cell sorting gating strategy and individual dot-scatter plots showing TIM-3 expression in OVA-specific CD8^+^ T cells.

**Supplementary Fig. 41.**
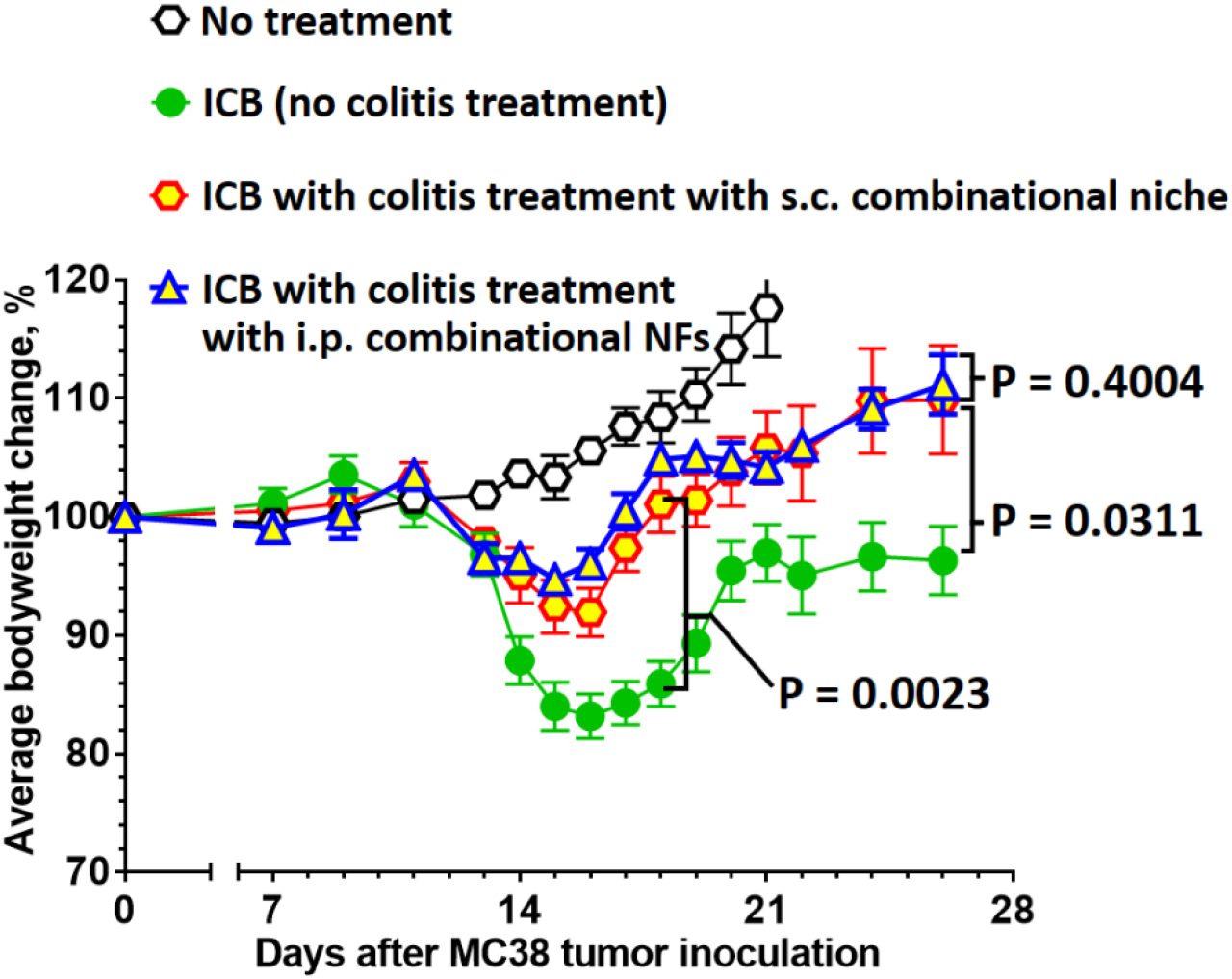
Colitis treatment with subcutaneously-inoculated combinational colon-specific immune niches or intraperitoneally-administered combinational immunosuppressive nanofibers does not affect immune checkpoint blockade cancer treatment in the MC38 tumor model. Bodyweight change after cancer and colitis treatment. All P values were analyzed using two-way ANOVA with Tukey’s HSD multiple comparisons post-hoc test.

## Extended Data for An Injectable Subcutaneous Colon-Specific Immune Niche For The Treatment Of Ulcerative Colitis

**Extended Data Fig. 1.**
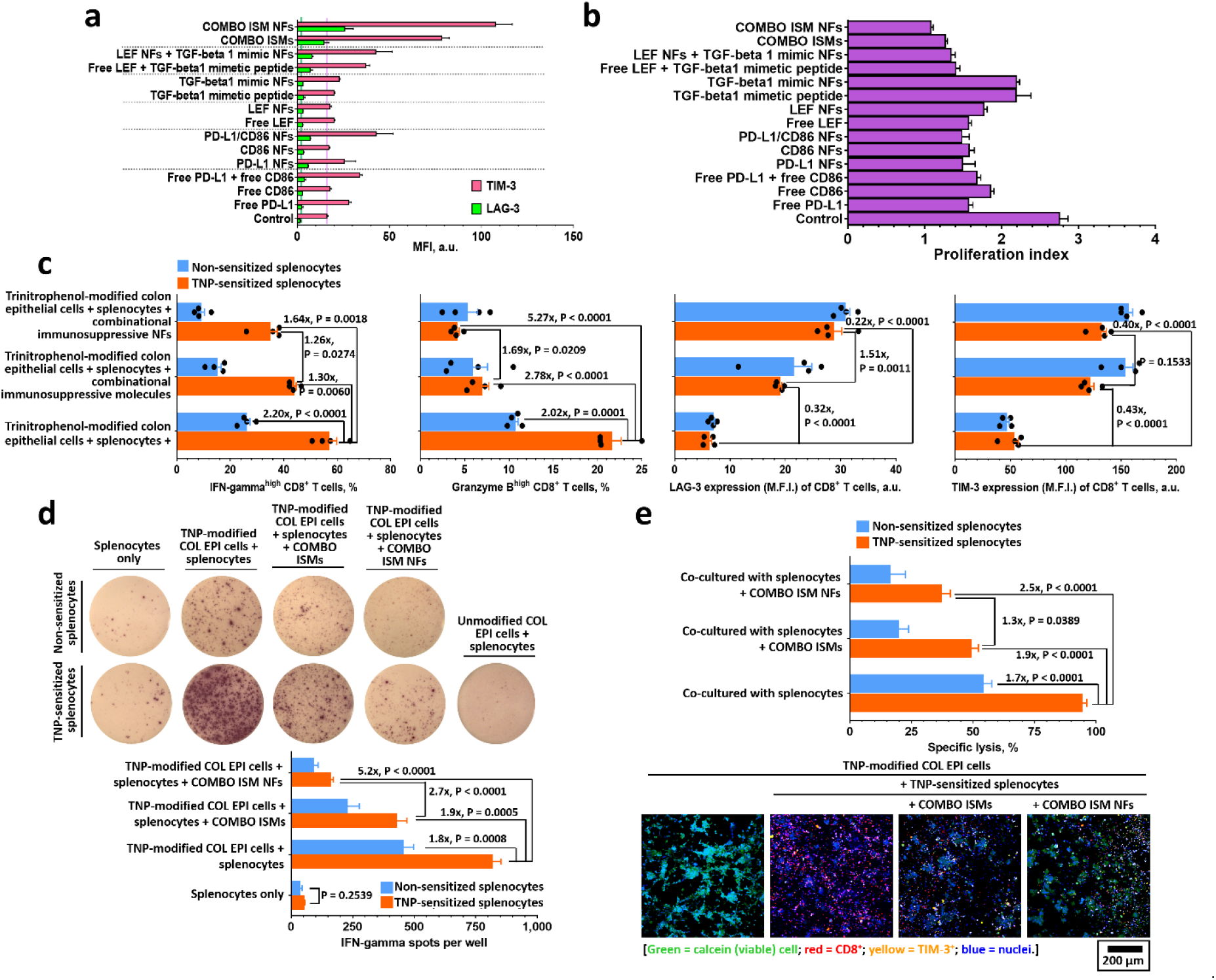
Combinational immunosuppressive nanofibers effectively inhibit antigen-specific CD8^+^ T cell activation *in vitro*. **a,** T cell inhibition (exhaustion) markers and proliferation indices of CD8^+^ T cells cultured with different functionalized NFs in the presence of anti-CD3/anti-CD28- functionalized T cell activation beads (Dynabeads) at a 1:1 ratio for 48 h, as quantified through fluorescence-activated cell sorting. The proliferation study was performed on CFSE-labeled CD8^+^ T cells. (n = 4). **b,** T cell activation and exhaustion markers on naïve and trinitrophenol-sensitized CD8^+^ T cells (in splenocytes) after co-culture with trinitrophenol-modified colon epithelial cells in the presence or absence of combinational immunosuppressive molecules or combinational immunosuppressive nanofibers (48 h). (n = 4). **c,** ELISpot assay of IFN-gamma in splenocytes after culture with unmodified or trinitrophenol-modified colon epithelial cells in the presence or absence of combinational immunosuppressive molecules or combinational immunosuppressive nanofibers (n = 4). **d,** Cytotoxicity of splenocytes against colon epithelial cells in the presence or absence of combinational immunosuppressive molecules or combinational immunosuppressive nanofibers (n = 7). Combinational nanofibers, the combination of PD-L1/CD86-functionalized nanofibers, leflunomide-encapsulated nanofibers and TGF-β1 mimetic peptide-encapsulated nanofibers; combinational immunosuppressive molecules, the combination of PD-L1/CD86-functionalized nanofibers, leflunomide-encapsulated nanofibers and TGF-β1 mimetic peptide-encapsulated nanofibers. Data are presented as the mean ± s.e.m. All P values were analyzed using two-way ANOVA with Tukey’s HSD multiple comparisons post-hoc test.

**Extended Data Fig. 2.**
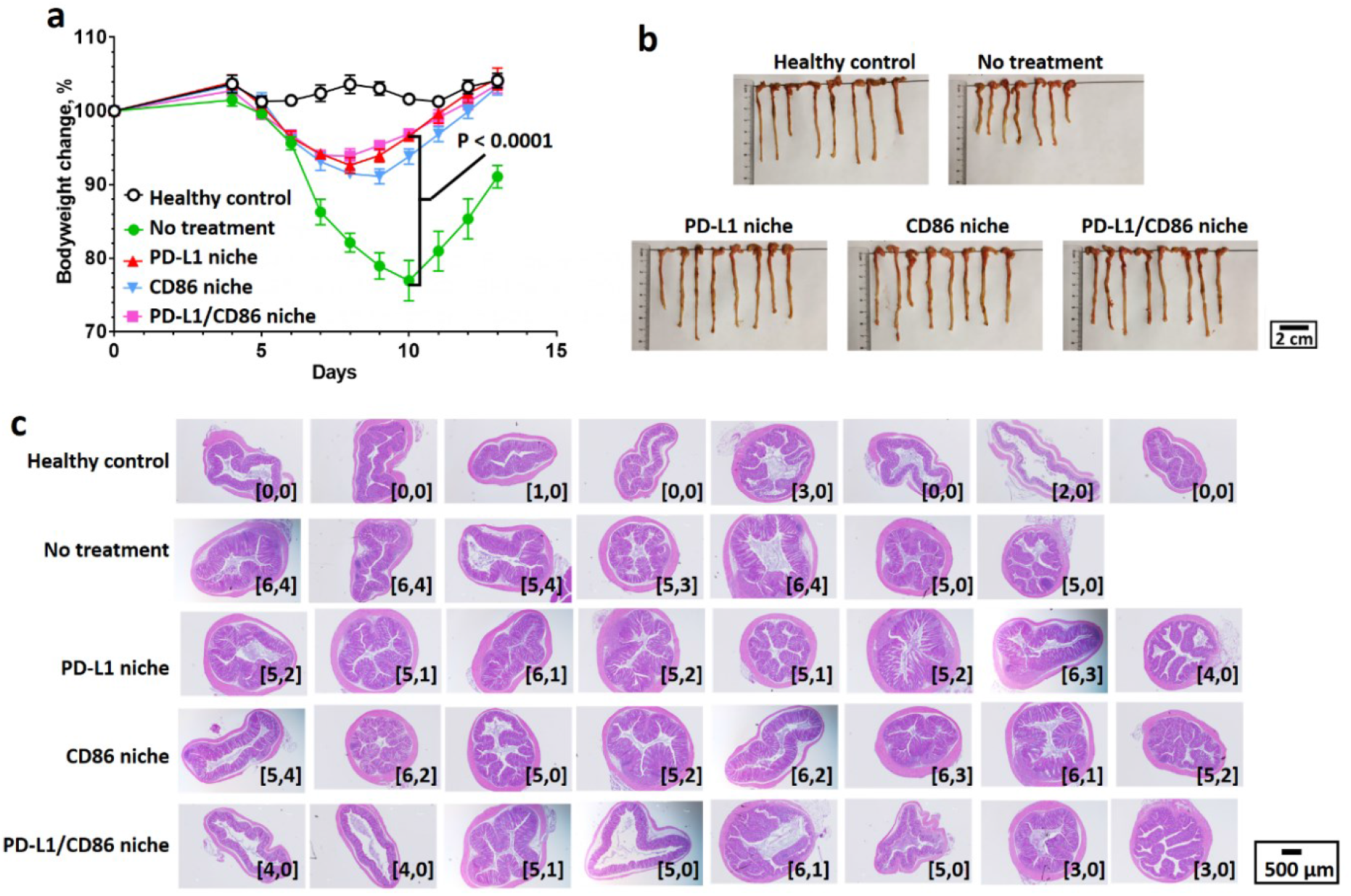
Therapeutic treatment with immune checkpoint molecule-bioengineered colon-specific immune niches prevents colitis-associated weight loss, and colon shortening and reduces immune cell infiltration into the lamina propria. **a,** Bodyweight changes throughout the study. **b,** Digital photographs of colons preserved at the study endpoint, i.e., 7 days after therapeutic treatments. **c**, Representative H&E-stained section of each colon preserved at the study endpoint after different treatments. The inserted label in each image represents the corresponding colon epithelial damage score and inflammatory cell infiltration score. P values were analyzed using one-way ANOVA (**a**) with Tukey’s HSD multiple comparisons post-hoc test.

**Extended Figure 3.**
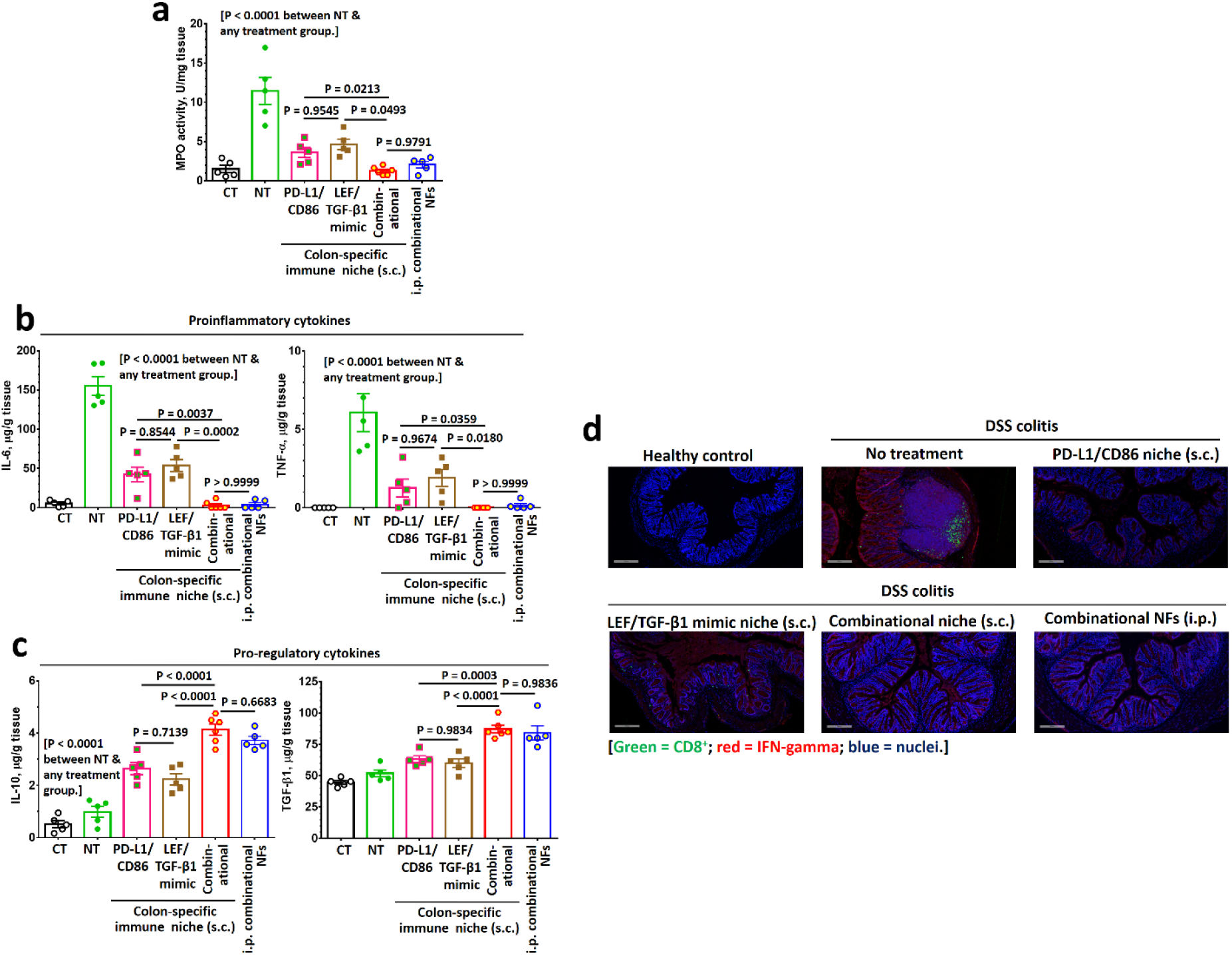
Treatment with subcutaneously-inoculated colon-specific immune niches or intraperitoneally administered combination of immunosuppressive molecule-functionalized nanofibers inhibits the production of colitis-associated myeloperoxidase and proinflammatory cytokines in the colon. **a-c**, Colonic myeloperoxidase activity (**a**), pro-inflammatory cytokine (**b**) and pro-regulatory cytokine (**c**) expression in the preserved colon specimens at the study endpoint (5 days after different therapeutic treatments) after different therapeutic treatments (n = 5). **d,** Representative immunofluorescence images of colons preserved from dextran sodium sulfate-induced colitis mice, captured 5 days after different therapeutic treatments, reveal lamina propria-infiltrated IFN-gamma^+^ CD8^+^ T cells. Data are presented as the mean ± s.e.m. All P values were analyzed using one-way ANOVA with Tukey’s HSD multiple comparisons post-hoc test.

**Extended Figure 4.**
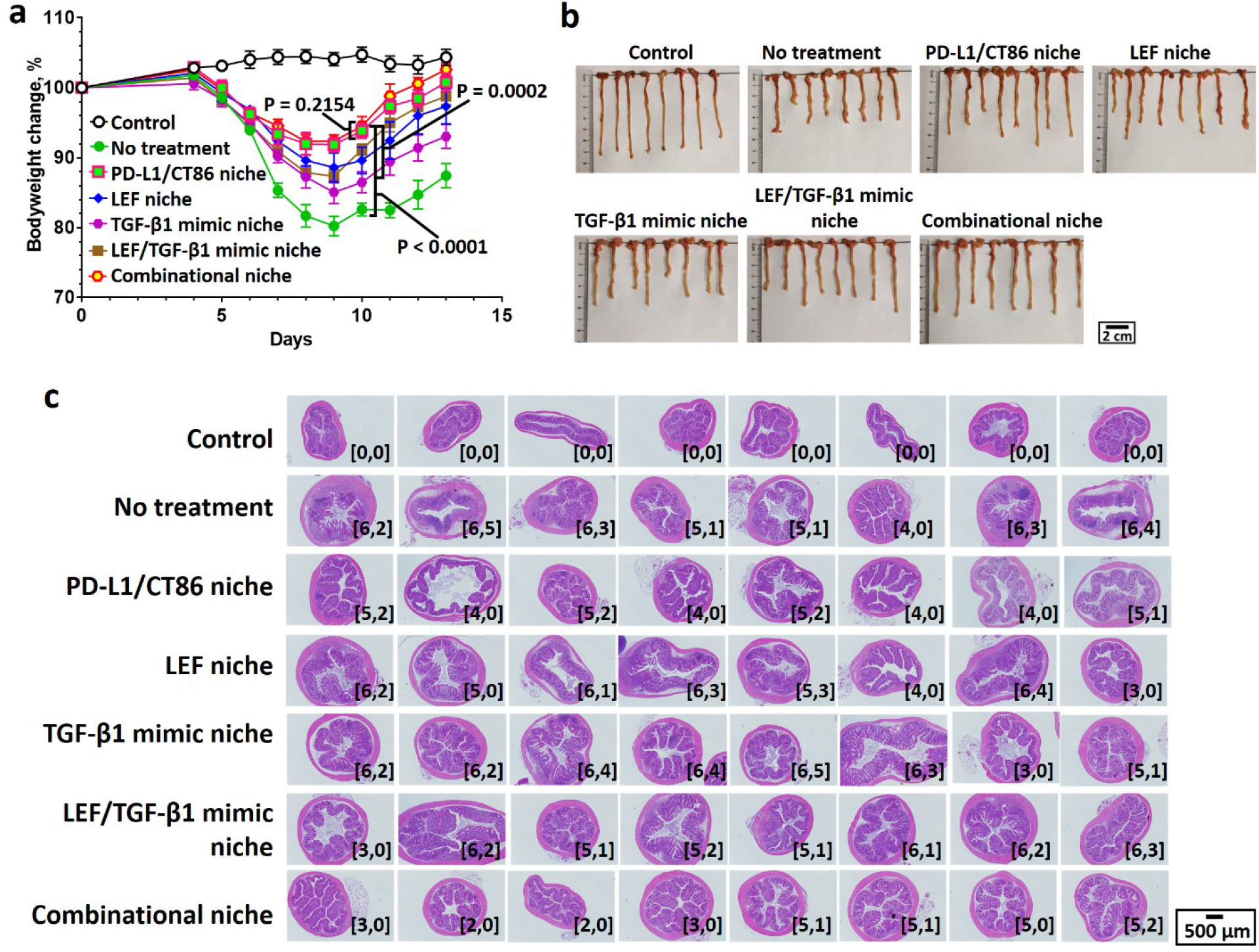
Therapeutic treatment with non-immune checkpoint molecule-functionalized colon-specific immune niches and colon-specific immune niches prevents colitis-associated colon shortening and reduces immune cell infiltration into the lamina propria. **a,** Bodyweight changes throughout the study. **b,** Digital photographs of colons preserved at the study endpoint, i.e., 7 days after different therapeutic treatments. **c**, Representative H&E-stained section of each colon preserved at the study endpoint after different treatments. The inserted label in each image represents the corresponding colon epithelial damage score and inflammatory cell infiltration score. P values were analyzed using one-way ANOVA (**a**) with Tukey’s HSD multiple comparisons post-hoc test.

**Extended Figure 5.**
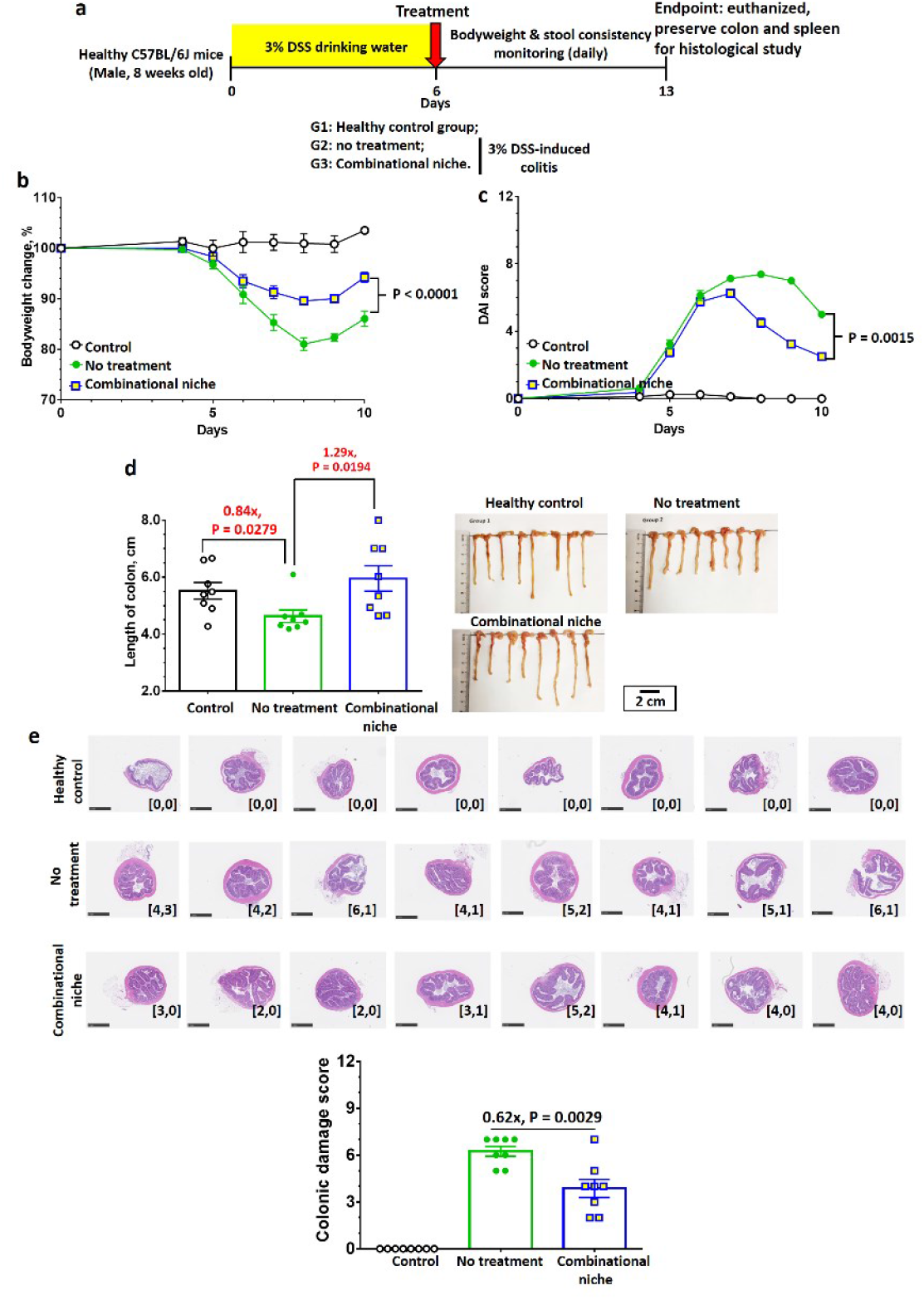
Colon-specific immune niches effectively ameliorate DSS-induced colitis in male C57BL/6 mice. **a-d**, Evaluation of the therapeutic treatment efficiency of combinational colon-specific immune niches in the dextran sodium sulfate-induced-induced colitis mouse model. Therapeutic treatment schedule (**a**): colitis mice received a single subcutaneous treatment with the combo immune niche at day 6. **b-c,** Bodyweight change (**b**) and disease activity index score (**c**) were monitored for up to 12 days. **d**, Digital images and length of colon preserved at the study endpoint. **e**, H&E-stained sections of each colon preserved at the study endpoint and their disease scores after therapeutic treatment with the combo immune niche. The inserted label in each image represents the corresponding colon epithelial damage score and inflammatory cell infiltration score. Data are presented as the mean ± standard error of the mean (s.e.m.). All P values were analyzed using one-way (**d,e**) or two-way (**b,c**) ANOVA with Tukey’s HSD multiple comparisons post-hoc test.

**Extended Figure 6.**
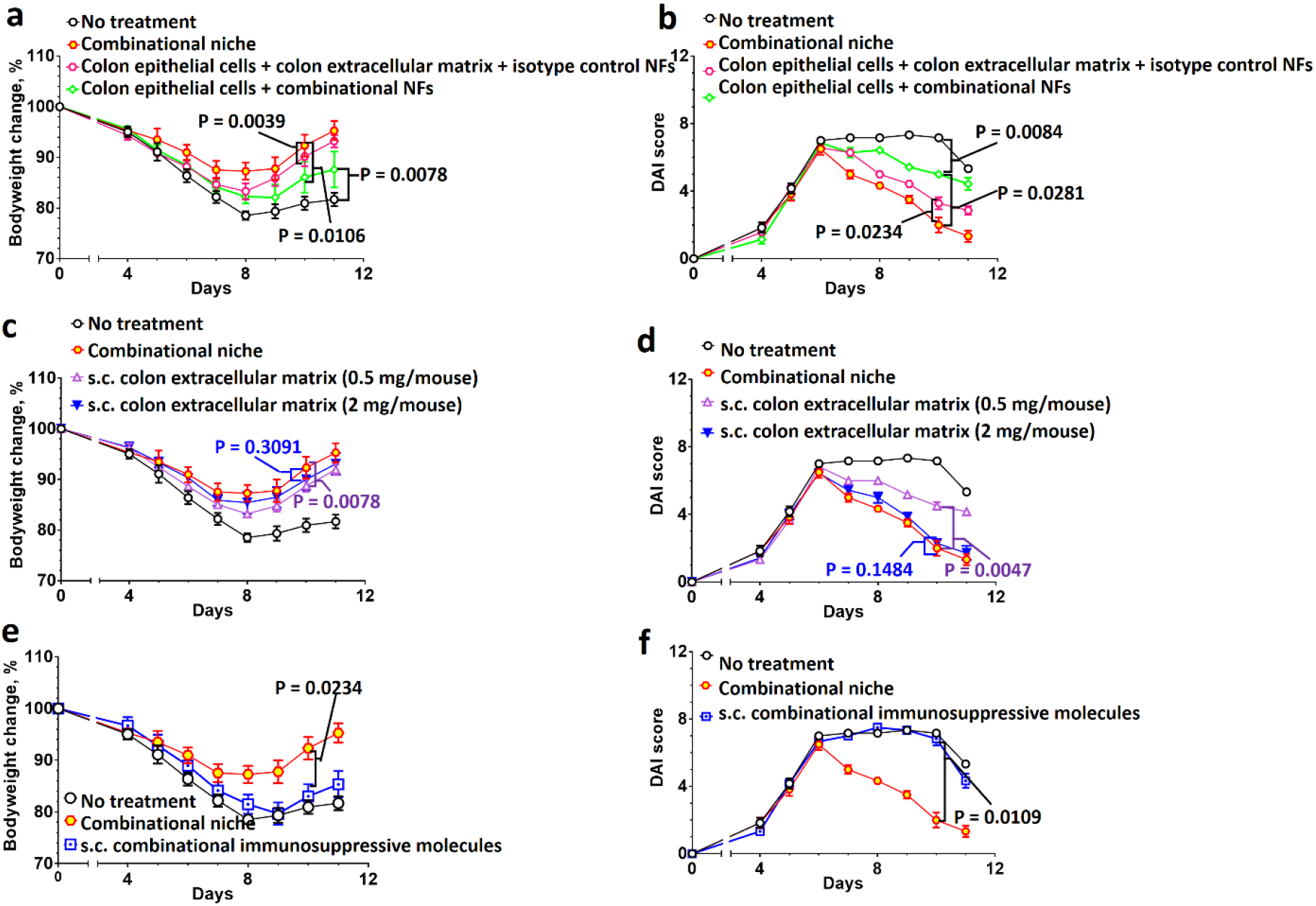
The combination of colon epithelial cells and combinational immunosuppressive nanofibers, or colon extracellular matrix alone are not as effective as combinational colon-specific immune niche (the combination of colon epithelial cells, colon extracellular matrix, combinational immunosuppressive nanofibers) in relieving the colitis symptoms in a dextran sodium sulfate-induced colitis model. Bodyweight changes (**a,c,e**) and DAI scores (**b,d,f**) after different control colitis treatments with different control colon-specific immune niches. Statistical analysis was performed 4 days after different colitis treatments. (n = 6 or 7) Data are presented as the mean ± s.e.m. All P values were analyzed using two-way ANOVA with Tukey’s HSD multiple comparisons post-hoc test.

**Extended Figure 7.**
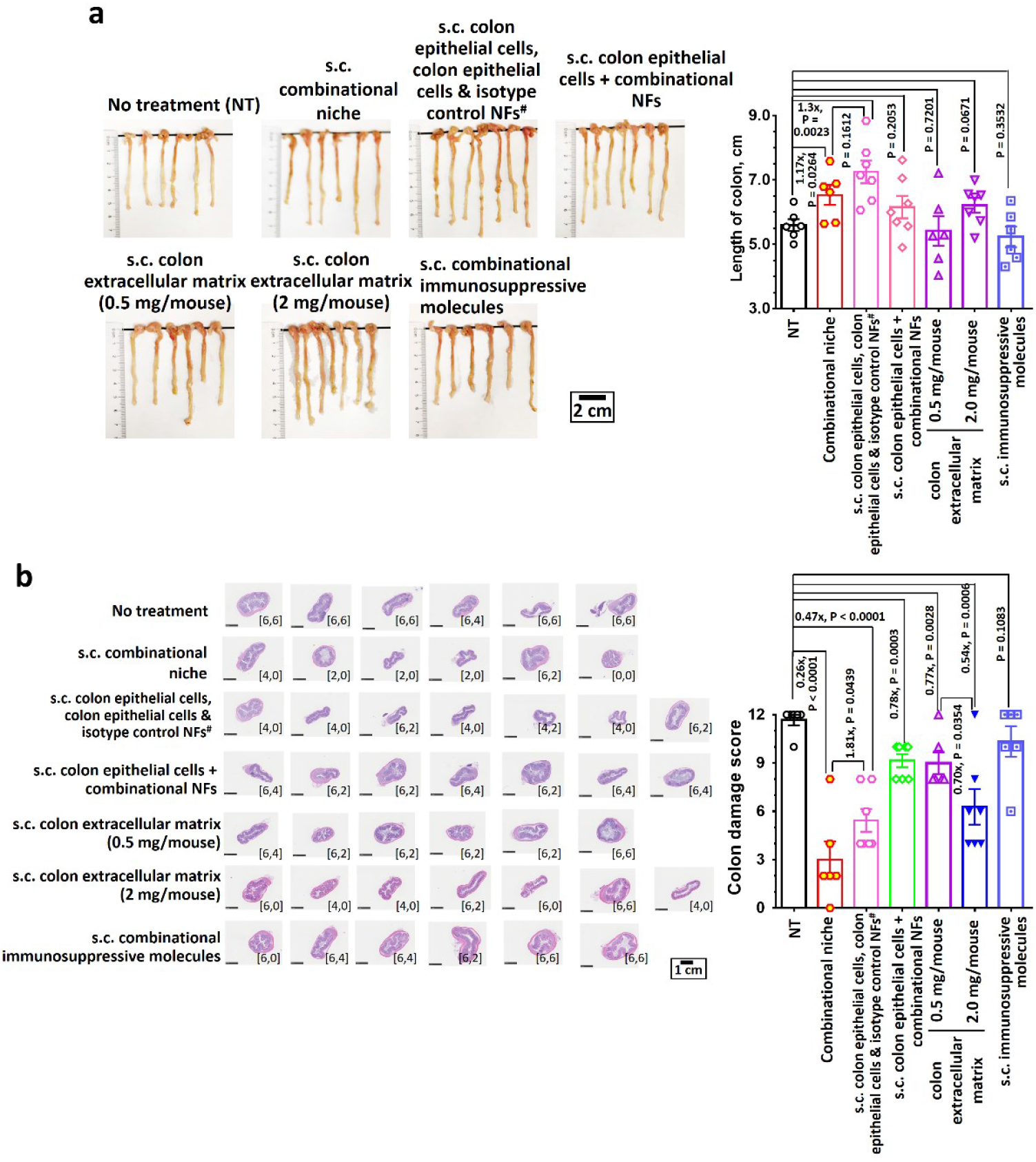
The combination of colon epithelial cells and combinational immunosuppressive nanofibers, or colon extracellular matrix alone are not as effective as colon-specific immune niches (the combination of colon epithelial cells, colon extracellular matrix, combinational immunosuppressive nanofibers) in preventing inflammation-associated colon shortening, colitis-induced colon epithelial damage and immune cell infiltration into the colon in the dextran sodium sulfate colitis model. **a,** Digital photographs and the lengths of colons preserved at the study endpoint after different colitis treatments. **b,** Representative H&E-stained sections of each colon and colon damage scores after different colitis treatments. The inserted label in each image represents the corresponding colon epithelial damage score and inflammatory cell infiltration score. (n = 6 or 7) All P values were analyzed using one-way (**a**) or two-way (**b**) ANOVA with Tukey’s HSD multiple comparisons post-hoc test.

**Extended Figure 8.**
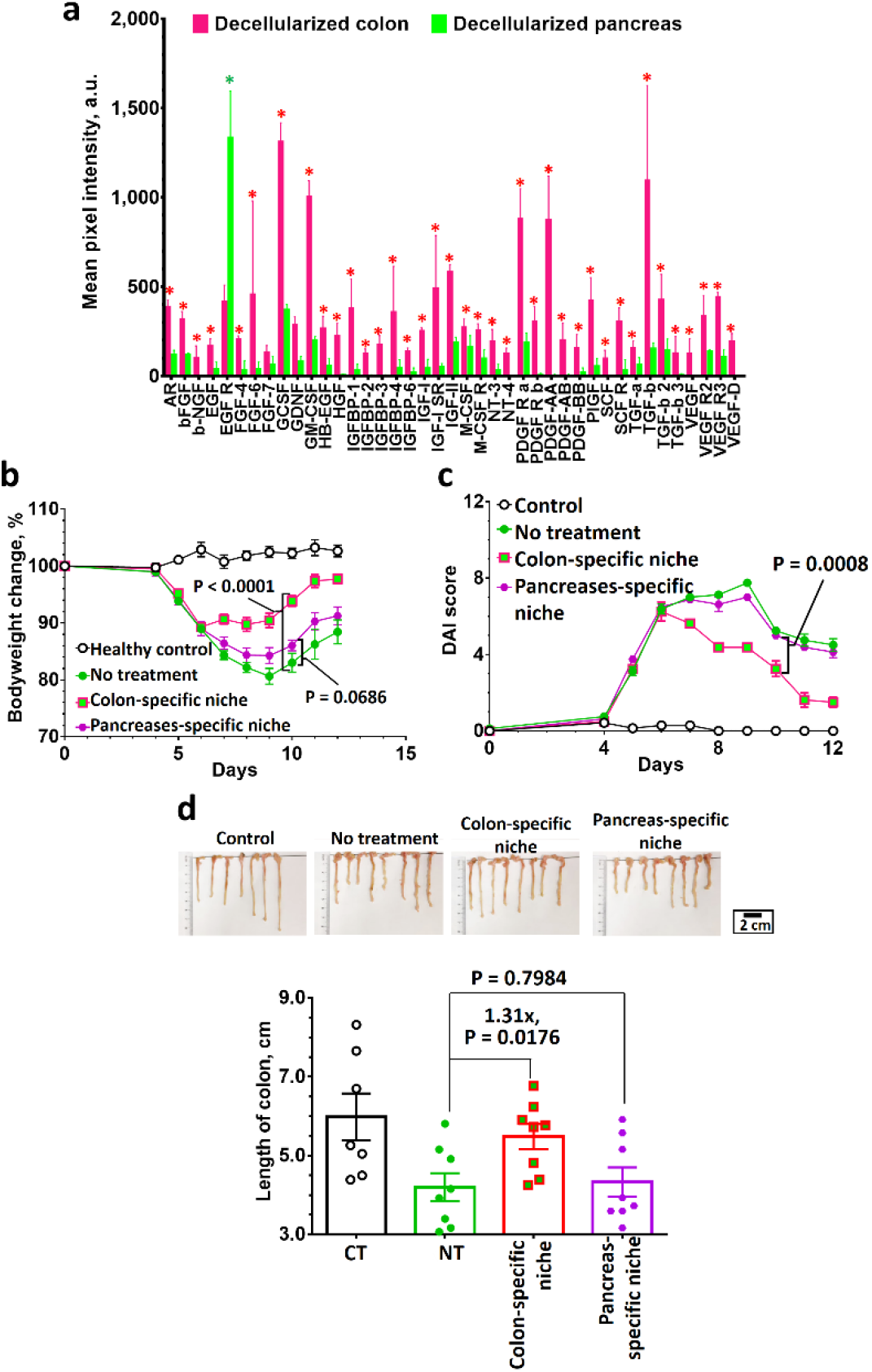
Pancreas-specific combinational immune niche cannot relieve colitis symptoms in dextran sodium sulfate colitis mice. **a,** Analysis of growth factors and cytokines present in decellularized colon extracellular matrix and decellularized mouse pancreas *via* sandwich-type antibody microarray assay. (n = 4 biologically independent samples) **b-c**, Bodyweight change (**b**) and disease activity index score (**c**) after colitis treatment with combinational colon-specific immune niche or combinational pancreas-specific immune niche. **d,** Digital photographs of colons preserved at the study endpoint (5 days after therapeutic treatment), and their lengths (**d**). (n = 7 or 8) All P values were analyzed using one-way (**d**) or two-way (**a,b,c**) ANOVA with Tukey’s HSD multiple comparisons post-hoc test.

**Extended Figure 9.**
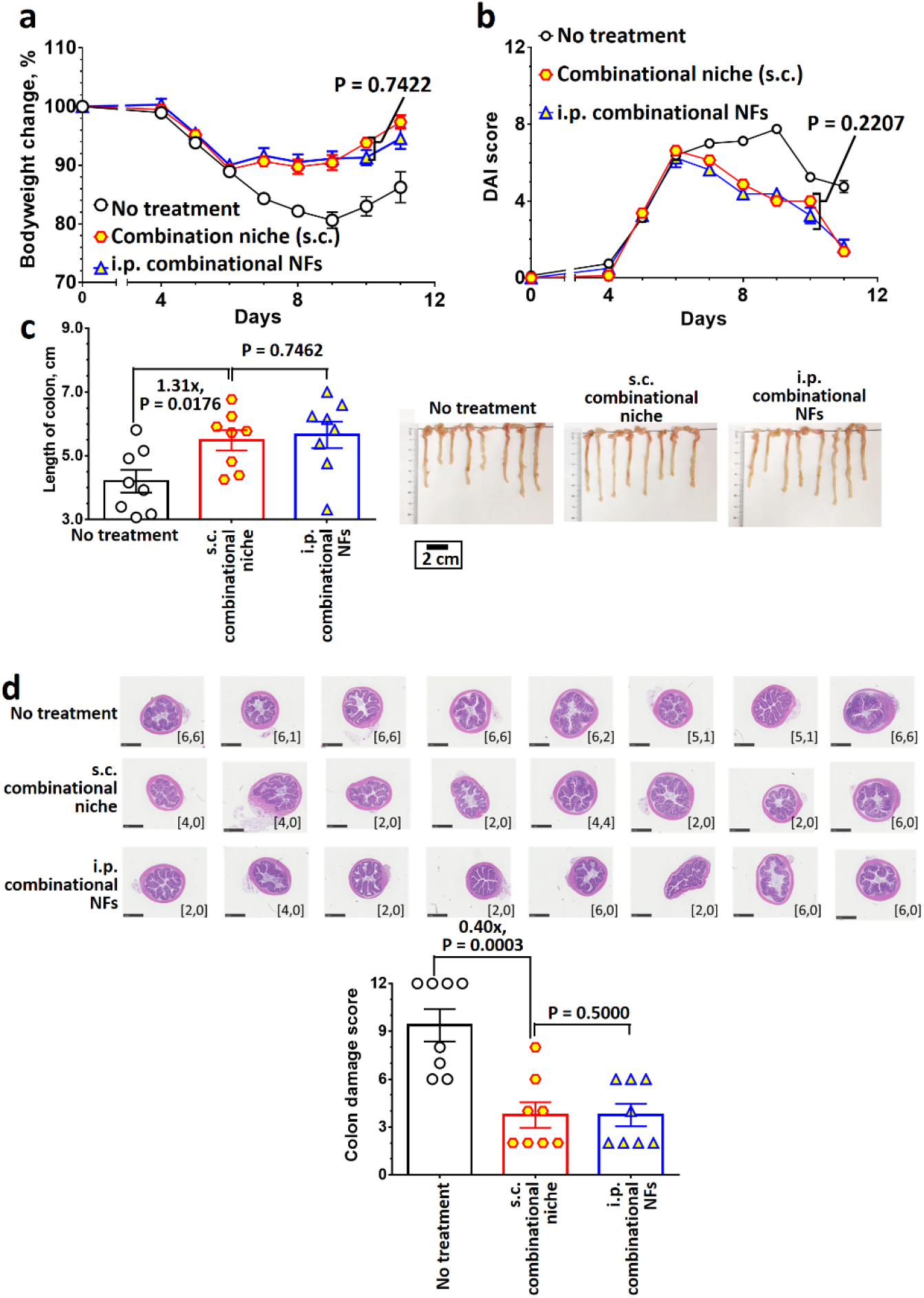
Intraperitoneal-administered combinational immunosuppressive nanofibers are as effective as subcutaneously-inoculated combinational colon-specific immune niche to treat dextran sodium sulfate-induced colitis. **a-b,** Bodyweight change (**a**) and disease activity index score (**b**) after colitis treatment with subcutaneously-administered combinational colon-specific immune niche or intraperitoneally-administrated combinational immunosuppressive nanofibers. **c,** Digital photographs of colons preserved at the study endpoint (5 days after therapeutic treatment), and their lengths. **d,** Representative H&E-stained sections of each colon and colon damage scores after different colitis treatments. The inserted label in each image represents the corresponding colon epithelial damage score and inflammatory cell infiltration score. (n = 8) All P values were analyzed using one-way (**c**) or two-way (**a,b,d**) ANOVA with Tukey’s HSD multiple comparisons post-hoc test.

**Extended Figure 10.**
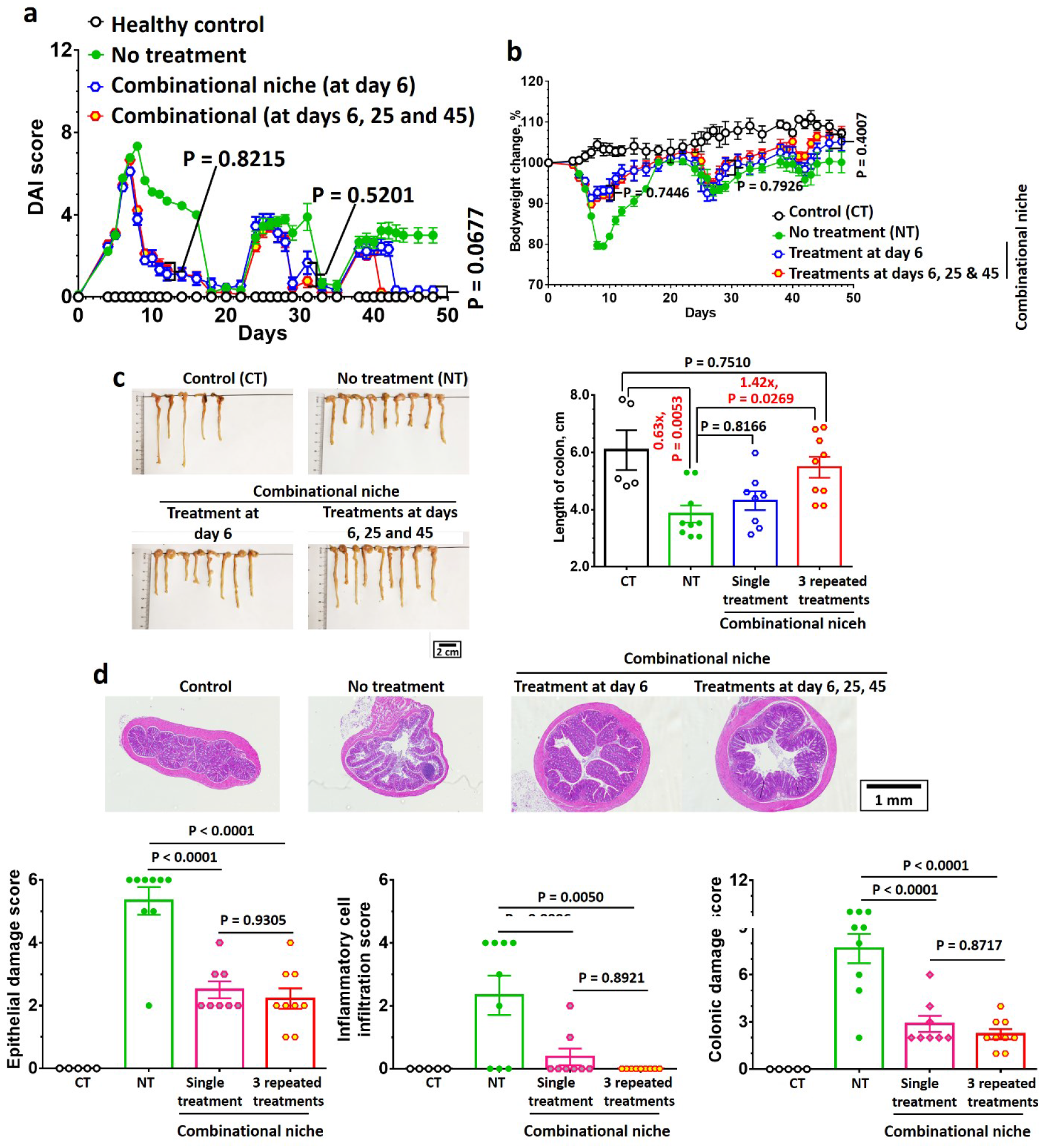
Single therapeutic treatment with combinational colon-specific immune niches effectively ameliorates colitis symptoms in a chronic dextran sodium sulfate-induced colitis model. **a-b,** Disease activity index score (**a**) and bodyweight (**b**) change after treatment(s) with combinational colon-specific immune niche. **b**, Digital photographs and length of colons preserved at the study endpoint (48 days after the initial colitis induction) after one or three therapeutic treatments with the combo immune niche. **d**, Representative H&E-stained images, colon epithelial damage score, inflammatory cell infiltration score, and colonic damage score of colons preserved at the study endpoint (48 days after the first induction of colitis) after one or three therapeutic treatments with combinational colon-specific immune niche.

**Supplementary Table 1.** Characterization of different poly(2-(methacryloyloxy)ethyl phosphorylcholine)-poly(hydroxypropyl methacrylate) (PMPC-PHPMA) diblock copolymers with different PMPC and PHPMA degrees of polymerization synthesized at different solid contents, as determined by dynamic light scattering (DLS) and transmission electron microscopy (TEM). (Note: ^ǁ^ sphere-equivalent Dh determined by DLS. All DLS measurements were based on the average of 3 independent measurements. All measurements were measured at 25 °C. ^ determined by TEM.)

| | Average intensity-average diameter ($D_h$), nm (average polydispersity index (PDI))<sup>l</sup> | Morphology<sup>^</sup> |
| --- | --- | --- |
| **PMPC27-PHPMA140**<br>(prepared at 25wt/wt % solid content) | $28 \pm 1$ ($0.080 \pm 0.006$) | Spheres |
| **PMPC27-PHPMA210**<br>(prepared at 25wt/wt % solid content) | $47 \pm 1$ ($0.175 \pm 0.008$) | Spheres + short nanofibers |
| **PMPC27-PHPMA280**<br>(prepared at 25wt/wt % solid content) | $490 \pm 2$ ($0.532 \pm 0.053$) | Nanofibers |
| **PMPC27-PHPMA280**<br>(prepared at 5wt/wt % solid content) | $66 \pm 1$ ($0.049 \pm 0.016$) | Spheres |
| **PMPC27-PHPMA350**<br>(prepared at 25wt/wt % solid content) | $597 \pm 3$ ($0.483 \pm 0.063$) | Nanofibers |
| **PMPC27-PHPMA420**<br>(prepared at 25wt/wt % solid content) | $320 \pm 4$ ($0.197 \pm 0.021$) | Nanofibers + vesicles |

**Supplementary Table 2.** Characterization of unmodified and different functionalized PMPC27-PHPMA280-based nanofibers (NFs) by dynamic light-scattering and aqueous electrophoresis methods. (Note: ^#^ Average number-average diameter (Dn); ^ǁ^ sphere-equivalent Dh; ^ all samples, unless specified, were dispersed in 1X PBS, and diluted 100 times with deionized water before dynamic light-scattering and aqueous electrophoresis studies. The pH of the background dispersion was 6.8 – 8.0. All measurements were based on the average of three independent measurements. All measurements, unless specified, were measured at 25 °C.)

|  | Average intensity-average diameter (D<sub>h</sub>), nm (average polydispersity index (PDI))^ | Average zeta potential, mV^ |
| --- | --- | --- |
| **Unmodified (COOH-functionalized) NFs** | 442 ± 7 (0.427 ± 0.007)<sup>l</sup>;<br>489 ± 6 (0.437 ± 0.012) at 4 °C in 1X PBS;<br>456 ± 5 (0.389 ± 0.012) at 25 °C in 1X PBS;<br>454 ± 9 (0.367 ± 0.009) at 37 °C in 1X PBS;<br>428 ± 8 (0.356 ± 0.016) at 45 °C in 1X PBS. | - 24.9 ± 0.9 |
| **DBCO-functionalized NFs** | 474 ± 10 (0.384 ± 0.067)<sup>l</sup> | - 14.1 ± 2.9 |
| **1:1 mixture of PD-L1-Ig /CD86-Ig** | 13 ± 2<sup>#</sup> | - 42.3 ± 2.9 |
| **PD-L1-Ig /CD86-Ig dual-functionalized NFs (PD-L1/CD86 NFs)** | 470 ± 20 (0.333 ± 0.052)<sup>l</sup> | - 23.0 ± 3.6 |
| **TGF-β1 mimetic peptide** | Not applicable | + 22.1 ± 3.1 |
| **TGF-β1 mimetic peptide-functionalized NFs (TGF-β1 mimic NFs)** | 503 ± 24 (0.714 ± 0.039)<sup>l</sup> | - 2.6 ± 4.2 |
| **LEF NPs** | 107 ± 1 (0.092 ± 0.003) | - 23.9 ± 0.5 |
| **LEF NP-functionalized NFs (LEF NFs)** | 477 ± 2 (0.392 ± 0.010)<sup>l</sup> | - 19.5 ± 1.5 |
| **Decellularized colon extracellular matrix** | Not applicable | - 34.5 ± 1.9 |

**Supplementary Table 5.** Antibody lists for flow and imaging mass cytometry studies.

| **List of antibodies for Helios flow mass cytometry** |  |  |
| --- | --- | --- |
| Mouse Helios Abs |  |  |
| Item Name * | Clone | metal |
| CD3e 152Sm | 145-2C11 | 152Sm |
| CD4 145Nd | RM4-5 | 145Nd |
| CD8a 168Er | 53-6.7 | 168Er |
| CD11b 148Nd | M1/70 | 148Nd |
| CD11c 142Nd | N418 | 142Nd |
| CD44 171Yb | IM7 | 171Yb |
| CD45 89Y | 30-F11 | 89Y |
| CD69 143Nd | H1.2F3 | 143Nd |
| CD86 172Yb | GL1 | 172Yb |
| CD206 175Lu | C068C2 | 175Lu |
| IA/IE 209Bi | m5/114.15.2 | 209Bi |
| Ly-6G 141Pr | 1A8 | 141Pr |
| NK1.1 170Er | PK136 | 170Er |
| F4/80 146Nd | BM8 | 146Nd |
| Foxp3 158Gd | FJK-16S | 158Gd |
| PD-1 159Tb | 29F.1A12 | 159Tb |
| IFNy 165Ho | XMG1.2 | 165Ho |
| TNFa 162Dy | MP6-T22 | 162Dy |
| CCR7 164Dy | 4B12 | 164Dy |

| **List of antibodies for Hyperion imaging mass cytometry** |  |  |
| --- | --- | --- |
| Mouse Hyperion Abs |  |  |
| Item Name * | Clone | metal |
| CD4 159Tb | BLR167J | 159Tb |
| CD8 152Sm | EPR21769 | 146Nd |
| CD11b 149Sm | EPR1344 | 149Sm |
| CD11c 154Sm | 3.9 | 154Sm |
| CD45 141Pr | 30-F11 | 141Pr |
| CD69 161Dy | poly | 161Dy |
| CD44 144Nd | IM7 | 144Nd |
| CD86 156Gd | GL-1 | 156Gd |
| F4/80 153Eu | D2S9R | 153Eu |
| IA-IE 147Sm | M5/114.15.2 | 147Sm |
| Gr-1 169Tm | RB6805 | 169Tm |
| FoxP3 155Gd | FJK-16s | 155Gd |
| TNFa 162Dy | MP6-T22 | 162Dy |

